# Standard and Non-Standard Measurements of Acidity and the Bacterial Ecology of Northern Temperate Mineral Soils

**DOI:** 10.1101/2020.10.01.323014

**Authors:** Michael J. Braus, Thea Whitman

## Abstract

Databases of soil pH values today guide the decisions of land managers and the experimental designs of microbiologists and biogeochemists. Soil acidity underpins fundamental properties and functions in the soil, such as the solubilities of exchangeable ions and nutrients, or bacterial use of gradients of internal and external acidity to generate ATP and turn flagellar motors. Therefore, it is perhaps unsurprising that soil pH has emerged as the strongest predictor of soil bacterial community composition. However, the measurement of these particular values today does not address whether soil pH accurately represents the *in situ* acidity of soil microhabitats where microorganisms survive and reproduce. This study analyzes and compares soils of a large-scale natural soil pH gradient and a long-term experimental soil pH gradient for the purposes of testing new methods of measuring and interpreting soil acidity when applied to soil ecology. We extracted and prepared soil solutions using laboratory simulation of levels of carbon dioxide and soil moisture more typical of soil conditions while also miniaturizing extraction methods using a centrifuge for extractions. The simulation of in situ soil conditions resulted in significantly different estimates of soil pH. Furthermore, for soils from the long-term experimental soil pH gradient trial, the simulated soil pH values substantially improved predictions of bacterial community composition (from *R*^2^ = 0.09 to *R*^2^ = 0.16). We offer suggestions and cautions for researchers considering how to better represent soil pH as it exists *in situ*.

## Introduction

Soil pH measurements have guided land management and biogeochemical research for over a century (Libohova et al., 2012; Miller and Kissel, 2010), aiding agronomists in optimizing crop yields from soils across the spectrum of pH values. A number of methods are used to measure soil pH, notably a dilute settled soil suspension, in which a glass pH probe is immersed. The standard soil pH method has produced large databases of soil pH values, which have provided microbial ecologists one of the best existing predictors of the composition of soil bacterial communities worldwide (Bahram et al., 2018; Delgado-Baquerizo et al., 2018; Shen et al., 2013; Wakelin et al., 2016). These measurements of soil acidity hold great potential for the management of the diversity and composition of bacterial communities in target soils (Fierer and Jackson, 2006, p. 627; Lauber et al., 2009, p. 5114; Tripathi et al., 2012). In general, neutral soils (standard soil pH approaching 7) exhibit the largest diversity and abundance of bacteria, with many signals of “acidity specialists” in acidic soils as well as “alkalinity specialists” in alkaline soils (Barberán et al., 2012; Jones and Bennett, 2017; Vieira et al., 2020). However, the exact biogeochemical mechanisms underpinning the relatively strong correlation between soil pH and soil bacterial community composition remain unknown or vague, reflecting the methodological challenge of explaining the optimal soil pH of cultivated soil bacteria in classic microbiological studies (Small, 1954, p. 212) as well as more recent studies that have utilized culture-independent molecular methods (Lauber et al., 2008; Rousk, Bååth, et al., 2010; Tecon and Or, 2017).

The acidity of soil is an emergent property relying on several interacting biotic and abiotic protic reservoirs of protons (Supplemental Figure 1). Most bacteria directly depend on their microenvironments to supply the elements and molecules necessary for life, as well as to supply the Nernstian potential for protonmotive force by which cells perform oxidative phosphorylation and many other powerful cellular processes, such as powering flagella (Junge and Nelson, 2015; Lerman, 1978). However, precise theories for the responsiveness of bacteria to the acidity of soil microenvironments are diverse and contested today (Mikutta et al., 2006; Sinsabaugh et al., 2008), including abiotic factors, such as pH-mediated nutrient availability in bulk soils or the rhizosphere (Song et al., 2015; Stark et al., 2014), biotic factors, such as limitations to microbial cell densities or metabolisms (Dennis et al., 2009; Poole, 1999), or an interaction of both abiotic and biotic factors entwined. Simultaneously, because most molecular methods use solutions and substances whose chemical behaviors are highly dependent on pH and ionic strength (Barrow, 1984; Kerndorff and Schnitzer, 1980; Kirk et al., 2004, p. 171; Naidu et al., 1994; Young et al., 2014), we should also be cautious of the risk of soil acidity causing chemical biases within molecular methods themselves, such as DNA extraction and PFLA extraction (Bååth and Anderson, 2003, pp. 958–959; Frostegård et al., 2011, p. 1624; Rousk, Brookes, et al., 2010a, 2010b).

Several guides exist for the measurement of pH of concentrated solutions (e.g. Thermo Fisher Scientific Application Note 009, 2014) and invariably provide cautionary notes for the interpretation of the pH values of such solutions: “ion mobility decreases in the high ionic strength samples and the activity differs from the concentration […] High ionic strength solutions change the liquid junction potential. This may lead to bias […]. (*Measuring pH of concentrated samples*, 2014, p. 1)” However, such guidance offers little by way of insight when solving the underlying chemical problem of the highly narrow thresholds of applicability of pH to systems such as soils as they exist naturally. Solution extracts from soils of typical moisture constitute “highly concentrated solutions” owing to their greater density of ions, biomolecules, and organic matter, in addition to clays, the smallest of which being highly chemically and catalytically reactive.

Given the spatial scale at which soil microbes meaningfully perceive their environments (Vos et al., 2013), in order to effectively investigate why soil pH is such a strong determinant of bacterial community composition and to represent the dynamic acidity of soil microhabitats, accurate and precise values of *in situ* soil pH will be required (Bjerrum and Gjaldbæk, 1919, p. 4). The conditions under which standard measurements of soil pH are made in the lab likely do not correspond to conditions in the field. Juxtaposing laboratory and field conditions, we can see that, generally, the chemical properties of solutions in the controlled conditions of the laboratory (“*ex situ*”), further altered with the addition of solutions and processing of extracts, are often highly incommensurable with the same chemical properties of solutions in the field (“*in situ*”). Soil conditions in the field are undisturbed, yet they are challenging to control experimentally as they are unpredictably variable over time. This methodological challenge extends also to gases in soils. The soil atmosphere often has much higher partial pressures of carbon dioxide than surface conditions, and these partial pressures change with depth (Belnap et al., 2003; Cary and Holder, 1982; Jury and Horton, 2004, p. 215; Vernadsky, 1913) reaching levels as high as 4% to 6% at depths at or below 2 [m] and levels approximating atmospheric carbon dioxide levels (400 [ppm] or 0.04%) at depths of < 5 [cm], and the lowest extreme (0 [ppm]) is not uncommon in photosynthetic biological crusts (Oh et al., 2005). Furthermore, typical laboratory atmospheres are approximately equal to the lower atmosphere, only several hundred parts per million (depending on the human investigators present and the lab’s collection of plants) and would therefore represent the lower bound of typical soil CO_2_ concentrations. If a soil sample collected from a soil profile at 1 [m] is moved to the laboratory for measurement of acidity or other chemical characteristics, does the fact that the *in situ* CO_2_ levels may be orders of magnitude lower than the *ex situ* conditions affect our measurements of soil properties such as pH?

CO_2_ in the soil atmosphere will equilibrate with the soil solution, as described by Henry’s law, K_*H*_ = *a*_*i*_/*P*_*i*_, where, for CO_2_, K_*H*_ signifies Henry’s constant (approximately 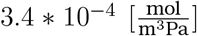 at standard temperature for carbon dioxide in water (Sander, 2015, p. 4488)), *a*_*i*_ (unitless) signifies the thermodynamic aqueous activity of CO_2_ benchmarked to the standard state, and *P*_*i*_ [Pa] signifies the partial pressure of CO_2_. As Strawn et al. (2020, pp. 90–97) explain with caution, in reference to early research (Smith et al. (1937); Whitney and Gardner (1943)) that first demonstrated the linear acidification effect of CO_2_ on soil pH of *dilute* suspensions:

> Several simplifying assumptions [are] required to solve the carbonate system equations that may not be possible or appropriate in other aqueous equilibrium problems. Additionally, the assumption that activity and concentrations are equal (ideal solution) is fine for showing trends, but activity corrections can cause significant changes in the predicted pH or concentrations of the species.

Therefore, although it would be challenging to predict the precise shift in soil pH expected from an increase in CO_2_, as Bjerrum curves relate the concentrations of carbonic acid to mono- and di-protic carbonate in dilute solutions (Andersen, 2002), elevated carbon dioxide partial pressures may not increase acidity in concentrated solutions, such as the extracts of solution from soils at typical soil water content. As noted by Šimunek and Suarez (1994) in reference to their previous two-part publication (Suarez and Šimunek, 1993; Šimunek and Suarez, 1993), “existing models also assume either a fixed pH or a fixed CO_2_, which are questionable assumptions for soils, which usually exhibit fluctuation of both of these variables.” Such “fixed” or non-varying pH and CO_2_ are obviously very uncommon in soils across textures, series, depth, and time, warranting fundamental reappraisal.

To address the overarching challenge of better representing *in situ* soil conditions in biogeochemical measurements and instrumentation, two approaches present themselves: to perform direct *in situ* measurements in the field while minimizing the perturbation of the original conditions of soil profiles (and the functionality of instruments), or to simulate the original conditions of intact soils during the analysis of soil samples that have been collected from the field and brought to the laboratory. Both of these approaches have complementary advantages and disadvantages, but both approaches are also a significant departure from traditional methods described in standard methodological references (Jacob et al., 2002, pp. 1481–1509). While most soil scientists rarely measure soil solution pH in the field, due to the numerous challenges of doing so, scientists in other fields are acutely aware of the value of *in situ* measurements or maintaining *in situ* conditions, as exemplified by the works of Sasowsky and Dalton (2005) on the importance of such measurements of water chemistry in caves, Parfitt et al. (1995) on the chemistry of aluminum in suspensions of orchard soils, and Matthiesen (2004) in archaeological excavations.

The present study expands upon the foundational soil acidity experiments performed by Whitney and Gardner (1943), with application to soil bacterial ecology. Additionally, beyond the improvement of the fundamental understanding of bacterial ecology of soils, the paradigm of “soil pH” itself is explored in terms of metrological interpretation in parallel with standard and non-standard soil acidity measurement protocols (acidimetry). This study presents a multifactorial chemical and microbial study across both natural and experimental soil pH gradients in temperate mineral soils in Wisconsin, USA. We assess the limitations of soil pH measurements using a non-standard methodology: extraction of soil solution at moisture levels approximating field capacity and drier, miniaturization of the resulting analyte to allow for high-throughput pH measurement, simulation of soil conditions during pH measurement, and exponentiation of pH values to hydrogen ion activity 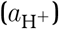. Non-standard soil pH values are then used to predict soil microbial community composition across said experimental and natural pH gradients in the Wisconsin region of the United States. We hypothesized that these protocols would improve correlations with both chemical properties of soils as well as microbial community features, due to the improved representation of *in situ* soil conditions, with the ultimate goal of better informing the mechanisms by which the acidity of soil microhabitats influences soil microorganisms.

## Methods

### Standard and Non-Standard Soil pH Values

Our objective was to determine whether standard soil pH measurements or nonstandard soil pH measurements (i.e., soil pH values under conditions simulating *in situ* soil conditions of moisture and carbon dioxide levels) were better predictors of bacterial community composition across soil pH gradients. For the purposes of this study, we define “standard soil pH” as the pH value measured at ambient carbon dioxide (approximately 0.04%) and a ratio of solution:soil of 1 : 1 (Thomas, 1996, pp. 487–488), where the solution may vary from deionized water (pH_W_) to a dilute (0.01 [mol/L]) electrolyte solution (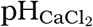 or pH_KCl_). For the comparison to standard soil pH in this study, “simulated soil pH” is defined as the multifactorial set of pH values measured at ambient and elevated carbon dioxide (2.2%(±0.05)) and a range of 1 : 2 to 1 : 4 solution:soil ratios. All solutions added to soils in this study were the dilute electrolyte 0.01 [mol/L] KCl. For each sample, we applied a miniaturized, centrifuge-based soil solution extraction method, manipulating solution:soil ratios and atmospheric CO_2_ levels during measurement using a glass microprobe to measure pH (specific details follow).

### Site Descriptions and Sample Collection

In order to investigate the effects of these methods on soils with similar underlying mineralogy, we collected and analyzed soils from a 25-year soil pH manipulation trial at the University of Wisconsin-Madison Spooner Agricultural Research Station (Spooner, WI; details of manipulation below). In order to investigate the effects of these methods on a wide range of soil types, we applied these methods to soil spanning a natural soil pH gradient of nine University of Wisconsin-Madison agricultural research stations from across the state. Where noted, “Topsoil” signifies any combination of A horizons, and “Subsoil” signifies all beneath the A horizon to the depth specified.

The pH manipulation trial at the Spooner Agricultural Research Station began in 1994 (“Long-term pH Trial”). The study soil is of the series Mahtomedi, consisting of very deep, excessively drained, rapidly permeable soils formed in sandy outwash of the Late Wisconsinan Age on glacial moraines and outwash plains. Corn, soy, and alfalfa have been grown at the site. Four replicates of 22 [m] wide by 220 [m] long field plots have been maintained at target soil pH values of 4.7, 5.2, 5.7, 6.2, and 6.7, through annual additions of pell lime or sulfur after annual soil tests (personal correspondence with Superintendent Phil Holman). Samples were collected on November 3, 2017. Three 1-inch diameter cores to 20 [cm] depth were randomly sampled at locations determined by a random number generator using the length of the long rectangular plots, avoiding the plot edges by 5 [ft].

The second set of sites (“Wisconsin Soils”) were selected using legacy chemical and physical data for University of Wisconsin Agricultural Research Stations from Web Soil Survey, retrieved on August 7th, 2018. From the database’s graphical user interface, a depth of 0 [cm] to 50 [cm] was selected for the following parameters: calcium carbonate, cation exchange capacity at pH 7 (CEC-7), electrical conductivity (EC), gypsum, soil pH, sodium adsorption ratio, available water capacity and supply, bulk density at 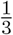 bar, liquid limit, percent organic matter, percent clay, percent sand, percent silt, and saturated hydraulic conductivity (*K*_*sat*_), parent material, and representative slope. These features were used to select a wide variety of characteristics, namely the widest breadth of textural classes, organic matter content, and soil pH values. The following research stations were selected, listing ID letter and soil pH values according to Web Soil Survey listed in parentheses: Kemp (K, 5.40), Rhinelander (R, 5.50), Marshfield (M, 5.65), Spooner (S or Sp, 5.80), Hancock (H, 6.20), Arlington (A, 6.50), Lancaster (L, 6.60), West-Madison (W, 6.70), and Peninsular (P, 7.20). Supplemental Figure 2 shows a map of the locations of these sites across the soil pH gradient in Wisconsin, while Supplemental Table 1 lists the latitude and longitude of each site (Kartesz, 2015).

The Wisconsin soils were collected from each of the two or three most common soil series of each agricultural research station listed above, between August and September, 2018. At each site, a soil pit was dug to > 50 [cm] depth and, after excavation, several kilograms of soil were gathered from each horizon evenly spanning the upper to the lower boundary. Horizon boundaries were easily visible, and photos of all soil profiles can be found in the Supplemental Materials. Soil samples were placed in sterile bags and transported within 24 hours of collection to the Department of Soil Science at the University of Wisconsin-Madison and placed in a refrigerator (4 [°C]). Within two days of arrival, each sample was homogenized, subsampled, and stored at −80°C.

### Soil Chemical Analyses

The Spooner Agricultural Research Station performed chemical analyses for the long-term experimental soil pH plots in 2017: organic matter was 2.15% (±0.24), phosphorus level was 33 (±6) [ppm], and potassium level was 93 (±25) [ppm] (personal correspondence with Superintendent Phil Holman). All samples of the Wisconsin set were homogenized, subsampled, and submitted to the University of Wisconsin Soil and Forage Laboratory where the samples were dried and sieved to conduct the following analyses: Routine Tests (pH using 1:1 water, P using Bray No 1 extraction test, K also using Bray No 1 extraction test, and OM using loss on ignition), Cation Exchange Capacity (summation, including calcium and magnesium), acidity extracted using ammonium acetate, and total nitrogen and organic carbon (dry combustion) (specific protocols in Burt and Staff (2014)).

### Soil Solution Extraction

The “suspension effect” has long been observed (Gorham, 1960; Jenny et al., 1950; Oman et al., 2007; Ponnamperuma et al., 1966), and describes the apparent decrease in pH when a pH probe is moved between the supernatant and sediment of a settled suspension, although the precise explanation for the problem is somewhat unresolved (Feldman, 1956; Fornasier et al., 2018). Sacchi et al. (2001) have recommended preparing fresh samples using the centrifugation method of extracting solutions from clay-water systems, pertaining to most unsaturated soils, with a risk of incomplete water extraction at extreme dry conditions. In order to minimize the “suspension effect”, we reduced the density of soil particles from solution extracts via centrifugation, and measured the supernatant rather than the sediment.

Soil solution was extracted as follows, informed by Gillman (1976) and Wolt (1994, pp. 95–120). Empty tubes were labeled and weighed, and masses were recorded. Packed fresh (not dried) soil was added to fill 1.0 [mL] to 1.3 [mL] of the tube, and the exact mass added was recorded. The soil mass was used to estimate the volume of 0.01 [M] KCl solution (specific mass approximately equal to water, or 1.0 [g/mL]) required to reach the target solution:soil ratio (1 : 1, 1 : 2, 1 : 3, or 1 : 4). The addition of a weak electrolyte such as 0.01 [M] KCl minimizes the liquid junction potential of glass probe pH acidimetry (Bates, 1973, pp. 31–58; Kadis and Leito, 2010; Libohova et al., 2014; MacInnes, 1915). This solution produces highly dilute spectator ions without acid-base reactivity that cannot increase ionic strength past the threshold beyond which pH is applicable while minimizing liquid junction potentials. Tubes were then vortexed until well-mixed and let rest 40 minutes to 1 hour. Tubes were centrifuged for 60 seconds at 8000[RPM], which causes a relative centrifugal force (RCF = RPM^2^ × 1.118 × 10^−5^ × rotational radius) equal to 7, 155 g force. 100[*μ*L] of supernatant was pipetted into a 0.5 [mL] tube for measurement. All aliquots were prepared and then frozen at −20[°*C*] for later thawing and pH measurement. The original soil remaining in the 1.5 [mL] tubes after centrifugation and supernatant extraction was then dried and massed. These dry soil mass values enabled the calculation of the starting gravimetric water content, from which the exact solution:soil ratios were calculated for subsequent analyses.

### Simulation of Soil Atmospheric Carbon Dioxide

For samples measured under elevated CO_2_, we used a vinyl anaerobic airlock chamber (Coy Laboratory Products, Inc., Grass Lake, Michigan, see Supplemental Figures 3 and 4) to maintain an atmosphere of 2.2%(±0.05) CO_2_. The elevated CO_2_ level decreased the pH of 1.0[mL] of 0.01[M] CaCl_2_, which was used as a standard throughout the experiment, from 7.0(±0.05) in normal laboratory conditions to 6.0(±0.05). CO_2_ was produced in the chamber through the initial reaction of 100[g] of NaHCO_3_ with excess 5% acetic acid, after which CO_2_ levels were adjusted to target levels with a combination of venting and additional reactions. The chamber air was mixed with a small fan and CO_2_ was monitored with a USB CO_2_ Probe Data Logger (CO_2_Meter.com, K-30 Probe, CM-0040) with a measurement range of 0% (0 [ppm]) to 30% (300, 000 [ppm]), with an error not exceeding 5% of the quantity measured and logged using the GasLab software (v. 2.2.1.36). Samples measured under ambient CO_2_ were measured in the same chamber fully open and vented to the laboratory space.

### Measurement of pH with a Microprobe

pH was measured using an InLab Micro pH glass microelectrode (Mettler-Toledo; Material No. 51343160; further details on probe can be found in Supplemental Materials). To monitor the quality of measurements throughout the analysis at elevated CO_2_, the pH of identical volumes of several controls were taken alongside the soil extract, including 100 [*μ*L] each of 0.01[M] CaCl_2_, 5% (0.833[M]) acetic acid, 0.01 KCl, and deionized water. The 0.01 KCl solution was measured every 50 soil pH measurements to detect probe drift. These control values deviated < 0.15 pH units during each series of measurements across the entire experiment. Exponentiation of the soil pH values did not require further measurements but rather calculated activity of hydrogen ions 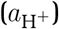, which adopts the units of moles per liter to represent effective concentration when the activity coefficient of hydrogen ions (i.e., hydronium and related cationic species of solvated protons) is 1.0 (de Levie, 2014).

### Statistical Analyses for Chemical Properties

To compare the non-standard pH values with standard values, we fit linear regression models to determine their relatinships. To determine which other soil chemical properties were the most strongly associated with soil pH as measured by the standard and non-standard methods, linear models correlating soil chemical measurements and all values of pH were analyzed using a Bayesian information criterion (BIC) approach (Kass and Wasserman, 1995). The calculations were performed in R (Team, 2018; Wickham, 2009) using the **regsubsets**function from the R package `leaps` (Lumley and Miller, 2020). Interpretation of the results involved assessing which factors, when added to the model, produce the most negative BIC, where more negative BIC values indicate better models when certaint factors are incorporated and others excluded. The collection of models with the most negative BIC values in the “BIC dropoff” region offer an assortment of models that best predict the factor of interest−in our case, pH. We calculated models and their associated BIC values using the soil chemical analyses as predictors for each of the four sets of soil pH values generated for the extremes of this study’s multifactorial: high and low CO_2_ and the highest and lowest soil solution content (1 : 1 and 1 : 4 solution-to-soil ratio by mass).

### Soil DNA Extraction and Bacterial Community Sequencing

Total genomic DNA was extracted from frozen soils using the PowerLyzer PowerSoil DNA Isolation Kit (Catalog No. 12888, Qiagen, Germantown, MD, USA). All DNA was stored at or below −20[°*C*] from the date of extraction throughout stages of sequencing. Because soil pH can potentially interact with the chemicals used for extracting DNA, we also investigated the predictive value of the pH of solutions along two steps of the DNA extraction protocol (see Supplemental Figure 9 and Supplemental Note 2). 16S rRNA genes were amplified from extracted DNA using polymerase chain reaction (PCR), with three replicate reactions per sample. Variable region V4 of the 16S rRNA gene was targeted using forward primer 515F and reverse primer 806R with modification by Walters et al. (2016), which increased degeneracy of bases that have caused detection bias among some bacterial clades. Primers also had barcodes and Illumina sequencing adapters added, following Kozich et al. (2013) (all primers in Supplemental Table 2). The following reagents were added to each PCR reaction: (1) 12.5[*μ*L] Q5 Hot Start High-Fidelity 2X Master mix (New England BioLabs INC., Ipswich, MA), (2) 1.25[*μ*L] 515f forward primer (10[mM]), (3) 1.25[*μ*L] 806r reverse primer (10[mM]), (4) 1[*μ*L] DNA extract, and (5) 7.75[*μ*L] PCR-grade water. The plate was sealed, gently vortexed, and briefly centrifuged to ensure all liquids were well mixed. The plate was then run on an Eppendorf Mastercycler nexus gradient thermal cycler (Hamburg, Germany) using the following parameters for 30 cycles: 98[°C] for 2 minutes + (98[°C] for 30 seconds + 58[°C] for 15 seconds + 72[°C] for 10 seconds) × (30 + 72) [°C] for 2 minutes and 4[°C] hold.

Successful amplification was verified via gel electrophoresis. To purify amplicons and normalize PCR products, we used a SequalPrep Normalization Plate Kit (Invitrogen Corporation, Thermo Fisher Scientific, Waltham, MA, USA). The PCR triplicates for each sample were pooled and normalized according to manufacturer’s instructions. The Wizard SV Gel and PCR Clean-Up System A9282 (Promega, Madison, WI) was used to extract and purify the combined PCR product library according to manufacturer’s instructions except for the following two deviations: (1) the SV Minicolumn incubation and centrifugation (steps 5.A.2-5.A.3) steps were repeated twice for each sample, and (2) nuclease-free water application was divided into and increments with the incubation step and centrifuge step after each addition (step 5.A.6). DNA was concentrated using a SpeedVac Vacuum Concentrator System (Thermo Fisher Scientific, Waltham, MA, USA) before and after using the Wizard SV Gel and PCR Clean-Up to meet the sequencing requirements of 15[ng*/μ*L]. The final library was sequenced at the University of Wisconsin-Madison Biotechnology Center on a Illumina MiSeq Sequencer using 2 × 250[bp] paired-end reads.

### Microbial Community Analyses

Sequencing generated 1.3M reads, with a mean of 104, 655 reads per sample (minimum 48, 207, maximum 257, 394 reads per sample). We quality-filtered and trimmed (truncation length 235 bp for forward and 144 bp for reverse reads, left trim of 5 bp for forward and reverse reads with other default settings), learned errors (using all sequences), dereplicated, determined operational taxonomic units (OTUs) (default settings), and removed chimeras using dada2 (Callahan et al., 2016) as implemented in R, and run on the UW-Madison Center for High-Throughput Computing cluster. This resulted in a final mean of 53, 777 reads per sample (minimum 18, 610, maximum 152, 682 reads per sample). All reads have been deposited at the National Center for Biotechnology Information Short Reads Archive under BioProject ID PRJNA643927.

We analyzed bacterial communities using the R packages `phyloseq` (McMurdie and Holmes, 2013) and vegan (Dixon, 2003). OTUs were filtered to remove mitochondria and chloroplast sequences and were normalized by relative abundance for each sample. We assessed the influence of pH measurement technique on community composition using permutational multivariate analysis of variance (PERMANOVA) (Anderson, 2014) with Bray-Curtis dissimilarities (Bray and Curtis, 1957) and illustrated these differences in community composition using non-metric multidimensional scaling (NMDS) plots (Agarwal et al., 2007). Code for all analyses can be found at https://github.com/michaeljbraus/usda-wisconsin-soil-ph.

## Results

### Non-Standard Soil pH Values at Four Levels of Soil Moisture

Soils from across Wisconsin’s natural soil pH gradient spanned a wide range (4.4 to 7.8) of standard soil pH values (Figure 1). In the ambient CO_2_ atmosphere, decreasing solution:soil ratios resulted in changes in measured pH spanning decreases of more than 1.0 to increases of more than 1.0, with 18% of measured values differing by more than 0.5 units from their standard soil pH measurements (Figure 2). Among the soils from the long-term pH manipulation trial, the more alkaline soil pH values > 6.5 tended to increase, by approximately 0.2 and up to 1.0 when soil moisture was lowered, whereas the soils of pH < 6.5 changed little with decreasing soil moisture and had somewhat lower variability (Figure 2). These trends were similar in the cross-Wisconsin dataset, where soils with pH exceeding approximately 6.0 tended to increase in pH with decreasing water soil moisture, whereas soils of lower pH tended to change less or to decrease. Across both datasets, and for all soils, pH tended to decrease among solution extracts of 3 : 1 soil:solution ratio in comparison to a 1 : 1 ratio, and then increase again at the 4 : 1 soil:solution ratio (Figure 2).

**Figure 1.**
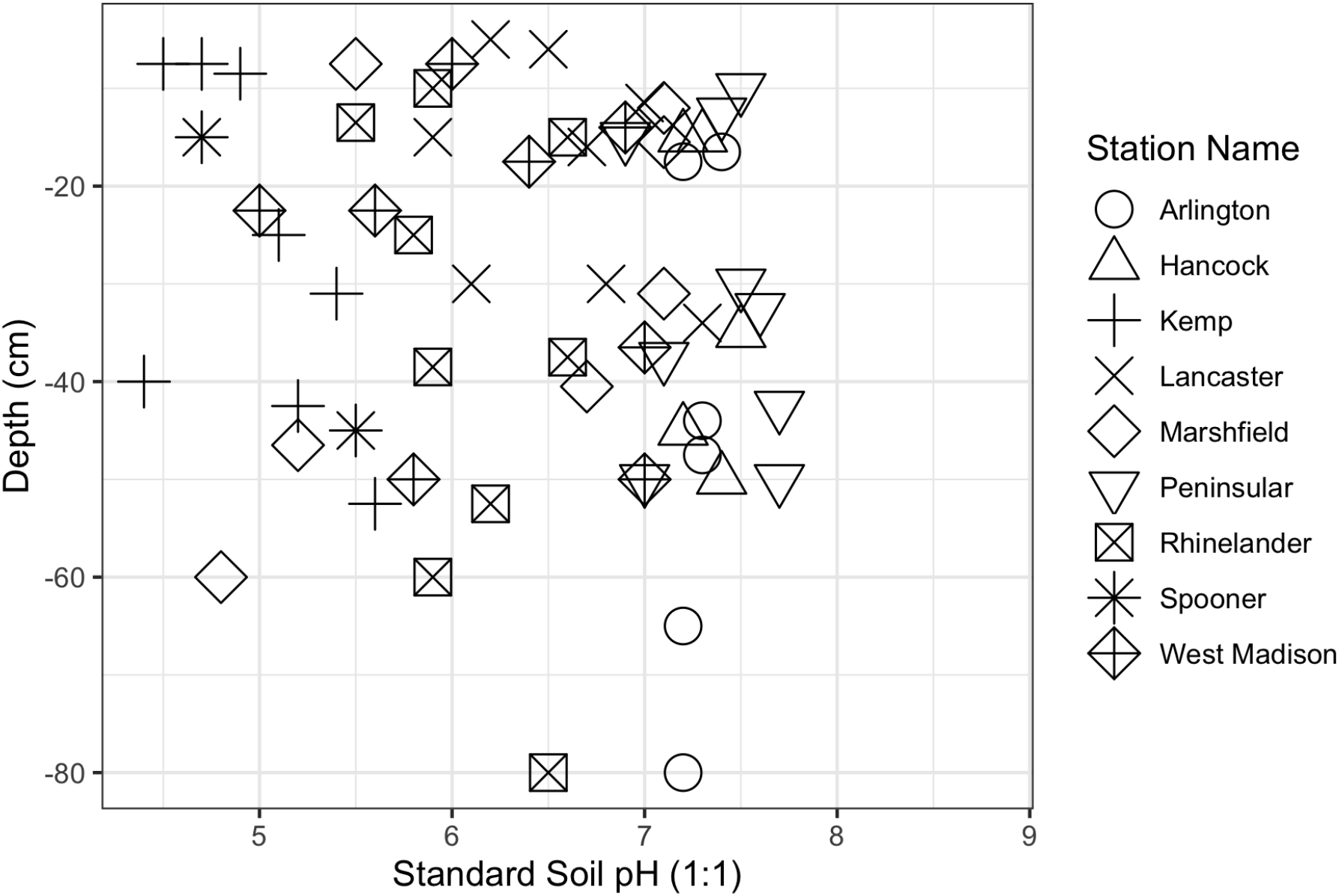
Standard soil pH values for all samples as a function of depth from soil samples from agricultural field stations across Wisconsin. See also Supplemental Figure 2 depicting the relative locations in Wisconsin of these stations.

**Figure 2.**
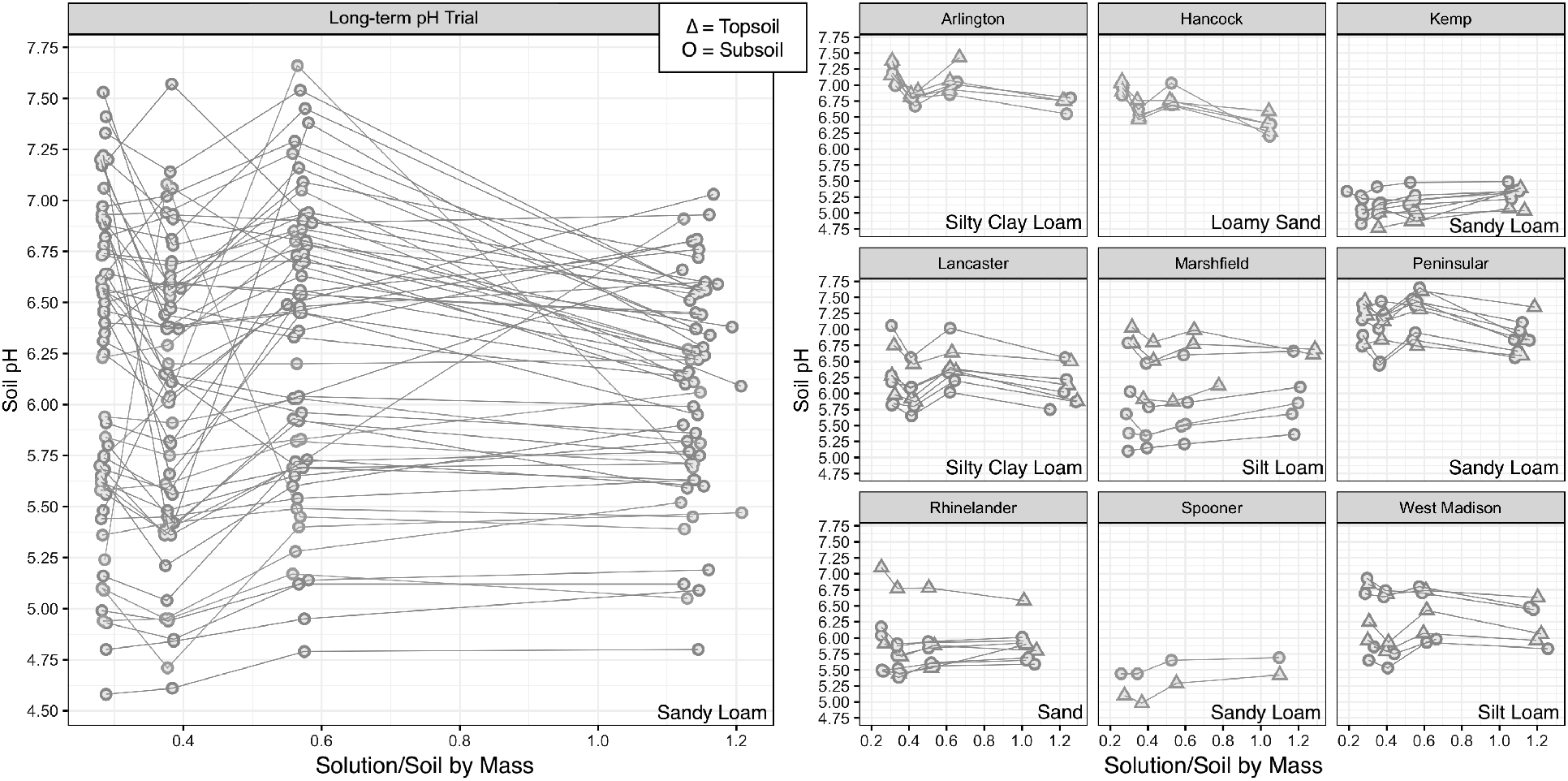
Soil pH as a function of solution-to-soil ratio for (A) soils from a long-term pH manipulation trial in Spooner, WI and (B) soils from across Wisconsin’s natural soil pH gradient. Each point represents a single pH measurement. Triangles indicate topsoil samples, while circles indicate subsoil samples for the Wisconsin dataset. Topsoil and subsoil are not distinguished in the long-term pH manipulation trial dataset. Points from the same soil sample are joined by straight lines for ease of comparison. Soil texture is indicated in the bottom right quadrant of each sub-plot. Note that exact solution:soil ratios are plotted, hence the small variation in the x-axis for a given moisture treatment.

Under the 2.2%(±0.05) CO_2_ atmosphere, all soils in the cross-Wisconsin dataset tended to increase in measured pH with decreasing solution:soil ratios, but the same general trend of higher pH soils being more affected by decreasing solution:soil ratios persisted (slopes 1.18 − 1.22). In the long-term soil pH manipulation trial, pH of most samples tended to increase with decreased moisture contents, and the higher pH samples again had somewhat greater variability (Table 1 and Supplemental Figures 5-8).

**Table 1.** Linear regressions of soil pH and soil activity 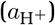 relating these values at ambient laboratory CO_2_ (0.04%) to values at a typical soil atmospheric CO_2_ (2.2%(±0.05)) as a result of chemical analysis of the Wisconsin Soils set from across a natural soil pH gradient and the Long-term pH Trial set of samples from an experimental soil pH gradient.

| Soil Set | Acidity Metric | Solution:Soil Ratio | Intercept | Slope | R-squared |
| --- | --- | --- | --- | --- | --- |
| Wisconsin Soils | pH | 1-to-1 | -0.442 | 1.049 | 0.976 |
| Wisconsin Soils | pH | 1-to-2 | -0.234 | 1.065 | 0.927 |
| Wisconsin Soils | pH | 1-to-3 | 0.099 | 1.033 | 0.967 |
| Wisconsin Soils | pH | 1-to-4 | 1.001 | 0.862 | 0.959 |
| Wisconsin Soils | a(H<sup>+</sup>) | 1-to-1 | 0.000 | 1.687 | 0.941 |
| Wisconsin Soils | a(H<sup>+</sup>) | 1-to-2 | 0.000 | 0.976 | 0.897 |
| Wisconsin Soils | a(H<sup>+</sup>) | 1-to-3 | 0.000 | 0.711 | 0.959 |
| Wisconsin Soils | a(H<sup>+</sup>) | 1-to-4 | 0.000 | 0.580 | 0.981 |
| Long-term pH Trial | pH | 1-to-1 | 0.218 | 0.965 | 0.843 |
| Long-term pH Trial | pH | 1-to-2 | 0.639 | 0.910 | 0.928 |
| Long-term pH Trial | pH | 1-to-3 | 1.190 | 0.842 | 0.909 |
| Long-term pH Trial | pH | 1-to-4 | 1.600 | 0.748 | 0.867 |
| Long-term pH Trial | a(H<sup>+</sup>) | 1-to-1 | 0.000 | 1.075 | 0.963 |
| Long-term pH Trial | a(H<sup>+</sup>) | 1-to-2 | 0.000 | 0.707 | 0.978 |
| Long-term pH Trial | a(H<sup>+</sup>) | 1-to-3 | 0.000 | 0.507 | 0.937 |
| Long-term pH Trial | a(H<sup>+</sup>) | 1-to-4 | 0.000 | 0.613 | 0.976 |

### Non-Standard Soil pH Values at Ambient and High CO_2_

Soil pH values were also affected by the level of carbon dioxide during measurement. In the long-term pH manipulation trial soils, increasing CO_2_ did not markedly change measured pH values for solution:soil ratios of 1 : 1 to 1 : 3. However, at solution:soil ratios of 1 : 4, increasing CO_2_ decreased measured pH values in the higher pH samples (pH > 6.5, approximately) (Figure 3A and 3C). In the cross-Wisconsin dataset, only solution extracts prepared according to the standard (1 : 1) ratio exhibited the expected trend of acidification at elevated carbon dioxide, with measured pH values decreasing by as much as 0.6, while in samples with lower solution:soil ratios, increasing CO_2_ increased measured pH slightly (Figure 3B and 3D).

**Figure 3.**
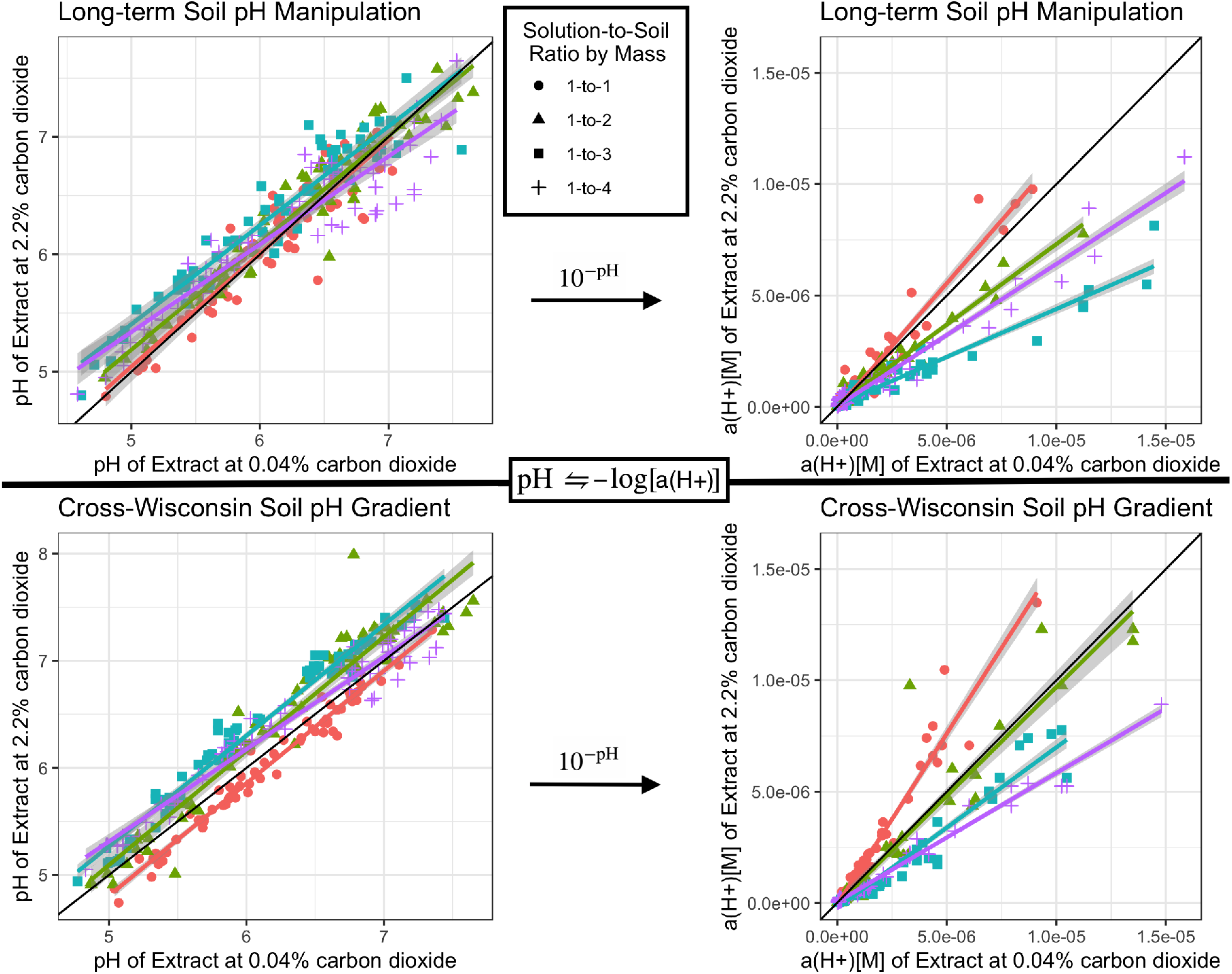
Soil pH and hydrogen ion activity 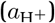 values, which are interchangeable according to the definition of pH scale as the negalitive log of hydrogen ion activity, measured at ambient or low (0.04%) and high (2.2%(±0.05)) carbon dioxide levels and soil water content at four levels from the natural cross-Wisconsin soil acidity gradient and long-term soil pH manipulation gradients. Grey regions surrounding linear regression lines are standard error, and the solid black line signifies *y* = *x*. Points are labelled by color and shape to signify solution:soil ratio, where red circles = 1 : 1, green triangles = 1 : 2, blue squares = 1 : 3, and purple crosses = 1 : 4.

### Correlations of Soil Properties with pH Measurements

For the cross-Wisconsin dataset, the factors significantly correlated with standard and simulated soil pH values fall into the broad categories of textural (sand, silt, and clay content), chemical (SOM, C, N, P, K), and exchangeable (CEC and exchangeable acidity). The most consistently influential correlates for soil pH values were the exchangeable factors and the Bray-extracted phosphorus (Figure 4). The decrease of water content from a solution:soil ratio of 1 : 1 to 1 : 4 generally caused the influence of textural factors to decrease and chemical factors to increase. Calcium was not influential in any model, and changing CO_2_ levels had little influence on the model results.

**Figure 4.**
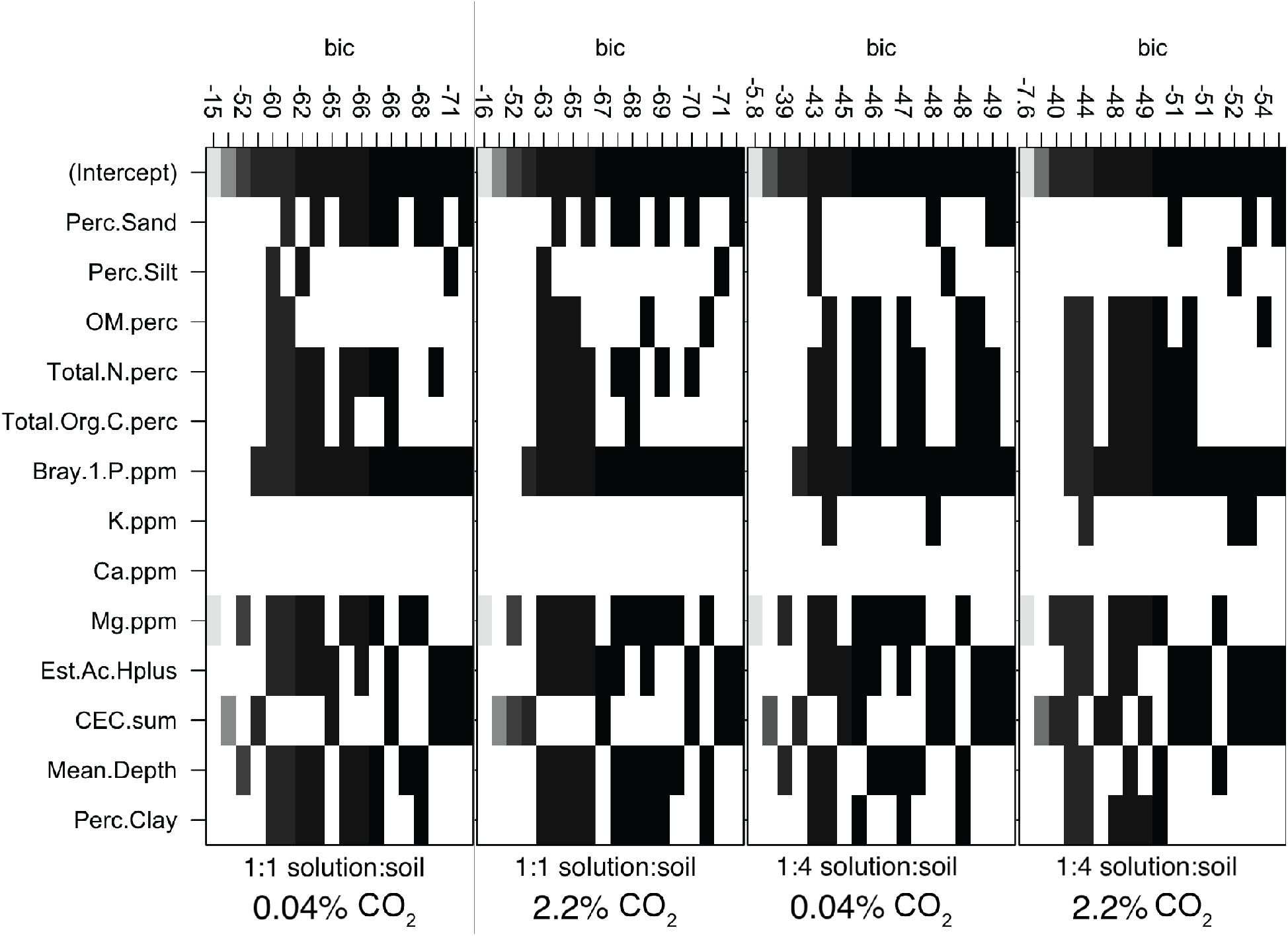
Bayesian information criterion (BIC) plot for soil properties as possible correlates of soil pH as determined by a ratio of solution:soil ratio of 1 : 1 compared to 1 : 4 and a soil atmosphere with approximately 0.04% compared to 2.2%(±0.05) carbon dioxide. Vertical axes are discrete and not continuous, where each value represents the ranked BIC value of the model using the input factors indicated by blocks. Shading of blocks indicates the degree to which a proposed model can be considered relevant, where the darker squares represent good selections to include in a chosen model.

### Microbial Communities

In both datasets, out of all tested soil properties (pH, total organic C, total N, percent sand, percent silt, CEC, K, Mg, Ca, Bray P, and soil depth), pH was the best predictor, with stronger effects for the long-term pH manipulation trial (PERMANOVA, *R*^2^ = 0.1341, *p* = 0.001; Figure 5), than the cross-Wisconsin soils (PERMANOVA, *R*^2^ = 0.0864, p= 0.001; also Figure 5). Other factors besides soil pH were also correlated with the microbial community dissimilarities found among the Wisconsin soils, but to lesser degrees (*R*^2^ < 0.086). In a multivariate model, every soil property that was included added explanatory power for community composition (PERMANOVA, *p* = 0.001-0.03, 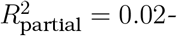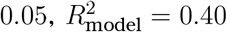).

**Figure 5.**
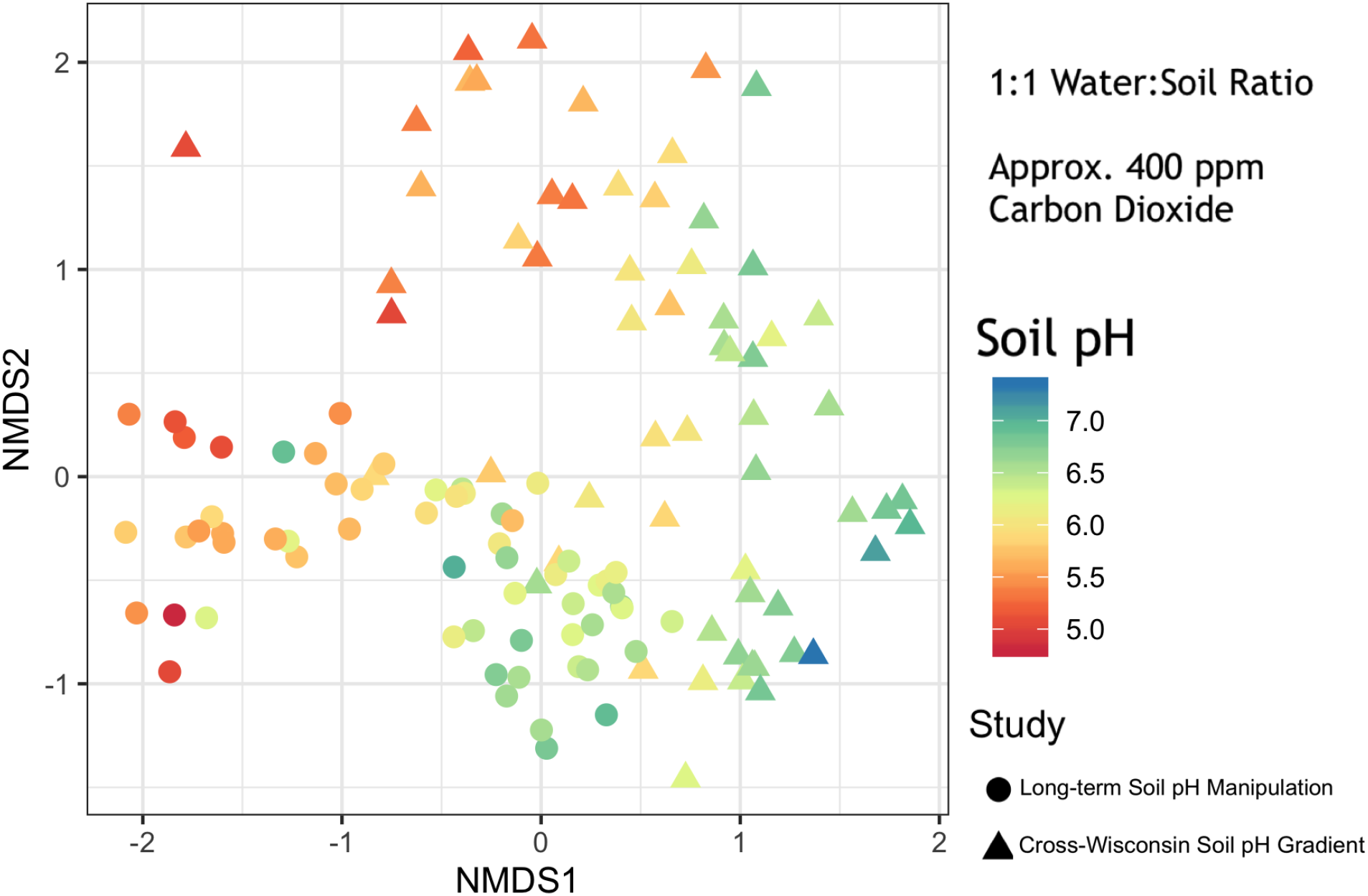
Non-metric multidimensional scaling plots of Bray-Curtis dissimilarities for soil bacterial communities from 16S amplicon analysis of two sets of samples: a long-term soil pH manipulation trial and a cross-Wisconsin soil dataset (*k* = 3, stress = 0.109). Points are coloured by standard soil pH (1 : 1 solution:soil and atmospheric CO_2_).

Low-moisture measurements of soil pH were better predictors of microbial community composition than standard soil pH in the long-term soil pH manipulation trial, explaining as much as 16% of bacterial community dissimilarity (Figure 6). However, low-moisture measurements did not substantially improve the predictive value for the Wisconsin soils (*R*^2^ = 0.086 ± 0.002 throughout). Carbon dioxide levels showed little influence on the predictive power of any measurement of soil acidity for both datasets. Activity measurements 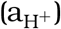 were poorer predictors of microbial community composition than pH (Figure 6). Our findings also suggested that biases in DNA extraction solutions did not explain the effects of pH on bacterial community composition, although the pH of the extraction solution was significantly and negatively correlated with soil calcium content (*p* < 0.001, *R*^2^ = 0.41; Supplementary Note 2)

**Figure 6.**
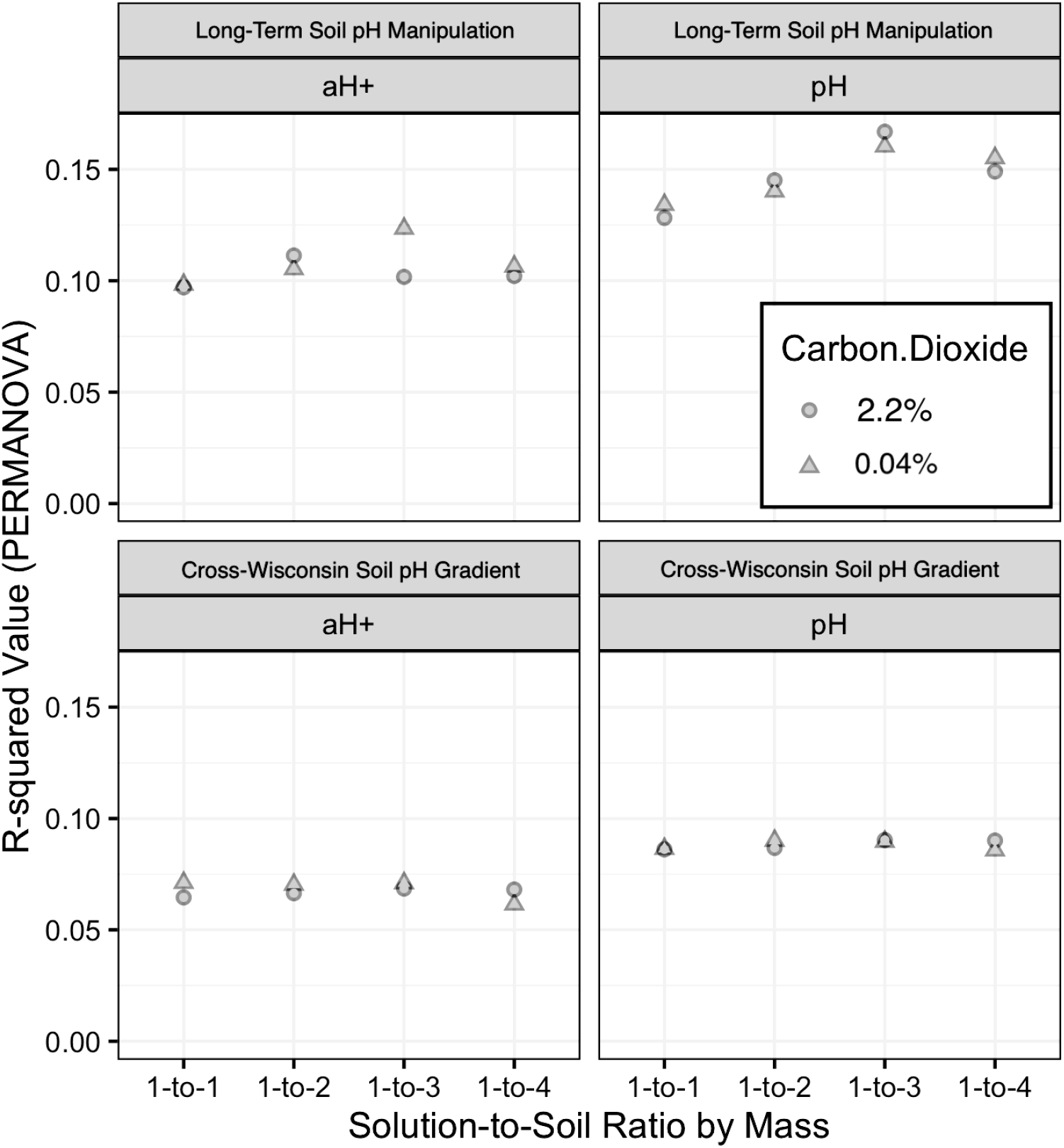
R-squared (*R*^2^) values yielded from a PERMANOVA analysis of all soil pH values and activity values 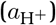 as factors predicting bacterial community composition, determined by 16S amplicons for the cross-Wisconsin soils set and long-term pH manipulation soils set analyzed in this investigation.

## Discussion

### Relation of Non-Standard Soil pH Values to Standard Soil pH

Standard soil pH measurements have underpinned fundamental advances in agronomy, allowing land managers to optimize the acidity of soils to support the production of diverse and abundant crops. However, while standardized methods allow for strong replicability across locations and time, these *ex situ* measurements of first dried and then saturated soil slurries were never designed to attempt to mimic accurately *in situ* soil conditions. The combined methods of extraction via centrifugation and miniaturization of analyte investigated in this study were designed to allow us to more accurately characterize the *in situ* acidity of soil microhabitats. However, our finding that standard soil pH values did not consistently correspond to simulated soil pH values as solution:soil ratios decreased (Figure 2) presents a confounding aspect of soil biogeochemistry. This finding echos the works of Bjerrum and Gjaldbæk (1919), review by Jackson (1958, p. 43) (Supplemental Figure 10), resurfacing of the issue by Kilian (1961) and Mubarak and Olsen (1976, p. 882), and revisited by a number of other more recent studies, such as the work of Elberling and Matthiesen (2007). We will discuss here two observed patterns when comparing standard and non-standard soil pH values.

First, the “zig-zagging” behaviour of measured pH as soil moisture was lowered from a slurry (1 : 1 solution:soil by mass) to a more typical soil moisture content (1 : 4 solution:soil by mass) (Figure 2) may be the result of a “chemical competition” between the various acidic and basic buffers present in soil solutions. Multiprotic acids, multiprotic bases, and the liquid junction potential together may compete for dominance in their influence on the solutions’ acidities, causing the oscillation of pH values observed as soil solution extracts grew increasingly concentrated. Because different chemical compounds all interact with each other to determine their respective equilibrium concentrations, effectively concentrating the soil solution by as much as 4× could certainly have different effects on chemical equilibria (and corresponding emergent pH values) as solution:soil ratios decrease. For example, an inital increase in carbonate dissolution could have caused the pH to rise, but then the effect could become overwhelmed as the strength of the acidity of the soil organic matter in solution was further concentrated. The BIC models support this changing-factor rationale: models predicting pH values for low solution:soil ratios were less influenced by the textural properties of the soils and more influenced by the chemical properties of the soils, as compared to models for standard soil pH (Figure 4).

Second, we expected that increasing CO_2_ would dissolve as carbonic acid and acidify the solution in all cases, as was outlined by Strawn et al. (2020, pp. 90–97). This effect was observed in the standard soil pH measurements only, and all concentrated soil solution extracts (i.e. typical soil moisture content) exhibited the opposite trend. Considering only the standard soil pH values of this study, Mubarak and Olsen (1976, p. 882) showed a comparable trend where, using standard 1 : 1 soil slurries, “the loss of CO_2_ from the soil samples caused the pH to increase from 0-0.3 pH units. In other words, an error of as much as +0.3 pH units can occur simply by allowing loss of CO_2_ from the sample by equilibration with the atmosphere.” Kaupenjohann and David (1996) found that degassing carbon dioxide raised soil pH values by as much as +0.3 as well, but these experiments were conducted using contained bottles, which may not correspond to an experiment testing soils exposed to the larger atmospher or chamber with carbon dioxide. In another study by Dahlgren et al. (1997), degassing carbon dioxide did not significantly affect soil pH, but a large decrease in ionic strength was observed, owing to the loss of of the 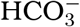 anion. Using a similar methodology to this study, the authors concluded that “failure to recognize this artifact could seriously affect the interpretation of data resulting from collection and analysis of soil solutions extracted by centrifugation.” Thus, if one wants to gain an accurate estimate of soil pH as it exists in the field, one must maintain or otherwise replicate the atmospheric conditions under which soil microhabitats existed *in situ*.

In contrast to our expectations, at the lower solution:soil ratios, increasing CO_2_ in the atmosphere during pH measurements had minimal effects or even alkalifying effects, instead of the consistent acidifying effect as predicted. This may be a relatively minor effect - for the lines of best fit relating standard pH to non-standard pH in the cross-Wisconsin dataset, the shifts in intercept were not large for 1 : 2 or 1 : 3 solution:soil ratios (Figure 3), and the slopes, although different for each ratio, are still very close to 1 in terms of effect size (1.06 and 1.03). While slope and intercept were both significant for the lowest soil:solution ratio (1 : 4), this may be largely driven by the clustering of points at the higher pH levels that responded as would be expected−i.e., decreasing under high CO_2_. For the long-term soil pH trial dataset, lines of best fit changed similarly to the cross-Wisconsin dataset with decreasing solution:soil ratios, suggesting that the small shifts in pH for lower solution:soil ratios with increased CO2 represent complex and unpredictable behaviour of solutions of high ionic strength (> 0.1[M]).

Our observations of the effects of solution concentration on pH in this study were generally consistent with the conclusions of Chapman et al. (1941, p. 200)−namely, that soils having a moisture content above approximately 30% gravimetric soil water content exhibit a more consistent soil pH value, approaching neutral with further dilution, whereas in soils of lower moisture content (i.e., most soils in the environment), these pH values will diverge in a variable magnitude and sign. Highly diluted solutions, such as those in which we typically measure soil pH, resemble the highly dilute solutions to which aqueous models apply well, but we must recognize that soils at typical soil moisture levels are considered highly concentrated solutions, and thus intractably violate the “dilute solution assumption” required for most models of aqueous chemistry. Without meeting this key assumption, we cannot accurately apply−without extreme caution−most aqueous chemical models, such as the Debye-Hückel theory (Debye and Hückel, 1923; Ferguson and Vogel, 1927), Sørensen’s acidity function named “pH” (MacInnes, 1948; Sørensen, 1909), and mean ionic activity itself (Lewis and Randall, 1921). Drained mineral soils and the sediments of brackish regions, such as the coasts of all oceans and saline seas, therefore have an effective ionic strength surpassing that which permit standard applications of pH measurements altogether. Only soils that are naturally highly saturated and would not require the addition of solution to produce a dilute supernatant for analysis would enable commensurability of soil pH to *in situ* pH, and even these soils risk substantial shifts in pH upon extraction due to degassing of CO_2_ and even other gasses, such as NH_3_ (Elberling and Matthiesen, 2007, p. 208).

### Non-Standard Soil pH and Microbial Communities

In this study, we have explored standard and non-standard measurements of soil pH for the prediction of soil bacterial community composition. As numerous other studies have found (Bahram et al., 2018; Bartram et al., 2014; Delgado-Baquerizo et al., 2018; Rousk, Bååth, et al., 2010), soil bacterial community composition was strongly correlated with pH across both small and large regions (Figure 5). We hypothesized that soil pH values taken during the simulation of soil conditions (elevated carbon dioxide and typical solution:soil ratios) would more closely represent *in situ* conditions of microhabitats and therefore predict bacterial community composition better than standard soil pH values. This hypothesis was supported in the long-term pH manipulation field trial, but was not meaningfully supported in the cross-Wisconsin dataset (Figure 6). This suggests that, by lowering solution:soil ratios to more typical moisture levels of mineral soils, we were better able to represent the conditions experienced by microbial communities that reflected in their composition as measured by molecular methods. This also suggests that the non-standard *in situ* soil pH method developed here will apply well to soils of similar texture but poorly to soils of diverse texture. Overall, the range of soil pH values grew widely at low moisture whereas the range of soil pH values varied little from neutral at high moisture, namely the standard soil suspension method. This growing range of soil pH values measured under more typical conditions results in improved predictions of microbial community composition, suggesting further that standard soil pH, as it is currently measured, fails to distinguish differences in environmental conditions that are relevant to microbial life in soils.

Why, then, did similar changes in non-standard pH in the cross-Wisconsin dataset not result in similarly improved predictions of microbial community composition? While the soils from the pH manipulation trial were controlled and relatively similar in all characteristics except soil pH, the Wisconsin soil dataset was designed to be diverse in texture, organic matter, and other factors. Thus, pH had weaker explanatory power to begin with, due to the presence of other influential differences in the Wisconsin dataset. Furthermore, the mechanisms by which adjusting solution content affects pH may differ in different soils. Additionally, we should recognize that the long-term experimental pH plots had been amended with lime to raise the soil pH and sulfur to lower the soil pH. We cannot rule out that some excess unreacted amendment may have persisted in the samples, whose suspension during preparation for analysis might have dissolved and reacted to affect the analyte. This would potentially help explain why the higher soil pH values increased and the lower ones decreased at lower solution:soil ratios but would *not* explain why the pH values of the improved method were more accurately related to the composition of respective soil microbial communities.

Why did increasing CO_2_ levels not affect predictive power of pH measurements? The effects of increasing CO_2_ levels were more consistent across the range of pH levels, i.e., the intercept changed, but the slope changed less than it did when changing solution:soil ratios (Figures 2 and 3). Thus, it is not surprising that we did not gain predictive power from adjusting CO_2_ levels during measurements. If one is concerned about an extremely accurate measurement of pH, then it may be advisable to measure the soil solution under CO_2_ levels designed to mirror those of the soil. However, if one is interested primarily in predictive values in mineral soils, then these results suggest that such an approach is not essential. The measurement of the effects of CO_2_ on *in situ* soil pH, when this effect is measured in the future, may prove more significant. We might also consider whether the causes of high CO_2_ levels in a given soil−e.g., optimal moisture, temperature, and organic matter availability for microbial respiration−are more directly influential on microbial composition than their indirect (and perhaps transient) effects of elevating CO_2_ that shifts the pH of the soil solution.

Finally, a comment should be made on the assumptions underpinning the correlations between pH and microbial community composition. A PERMANOVA effectively tests for the presence of a linear relationship between microbial community dissimilarities and the variables of interest. As we are all well aware, pH is logarithmically related to 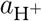. Although studies have historically found a significant and large relationship between pH and microbial community composition, there is no reason that the causative factors underpinning the specific effect of pH on soil microbial communities should be specifically proportional to the negative log of 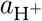 . That is to say, there is not an obvious reason that a 10x increase in 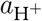 should have half the effect on the microbial community composition that a 100x increase does, nor would we necessarily expect differences in community composition to be linearly related to 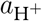 itself (Figure 6). It is important to consider this caveat when exponentiating soil pH values and performing statistical analyses with these calculated values in molar units.

### Recommendations

Because the microbial ecology of soil microorganisms, the acidity and acidification of soils, and the mechanisms by which soil bacteria survive are all of great relevance to sustainable crop production and biogeochemical models, non-standard soil pH values may offer both microbiologists and agonomists more targeted metrics to monitor and ultimately improve soil health (Meena, 2019, pp. 113–159). Unfortunately, whether and how to choose an appropriate non-standard protocol can be challenging, even if we recognize the need for alternate approaches. On the one hand, the use of non-standard methods of measuring soil acidity risks violating the commensurability of an investigator’s pH values to the standard soil pH values found in large databases (Minasny et al., 2011). On the other hand, the large diversity and variability through time of soil environments warrants diversification and customization of methods as well as the subsequent interpretation of the values that novel or adapted methods yield. For example, most soils collected at field capacity do not require the addition of excess analytical solution to extract soil solution via centrifugation (Geibe et al., 2006; Wolt, 1994, p. 104). A saturated peatland may require neither drying nor addition of solution but simply gentle centrifugation and analysis of the supernatant with a glass pH probe to obtain an informative measurement of pH. On the opposite extreme, a study of saline desert soils inhabited by plants having halotolerant root physiology would require the addition of a solution, almost certainly equal to or in excess of the typical 1 : 1 solution:soil ratio by mass, to create solution extract dilute enough for pH measurement. We must also continue (or begin) to ask what “soil pH” fundamentally means for frozen systems. In many regions of Earth’s surface, the soil solution is in solid phase for all or a large period of the year, whereby the solution is intractably shifted away from away from states resembling lab conditions.

We may reformulate soil pH measurement recommendations for the improved use of such values in microbial ecology, possibly viewing the elevated concentration of solutes and carbonate in the analytes of these sites as a means of both heightening the detection of important acids and bases found in typical soil solution by the glass probe as well as improving the representation of *in situ* conditions of soil microhabitats (Sumner, 1994). However, the further concentration of analyte beyond a 1 : 4 solution:soil ratio may cause the analyte to begin interfering with the functioning of the glass probe, which, as stated above, only functions without error *≤* 5% in analytes of ionic strength *≤* 0.01 moles per liter (Anderegg and Kholeif, 1994, p. 1521; Baucke, 2002, p. 774; Butler, 1998, pp. 462–463; Covert and Hore, 2016, pp. 235–238; de Levie, 2014, p. 615, 2010; Dobrovolskii et al., 2018, p. 87; Galster, 1991, p. 16; Sparks, 1998, p. 112; Spitzer and Pratt, 2011, p. 75; Volk and Rozen, 1977; Wright, 2007, p. 382; p. 1569; Pourbaix, 1974, p. 14; Ashcraft, 1957, p. 3, 1947, p. 29; Bates and Guggenheim, 1960, p. 167; Debye and Hückel, 1923, p. 197; Feldman, 1956, p. 1865, 1956, p. 1865; MacInnes, 1939, p. 148; Sena, 1972, Appendix 3).

Standard measurements of soil pH, such as those that populate our national or global soil databases, have been extremely useful for agronomy, and have also correlated strongly with bacterial community composition. However, we recognize that these methods offer only a limited representation of the acidity of soil microhabitats as they are experienced by microbes. By using methods for measuring soil acidity that simulate the *in situ* conditions of soils, we may improve the predictive models of the ecology of soil bacteria. The tools and equipment used here are all common to a molecular microbiology laboratory, and as such offer investigators the ability to miniaturize and concentrate the soil-solution suspension. Miniaturization of soil solution preparation also enables the analysis of more measurements at a higher throughput, as well as more readily simulating the conditions of soil microhabitats in the laboratory to measure *in situ* soil pH in a glove box to simulate non-standard atmospheric conditions, if desired. Such non-standard soil pH values have the potential to improve the modeling of temporal variability and enhance the characterization of study systems of both agronomists and microbial ecologists.

## Supporting information

All Soil Pit Profiles

## Funding and Acknowledgements

This work was supported by a Hatch grant (MSN210615) and by the UW-Madison O.N. Allen Professorship in Soil Science. This research was performed using the compute resources and assistance of the UW-Madison Center For High Throughput Computing (CHTC) in the Department of Computer Sciences. The CHTC is supported by UW-Madison, the Advanced Computing Initiative, the Wisconsin Alumni Research Foundation, the Wisconsin Institutes for Discovery, and the National Science Foundation, and is an active member of the Open Science Grid, which is supported by the National Science Foundation and the U.S. Department of Energy’s Office of Science. We would also like to thank Carrie Laboski, Phil Holman, Mattie Urrutia, the staff of the Wisconsin Agricultural Research Stations, Harry Read, Nayela Zeba, and Jaime Woolet.

## Supplemental Materials

### Supplemental Figures

**Supplemental Figure 1.**
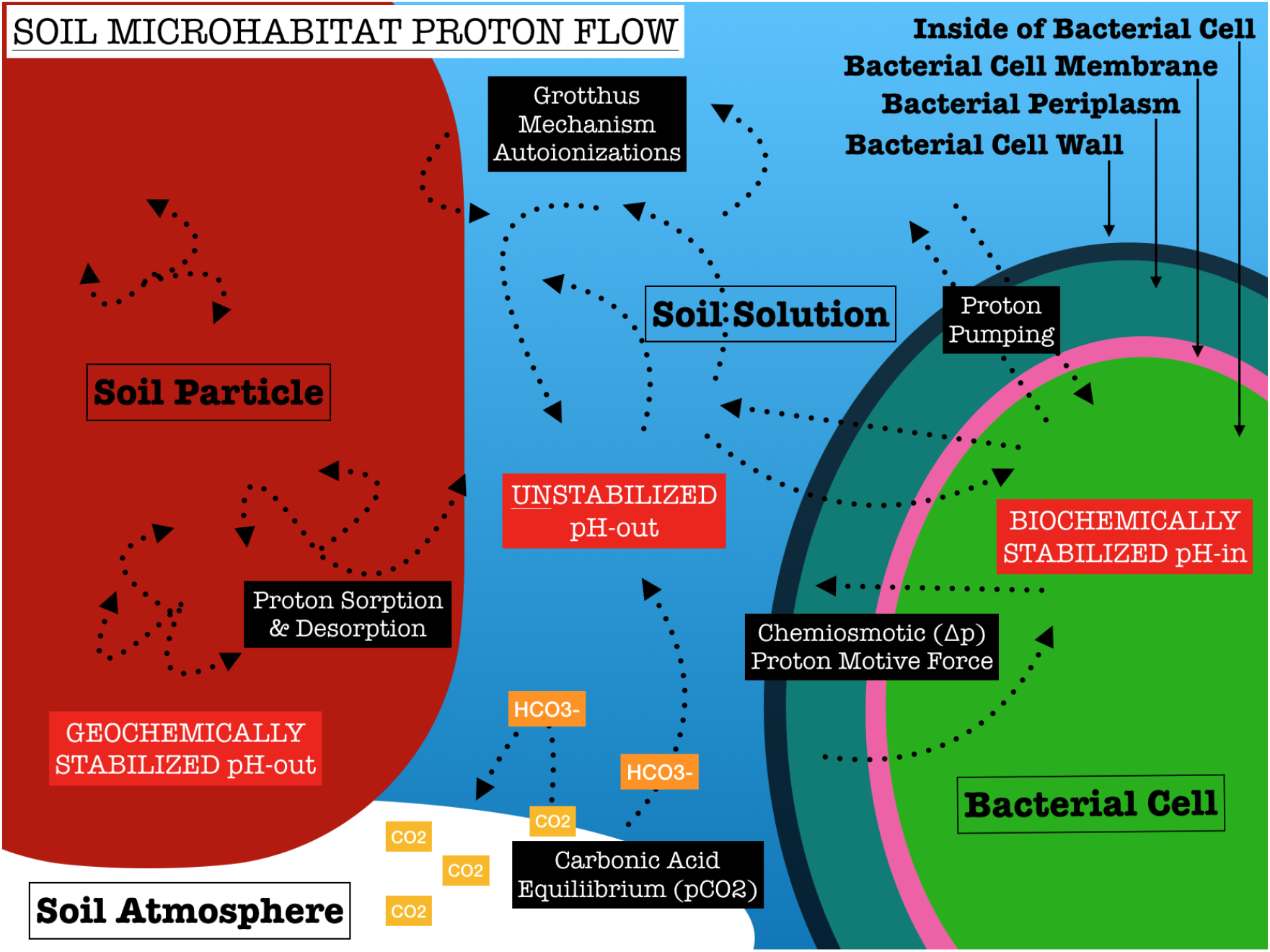
Soil microhabitat proton flow describes the biogeochemical processes connecting abiotic and biotic proton reservoirs.

**Supplemental Figure 2.**
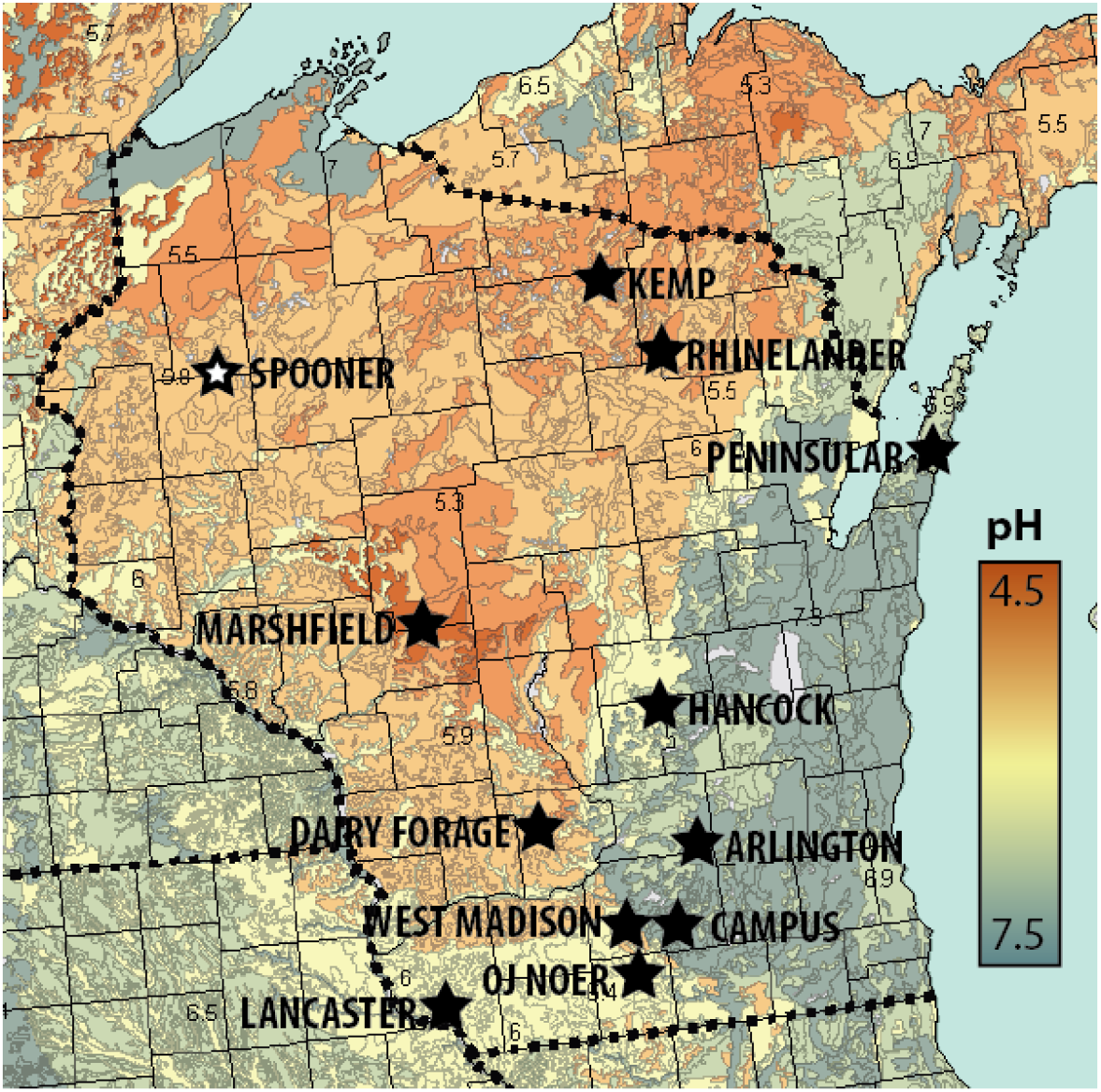
Map of field locations in Wisconsin with reference to the natural soil pH gradient across this region. Modified with permission from bonap.org (Kartesz, 2015).

**Supplemental Figure 3.**
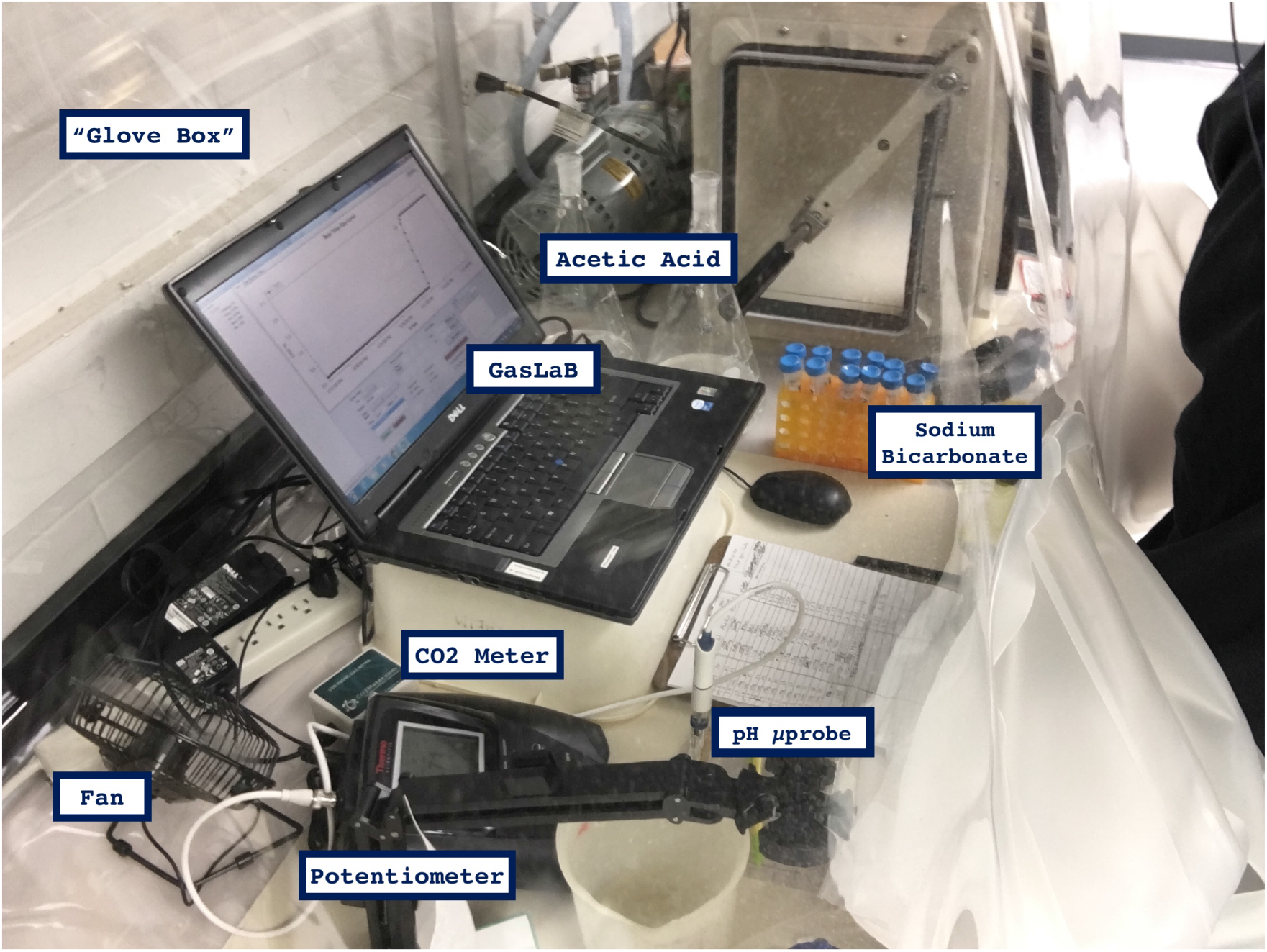
“Simulated soil pH” experimental rig, equipment, and reagents for acidimetry under elevated carbon dioxide resembling a typical *in situ* soil atmosphere.

**Supplemental Figure 4.**
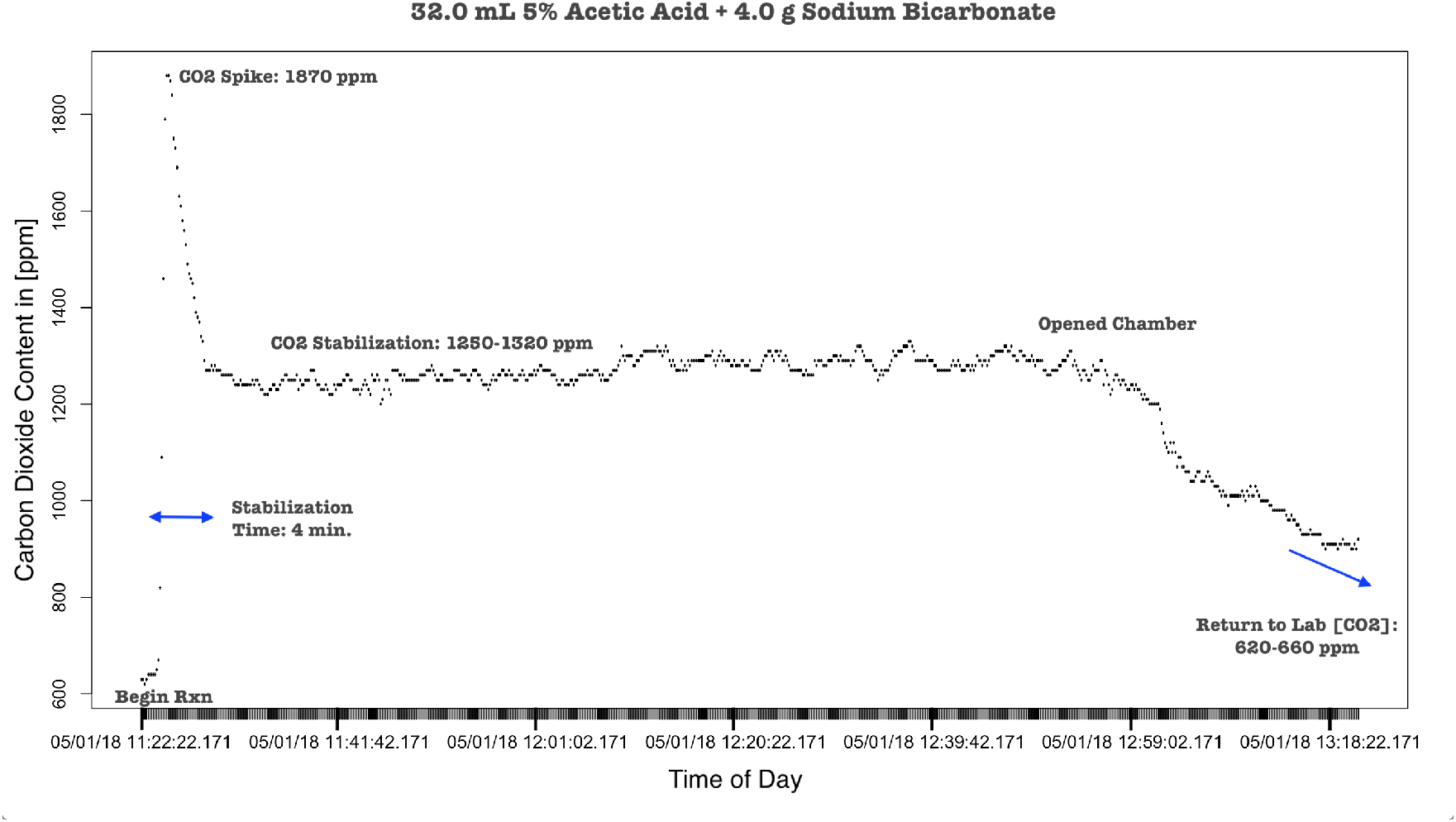
Test showing that the chamber (“glove box”) and gas analyzer provide a stable and controllable elevated carbon dioxide atmosphere for sufficient time and elvels to perform chemical analyses such as acidimetry while simulating soil atmospheric conditions. The carbon dioxide content exhibits an initial spike, stabilization, then an extended period whereby the chamber has an elevated carbon dioxide creating a partially simulated soil atmosphere. The chamber can be opened and vented once more to return to laboratory carbon dioxide levels.

**Supplemental Figure 5.**
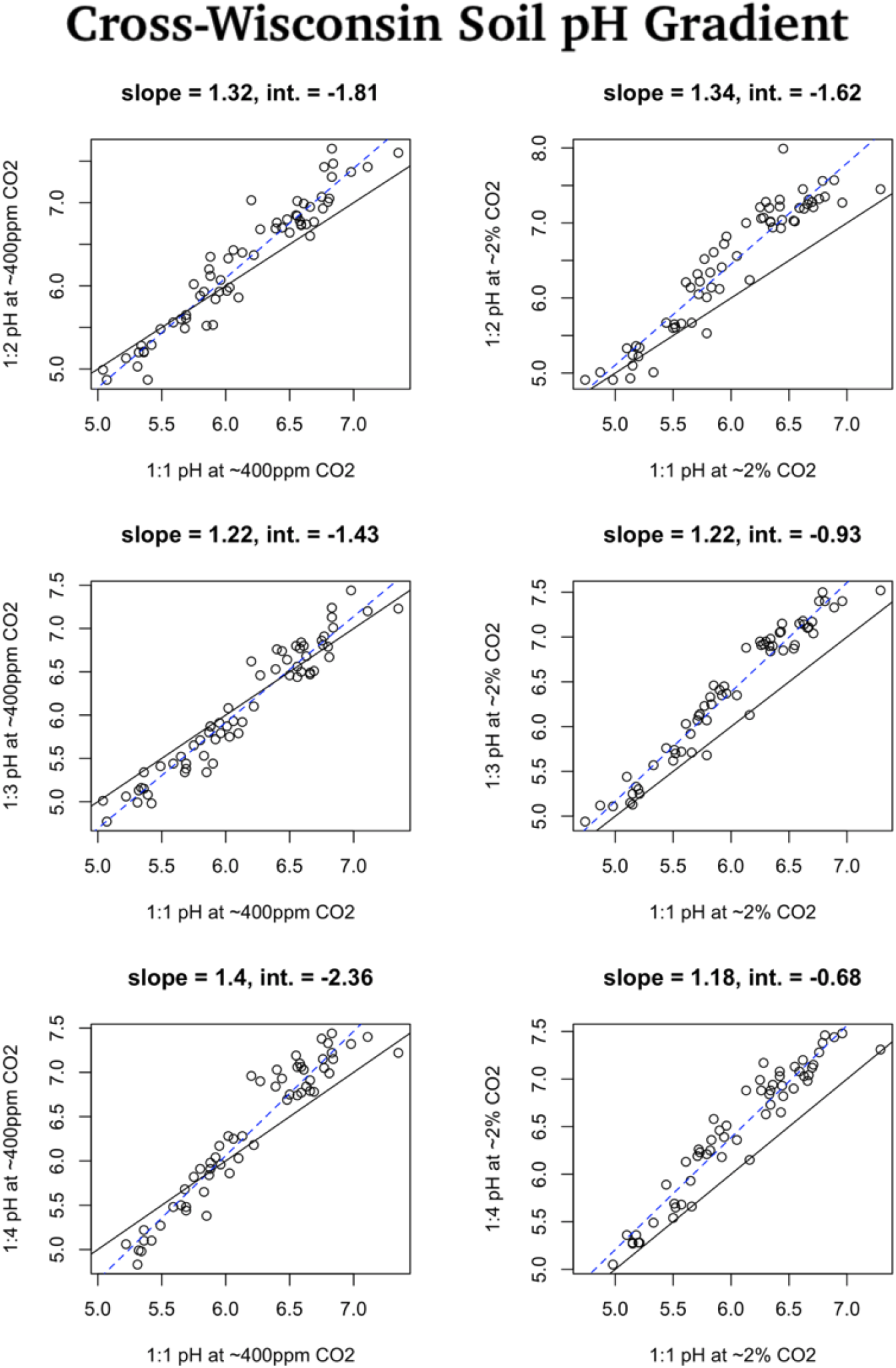
Standard soil pH (solution:soil ratio) of cross-Wisconsin soils compared to three other ratios (2 : 1, 3 : 1, 4 : 1) at two levels of carbon dioxide (0.04% ppm and 2.2%(±0.05)).

**Supplemental Figure 6.**
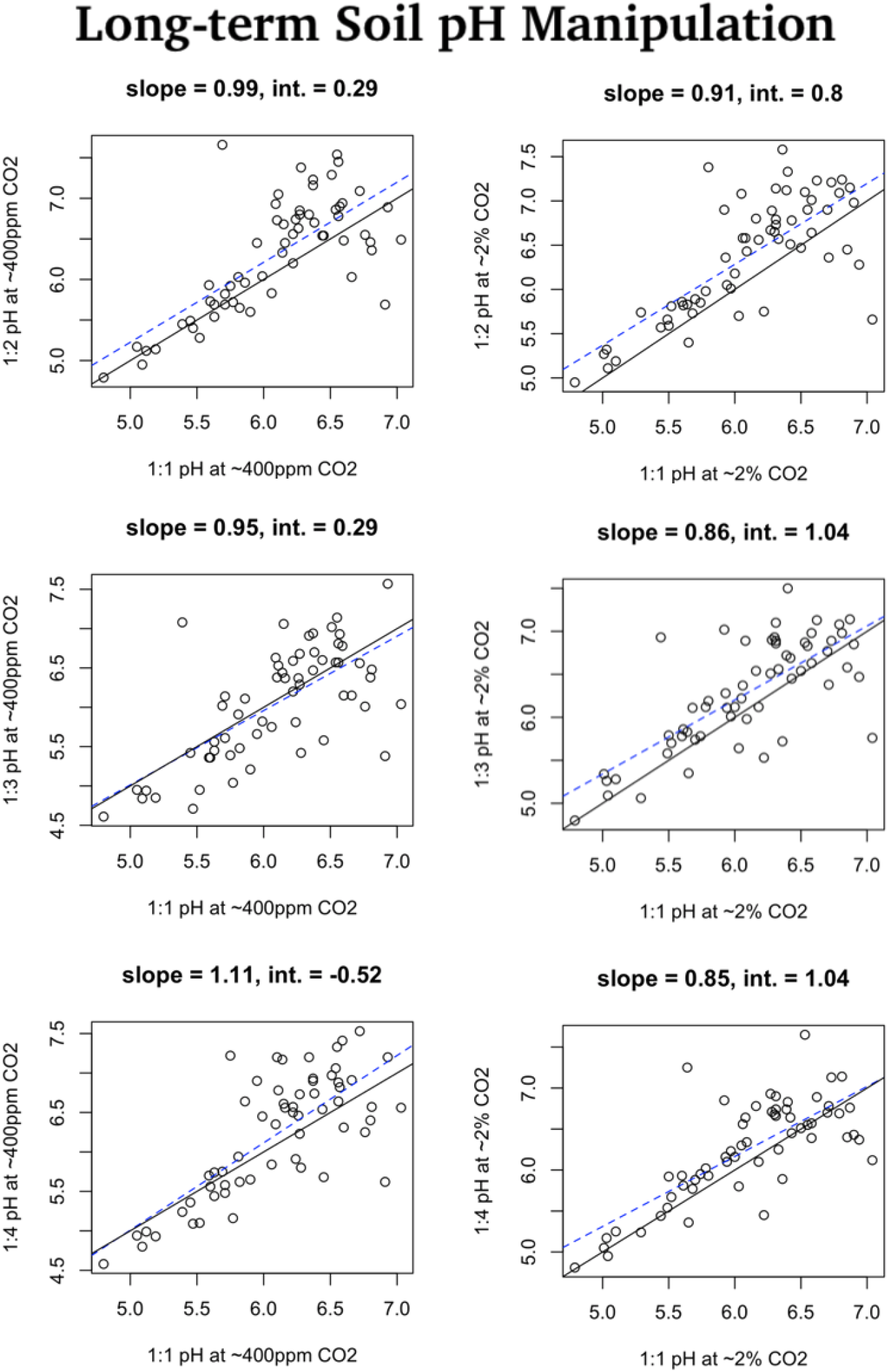
Standard soil pH (1 : 1 solution:soil ratio) of long-term pH manipulation soils compared to three other ratios (2 : 1, 3 : 1, 4 : 1) at two levels of carbon dioxide (0.04% and 2.2%(±0.05)).

**Supplemental Figure 7.**
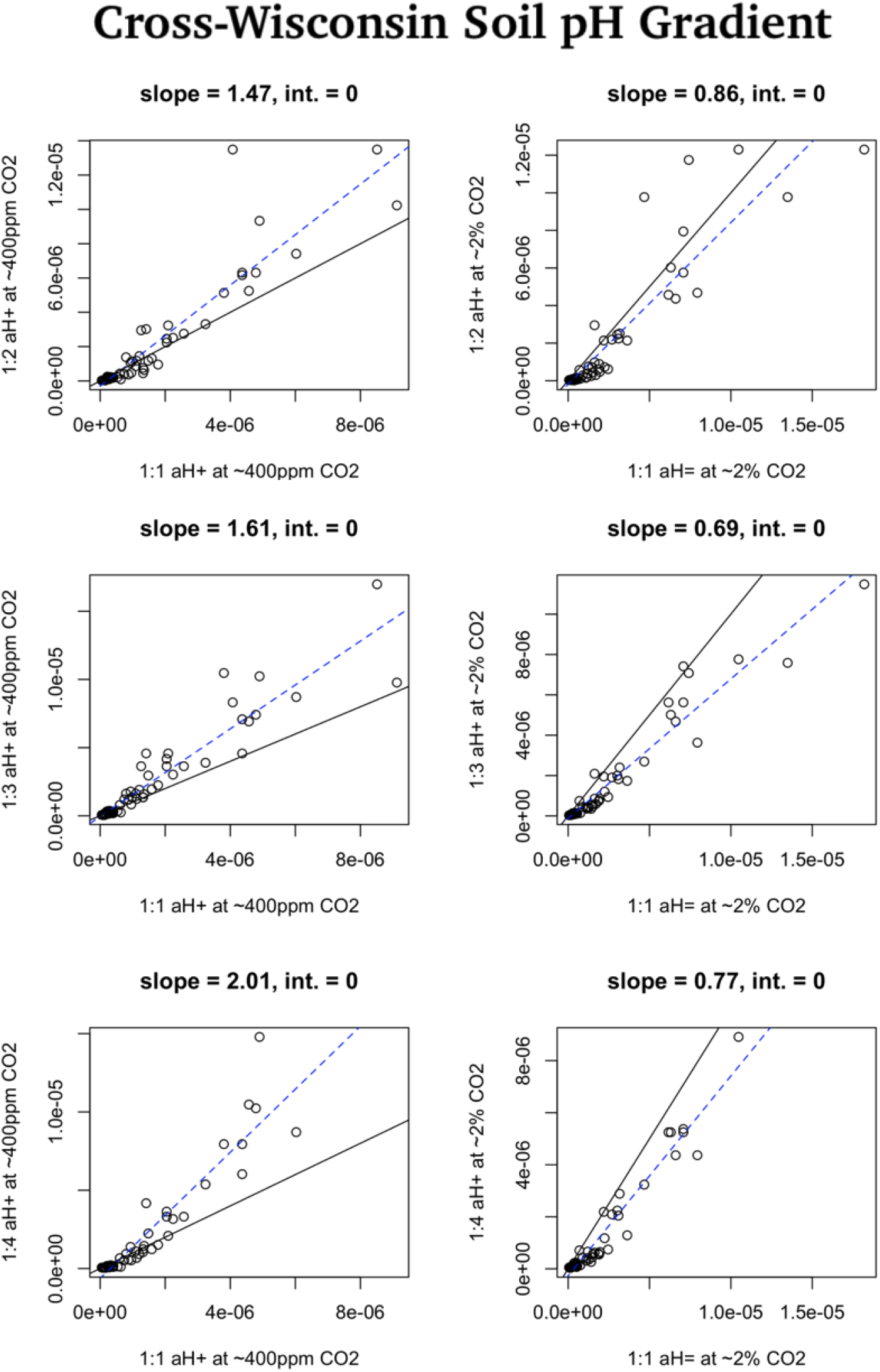
Soil 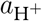 (1 : 1 solution:soil ratio) of cross-Wisconsin soils compared to three other ratios (2 : 1, 3 : 1, 4 : 1) at two levels of carbon dioxide (0.04% and 2.2%(±0.05)).

**Supplemental Figure 8.**
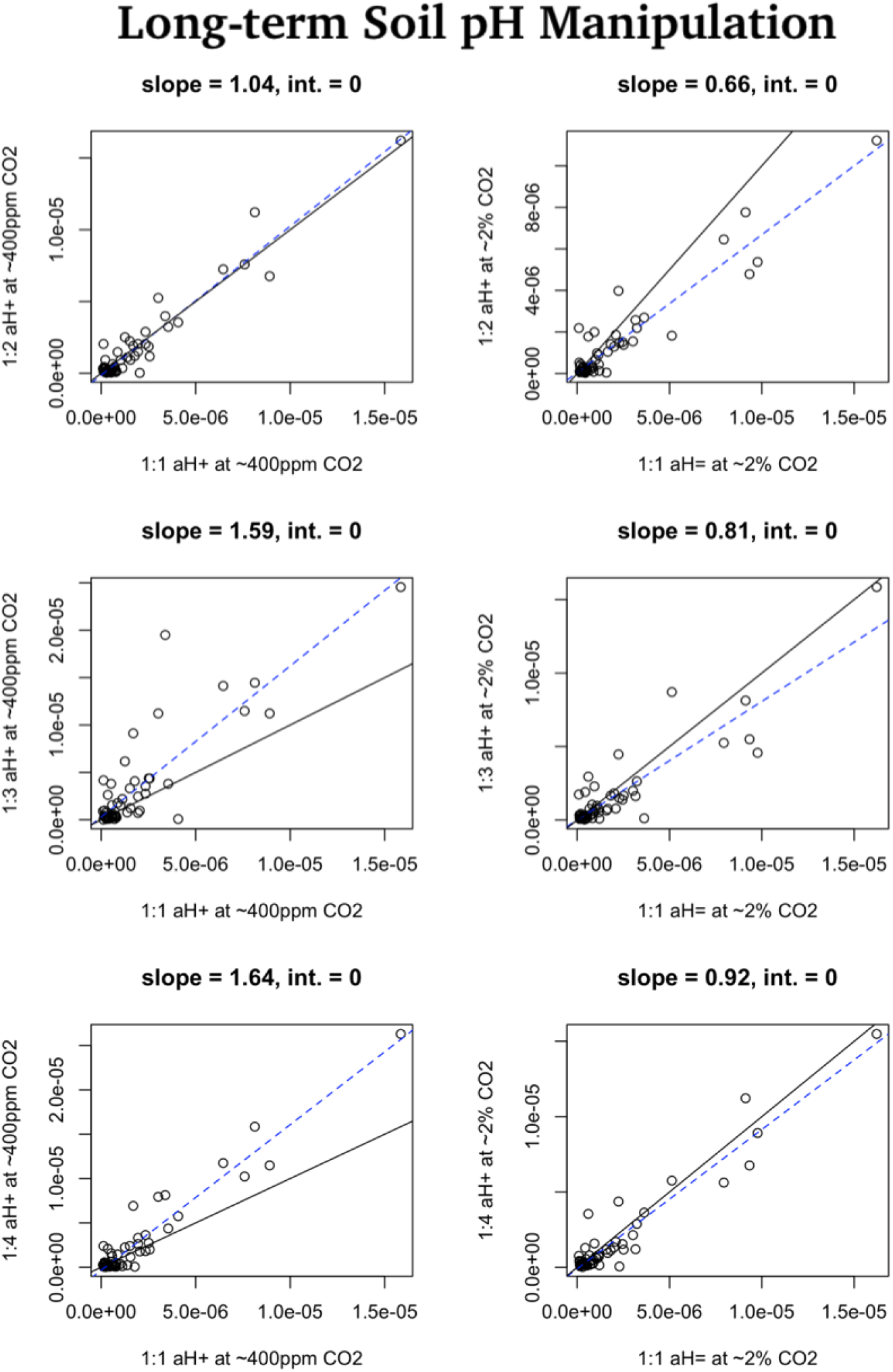
Soil 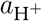 (1 : 1 solution:soil ratio) of long-term pH manipulation soils compared to three other ratios (2 : 1, 3 : 1, 4 : 1) at two levels of carbon dioxide (0.04% and 2.2%(±0.05)).

**Supplemental Figure 9.**
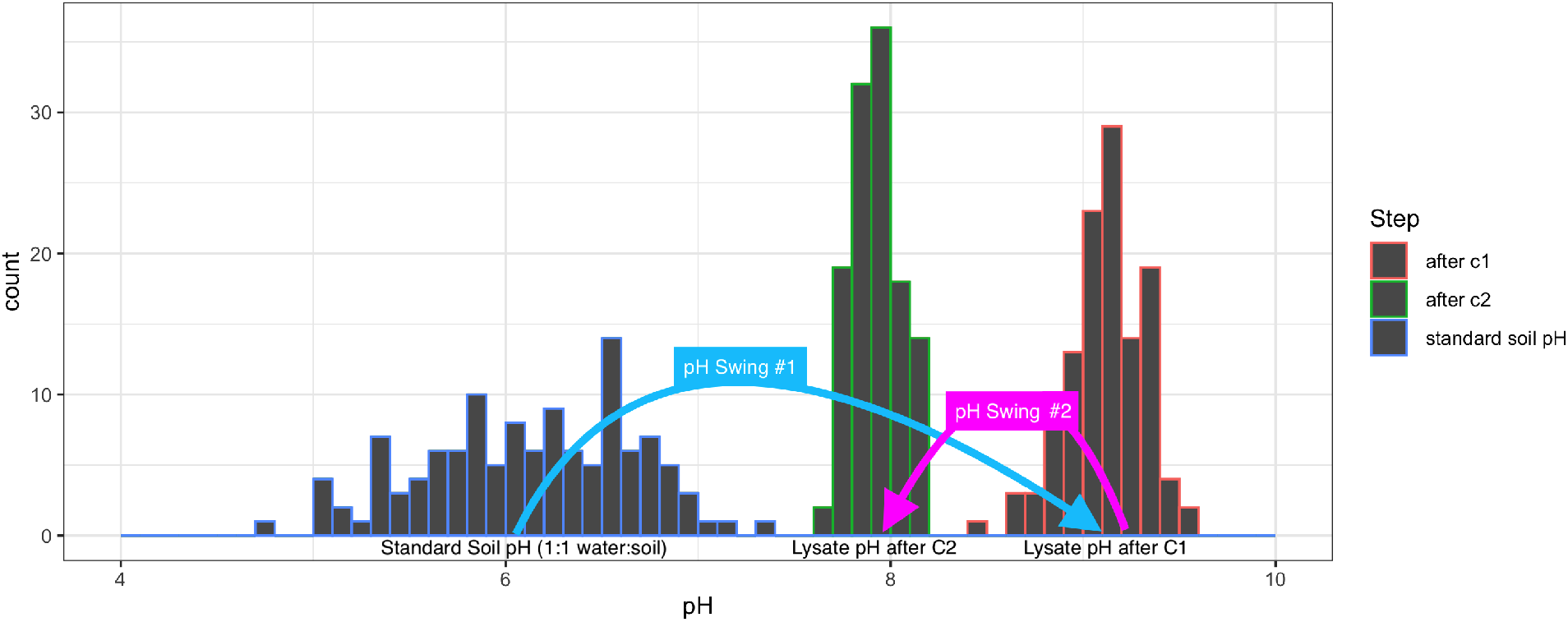
Histograms of standard soil pH and the pH of the lysate supernatants after treatment with buffers “C1” and “C2”, respectively, of the first two steps (“pH swings”) of the soil DNA extraction protocol.

**Supplemental Figure 10.**
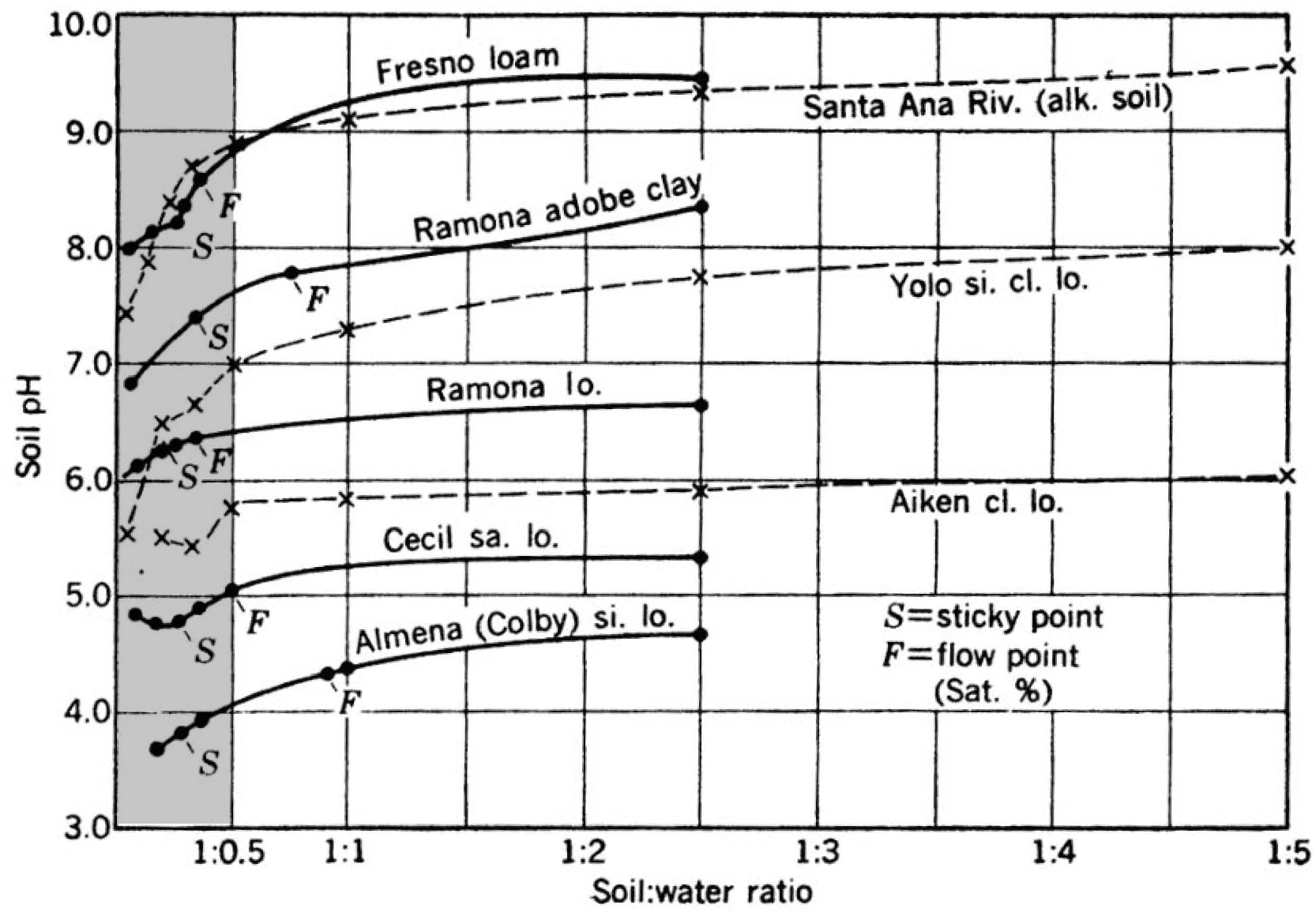
The slurry-to-paste dilution pH differentiation trend, adapted from Jackson (1958, p. 43). Solid lines were derived from Chapman et al. (1941) and dashed lines were derived from Huberty and Haas (1940). The moisture levels considered typical and representative of *in situ* conditions that are much less diluted than standard soil pH (1 : 1 solution:soil by mass) in this study are greyed.

### Supplemental Notes

Supplemental Note 1: pH Probe Details

The pH electrode had a shaft length of 60 [mm] and a diameter of 3 [mm] with a built-in ARGENTHAL reference system of 3.0 [M] KCl reference electrolyte. The probe was stored in either 3.0 [M] KCl saturated electrolyte solution or InLab storage solution (Material No. 30111142). The probe’s glass was made from U Glass with a membrane resistance of 600 [Mohm]. The probe, owing to the sorption of solution to its surface, will remove approximately 5 [*μ*L] per measurement, and the accuracy during this study was < %15 while performing ≤ 3 repeated measurements of the same extracts in different simulated conditions.

Supplemental Note 2: “pH Swings” of Soil DNA Lysate During Extraction

Refer to Supplemental Figure 9. The pH values of miniaturized analytes of the first two steps of a standard soil DNA extraction protocol were measured. Two sets of DNA extraction kits with bead-beating tubes and solutions C1 and C2, which are identical to the solutions and materials used in the PowerLyzer PowerSoil DNA Isolation Kit used for 16S amplicon sequencing in this study, were used to generate lysates of the first two steps of the soil DNA extraction. Excess addition of C1 and C2 solutions allowed for the removal of small aliquots of solution without disrupting the chemical events and buffers of the first steps of DNA extraction. 100 [*μ*L] was removed from the lysate after the addition and bead-beating with solution C1, and another 100 [*μ*L] was removed from the lysate after the addition of solution C2. The pH values of these solutions (“after C1” and “after C2”) were compared to the standard soil pH values (i.e., 1 : 1 solution:soil ratio at ambient carbon dioixide levels).

The first pH swings to from the the more variable and acidic standard soil pH values (1 : 1 solution:soil), then the second pH swings down to approximately, narrowing the range of pH values as the DNA extraction progresses (Supplemental Figure 11). The acidic soils (< 5.5) were nearly 100× more acidic than the neutral-to-basic soils (< 7.0) according to their standard soil pH measurement, but the DNA extraction kit treated these soils with an identical alkaline buffer in the first step.

Although solution C1 pH and solution C2 pH were both significant predictors of community composition on their own, after controlling for other soil properties, neither was a significant predictor, nor were they correlated with soil pH measurements (*p*_C1_ = 0.46 and *p*_C2_ = 0.69). However, they were significantly negatively correlated with total Ca (*p* < 0.001, *R*^2^ = 0.41).

### Supplemental Tables

**Supplemental Table 1.**
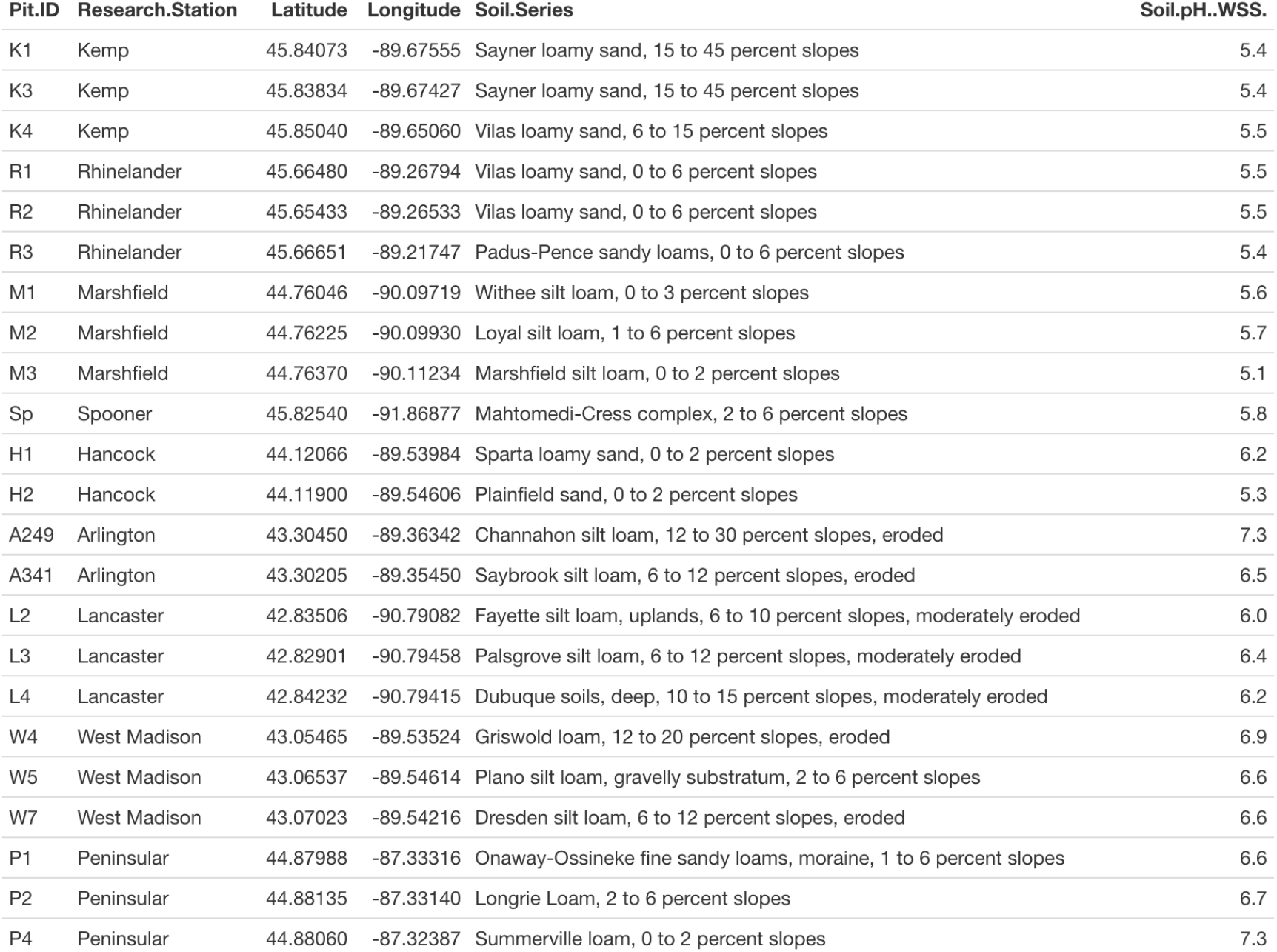
Latitude, longitude, soil series, and soil pH of field sites according to the Web Soil Survey database.

**Supplemental Table 2.** Primers used to amplify 16S gene.

| Sample.Name | Fwd_Primer_ID | Fwd_Primer_Barcode | Rev_Primer_ID | Rev_Primer_Barcode |
| --- | --- | --- | --- | --- |
| 001.K1.0.17 | 515f_SA501 | ATCGTACG | 806r_SA701 | AACTCTCG |
| 002.K1.17.45 | 515f_SB501 | CTACTATA | 806r_SA709 | GTCGTAGT |
| 003.K1.45.60 | 515f_SC501 | ACGACGTG | 806r_SB705 | CGTAGATC |
| 004.K2.Muck | 515f_SA502 | ACTATCTG | 806r_SA702 | ACTATGTC |
| 005.K3.0.15 | 515f_SB502 | CGTTACTA | 806r_SA710 | TAGCAGAC |
| 006.K3.15.35 | 515f_SC502 | ATATACAC | 806r_SB706 | CTCGTTAC |
| 007.K3.35.50 | 515f_SA503 | TAGCGAGT | 806r_SA703 | AGTAGCGT |
| 008.K4.0.15 | 515f_SB503 | AGAGTCAC | 806r_SA711 | TCATAGAC |
| 009.K4.15.30 | 515f_SC503 | CGTCGCTA | 806r_SB707 | GCGCACGT |
| 010.K4.30.50 | 515f_SA504 | CTGCGTGT | 806r_SA704 | CAGTGAGT |
| 011.R1.0.27 | 515f_SB504 | TACGAGAC | 806r_SA712 | TCGCTATA |
| 012.R1.27.50 | 515f_SC504 | CTAGAGCT | 806r_SB708 | GGTACTAT |
| 013.R1.50.70 | 515f_SA505 | TCATCGAG | 806r_SA705 | CGTACTCA |
| 014.R2.0.30 | 515f_SB505 | ACGTCTCG | 806r_SB701 | AAGTCGAG |
| 015.R2.30.45 | 515f_SC505 | GCTCTAGT | 806r_SB709 | GTATACGC |
| 016.R2.45.60 | 515f_SA506 | CGTGAGTG | 806r_SA706 | CTACGCAG |
| 017.R2.60.100 | 515f_SB506 | TCGACGAG | 806r_SB702 | ATACTTCG |
| 018.R3.0.20 | 515f_SC506 | GACACTGA | 806r_SB710 | TACGAGCA |
| 019.R3.20.30 | 515f_SA507 | GGATATCT | 806r_SA707 | GGAGACTA |
| 020.M1.0.31 | 515f_SB507 | GATCGTGT | 806r_SB703 | AGCTGCTA |
| 021.M1.31.50 | 515f_SC507 | TGCGTACG | 806r_SB711 | TCAGCGTT |
| 022.M1.50.70 | 515f_SA508 | GACACCGT | 806r_SA708 | GTCGCTCG |
| 023.M2.0.24 | 515f_SB508 | GTCAGATA | 806r_SB704 | CATAGAGA |
| 024.M2.24.38 | 515f_SC508 | TAGTGTAG | 806r_SB712 | TCGCTACG |
| 025.M2.38.55 | 515f_SA501 | ATCGTACG | 806r_SB712 | TCGCTACG |
| 026.M3.0.15 | 515f_SB501 | CTACTATA | 806r_SA701 | AACTCTCG |
| 027.M3.15.30 | 515f_SC501 | ACGACGTG | 806r_SA709 | GTCGTAGT |
| 028.S.0.30 | 515f_SA502 | ACTATCTG | 806r_SB705 | CGTAGATC |
| 029.S.30.60 | 515f_SB502 | CGTTACTA | 806r_SA702 | ACTATGTC |
| 030.H1.0.30 | 515f_SC502 | ATATACAC | 806r_SA710 | TAGCAGAC |
| 031.H1.30.40 | 515f_SA503 | TAGCGAGT | 806r_SB706 | CTCGTTAC |
| 032.H1.40.60 | 515f_SB503 | AGAGTCAC | 806r_SA703 | AGTAGCGT |
| 033.H2.0.30 | 515f_SC503 | CGTCGCTA | 806r_SA711 | TCATAGAC |
| 034.H2.30.60 | 515f_SA504 | CTGCGTGT | 806r_SB707 | GCGCACGT |
| 035.A249.0.35 | 515f_SB504 | TACGAGAC | 806r_SA704 | CAGTGAGT |
| 036.A249.35.60 | 515f_SC504 | CTAGAGCT | 806r_SA712 | TCGCTATA |
| 037.A341.0.33 | 515f_SA505 | TCATCGAG | 806r_SB708 | GGTACTAT |
| 038.A341.33.55 | 515f_SB505 | ACGTCTCG | 806r_SA705 | CGTACTCA |
| 039.A341.55.75 | 515f_SC505 | GCTCTAGT | 806r_SB701 | AAGTCGAG |
| 040.A341.75.85 | 515f_SA506 | CGTGAGTG | 806r_SB709 | GTATACGC |
| 041.L2.0.23 | 515f_SB506 | TCGACGAG | 806r_SA706 | CTACGCAG |
| 042.L2.23.45 | 515f_SC506 | GACACTGA | 806r_SB702 | ATACTTCG |
| 043.L3.0.12 | 515f_SA507 | GGATATCT | 806r_SB710 | TACGAGCA |
| 044.L3.12.20 | 515f_SB507 | GATCGTGT | 806r_SA707 | GGAGACTA |
| 045.L3.20.40 | 515f_SC507 | TGCGTACG | 806r_SB703 | AGCTGCTA |
| 046.L4.0.10 | 515f_SA508 | GACACCGT | 806r_SB711 | TCAGCGTT |
| 047.L4.10.20 | 515f_SB508 | GTCAGATA | 806r_SA708 | GTCGCTCG |
| 048.L4.20.40 | 515f_SC508 | TAGTGTAG | 806r_SB704 | CATAGAGA |
| 049.W3.Compost | 515f_SA501 | ATCGTACG | 806r_SB704 | CATAGAGA |
| 050.W4.0.28 | 515f_SB501 | CTACTATA | 806r_SB712 | TCGCTACG |
| 051.W4.28.45 | 515f_SC501 | ACGACGTG | 806r_SA701 | AACTCTCG |
| 052.W4.45.55 | 515f_SA502 | ACTATCTG | 806r_SA709 | GTCGTAGT |
| 053.W5.0.35 | 515f_SB502 | CGTTACTA | 806r_SB705 | CGTAGATC |
| 054.W5.35.65 | 515f_SC502 | ATATACAC | 806r_SA702 | ACTATGTC |
| 055.W7.0.15 | 515f_SA503 | TAGCGAGT | 806r_SA710 | TAGCAGAC |
| 056.W7.15.30 | 515f_SB503 | AGAGTCAC | 806r_SB706 | CTCGTTAC |
| 057.P1.0.30 | 515f_SC503 | CGTCGCTA | 806r_SA703 | AGTAGCGT |
| 058.P1.30.45 | 515f_SA504 | CTGCGTGT | 806r_SA711 | TCATAGAC |
| 059.P1.45.55 | 515f_SB504 | TACGAGAC | 806r_SB707 | GCGCACGT |
| 060.P2.0.20 | 515f_SC504 | CTAGAGCT | 806r_SA704 | CAGTGAGT |
| 061.P2.20.45 | 515f_SA505 | TCATCGAG | 806r_SA712 | TCGCTATA |
| 062.P2.45.55 | 515f_SB505 | ACGTCTCG | 806r_SB708 | GGTACTAT |
| 063.P4.0.25 | 515f_SC505 | GCTCTAGT | 806r_SA705 | CGTACTCA |
| 064.P4.25.35 | 515f_SA506 | CGTGAGTG | 806r_SB701 | AAGTCGAG |
| 065.P4.35.50 | 515f_SB506 | TCGACGAG | 806r_SB709 | GTATACGC |
| 181.Sp11.Cr31.0.20 | 515f_SC506 | GACACTGA | 806r_SA706 | CTACGCAG |
| 182.Sp11.Cr32.0.16 | 515f_SA507 | GGATATCT | 806r_SB702 | ATACTTCG |
| 183.Sp11.Cr33.0.20 | 515f_SB507 | GATCGTGT | 806r_SB710 | TACGAGCA |
| 184.Sp12.Cr34.0.16 | 515f_SC507 | TGCGTACG | 806r_SA707 | GGAGACTA |
| 185.Sp12.Cr35.0.18 | 515f_SA508 | GACACCGT | 806r_SB703 | AGCTGCTA |
| 186.Sp12.Cr36.0.20 | 515f_SB508 | GTCAGATA | 806r_SB711 | TCAGCGTT |
| 187.Sp13.Cr37.0.20 | 515f_SC508 | TAGTGTAG | 806r_SA708 | GTCGCTCG |
| 188.Sp13.Cr38.0.20 | 515f_SA501 | ATCGTACG | 806r_SA708 | GTCGCTCG |
| 189.Sp13.Cr39.0.19 | 515f_SB501 | CTACTATA | 806r_SB704 | CATAGAGA |
| 190.Sp14.Cr40.0.17 | 515f_SC501 | ACGACGTG | 806r_SB712 | TCGCTACG |
| 191.Sp14.Cr41.0.18 | 515f_SA502 | ACTATCTG | 806r_SA701 | AACTCTCG |
| 192.Sp14.Cr42.0.20 | 515f_SB502 | CGTTACTA | 806r_SA709 | GTCGTAGT |
| 193.Sp15.Cr43.0.18 | 515f_SC502 | ATATACAC | 806r_SB705 | CGTAGATC |
| 194.Sp15.Cr44.0.17 | 515f_SA503 | TAGCGAGT | 806r_SA702 | ACTATGTC |
| 195.Sp15.Cr45.0.20 | 515f_SB503 | AGAGTCAC | 806r_SA710 | TAGCAGAC |
| 196.Sp16.Cr46.0.20 | 515f_SC503 | CGTCGCTA | 806r_SB706 | CTCGTTAC |
| 197.Sp16.Cr47.0.17 | 515f_SA504 | CTGCGTGT | 806r_SA703 | AGTAGCGT |
| 198.Sp16.Cr48.0.20 | 515f_SB504 | TACGAGAC | 806r_SA711 | TCATAGAC |
| 199.Sp17.Cr49.0.20 | 515f_SC504 | CTAGAGCT | 806r_SB707 | GCGCACGT |
| 200.Sp17.Cr50.0.18 | 515f_SA505 | TCATCGAG | 806r_SA704 | CAGTGAGT |
| 201.Sp17.Cr51.0.18 | 515f_SB505 | ACGTCTCG | 806r_SA712 | TCGCTATA |
| 202.Sp18.Cr52.0.18 | 515f_SC505 | GCTCTAGT | 806r_SB708 | GGTACTAT |
| 203.Sp18.Cr53.0.19 | 515f_SA506 | CGTGAGTG | 806r_SA705 | CGTACTCA |
| 204.Sp18.Cr54.0.20 | 515f_SB506 | TCGACGAG | 806r_SB701 | AAGTCGAG |
| 205.Sp19.Cr55.0.20 | 515f_SC506 | GACACTGA | 806r_SB709 | GTATACGC |
| 206.Sp19.Cr56.0.18 | 515f_SA507 | GGATATCT | 806r_SA706 | CTACGCAG |
| 207.Sp19.Cr57.0.18 | 515f_SB507 | GATCGTGT | 806r_SB702 | ATACTTCG |
| 208.Sp20.Cr58.0.20 | 515f_SC507 | TGCGTACG | 806r_SB710 | TACGAGCA |
| 209.Sp20.Cr59.0.19 | 515f_SA508 | GACACCGT | 806r_SA707 | GGAGACTA |
| 210.Sp20.Cr60.0.20 | 515f_SB508 | GTCAGATA | 806r_SB703 | AGCTGCTA |
| 211.Sp1.Cr1.0.20 | 515f_SC508 | TAGTGTAG | 806r_SB711 | TCAGCGTT |
| 212.Sp1.Cr2.0.20 | 515f_SA501 | ATCGTACG | 806r_SB711 | TCAGCGTT |
| 213.Sp1.Cr3.0.20 | 515f_SB501 | CTACTATA | 806r_SA708 | GTCGCTCG |
| 214.Sp2.Cr4.0.20 | 515f_SC501 | ACGACGTG | 806r_SB704 | CATAGAGA |
| 215.Sp2.Cr5.0.20 | 515f_SA502 | ACTATCTG | 806r_SB712 | TCGCTACG |
| 216.Sp2.Cr6.0.20 | 515f_SB502 | CGTTACTA | 806r_SA701 | AACTCTCG |
| 217.Sp3.Cr7.0.20 | 515f_SC502 | ATATACAC | 806r_SA709 | GTCGTAGT |
| 218.Sp3.Cr8.0.20 | 515f_SA503 | TAGCGAGT | 806r_SB705 | CGTAGATC |
| 219.Sp3.Cr9.0.20 | 515f_SB503 | AGAGTCAC | 806r_SA702 | ACTATGTC |
| 220.Sp4.Cr10.0.20 | 515f_SC503 | CGTCGCTA | 806r_SA710 | TAGCAGAC |
| 221.Sp4.Cr11.0.20 | 515f_SA504 | CTGCGTGT | 806r_SB706 | CTCGTTAC |
| 222.Sp4.Cr12.0.20 | 515f_SB504 | TACGAGAC | 806r_SA703 | AGTAGCGT |
| 223.Sp5.Cr13.0.20 | 515f_SC504 | CTAGAGCT | 806r_SA711 | TCATAGAC |
| 224.Sp5.Cr14.0.20 | 515f_SA505 | TCATCGAG | 806r_SB707 | GCGCACGT |
| 225.Sp5.Cr15.0.20 | 515f_SB505 | ACGTCTCG | 806r_SA704 | CAGTGAGT |
| 226.Sp6.Cr16.0.15 | 515f_SC505 | GCTCTAGT | 806r_SA712 | TCGCTATA |
| 227.Sp6.Cr17.0.20 | 515f_SA506 | CGTGAGTG | 806r_SB708 | GGTACTAT |
| 228.Sp6.Cr18.0.20 | 515f_SB506 | TCGACGAG | 806r_SA705 | CGTACTCA |
| 229.Sp7.Cr19.0.20 | 515f_SC506 | GACACTGA | 806r_SB701 | AAGTCGAG |
| 230.Sp7.Cr20.0.20 | 515f_SA507 | GGATATCT | 806r_SB709 | GTATACGC |
| 231.Sp7.Cr21.0.13 | 515f_SB507 | GATCGTGT | 806r_SA706 | CTACGCAG |
| 232.Sp8.Cr22.0.20 | 515f_SC507 | TGCGTACG | 806r_SB702 | ATACTTCG |
| 233.Sp8.Cr23.0.20 | 515f_SA508 | GACACCGT | 806r_SB710 | TACGAGCA |
| 234.Sp8.Cr24.0.20 | 515f_SB508 | GTCAGATA | 806r_SA707 | GGAGACTA |
| 235.Sp9.Cr25.0.15 | 515f_SC508 | TAGTGTAG | 806r_SB703 | AGCTGCTA |
| 236.Sp9.Cr26.0.20 | 515f_SA501 | ATCGTACG | 806r_SB703 | AGCTGCTA |
| 237.Sp9.Cr27.0.15 | 515f_SB501 | CTACTATA | 806r_SB711 | TCAGCGTT |
| 238.Sp10.Cr28.0.20 | 515f_SC501 | ACGACGTG | 806r_SA708 | GTCGCTCG |
| 239.Sp10.Cr29.0.15 | 515f_SA502 | ACTATCTG | 806r_SB704 | CATAGAGA |
| 240.Sp10.Cr30.0.20 | 515f_SB502 | CGTTACTA | 806r_SB712 | TCGCTACG |

