## Supplementary figures and images for "Standard and Non-Standard Measurements of Acidity and the Bacterial Ecology of Northern Temperate Mineral Soils"

### a249-0cm-60cm.jpg

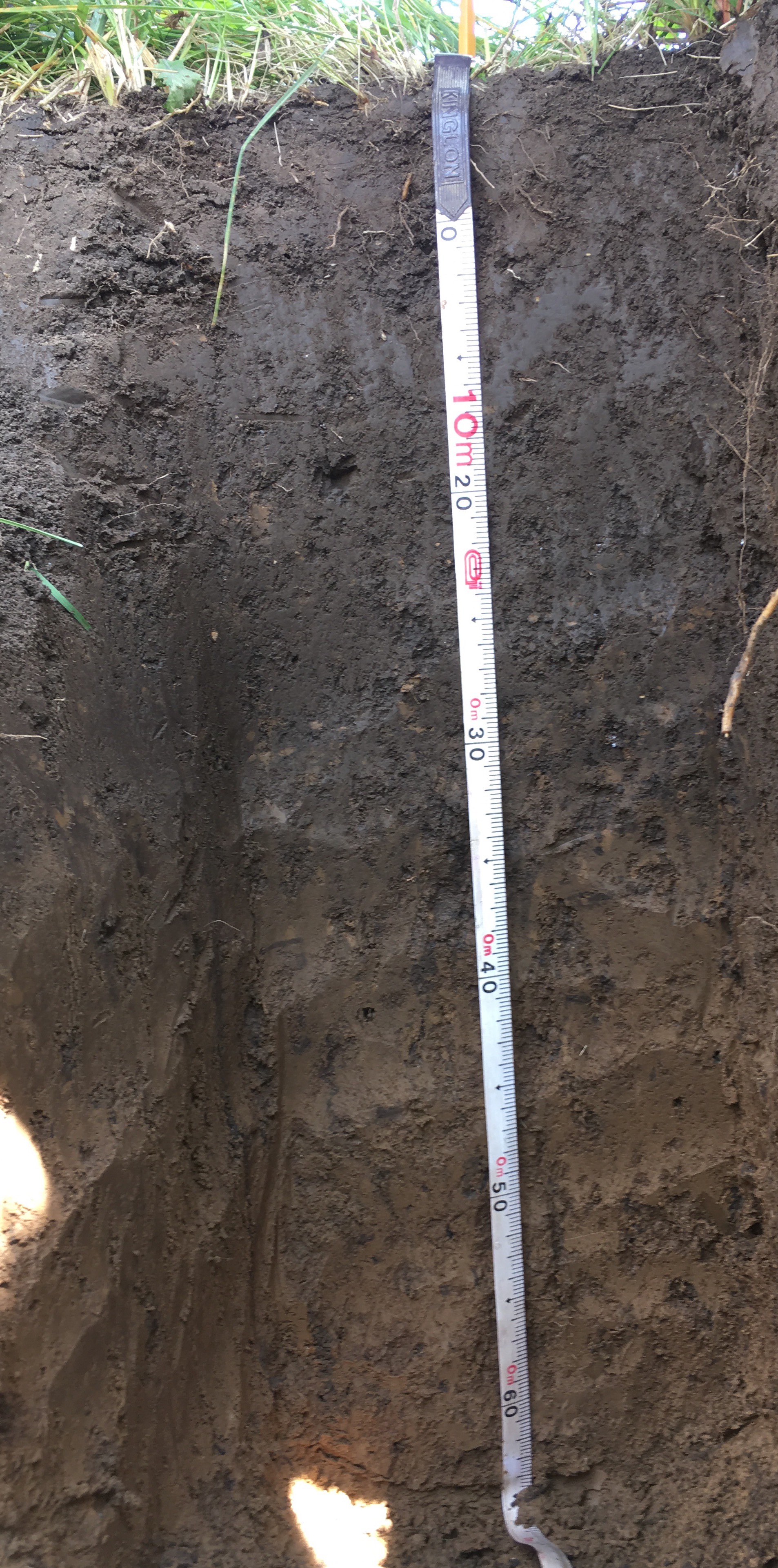

### a341-0cm-60cm.jpg

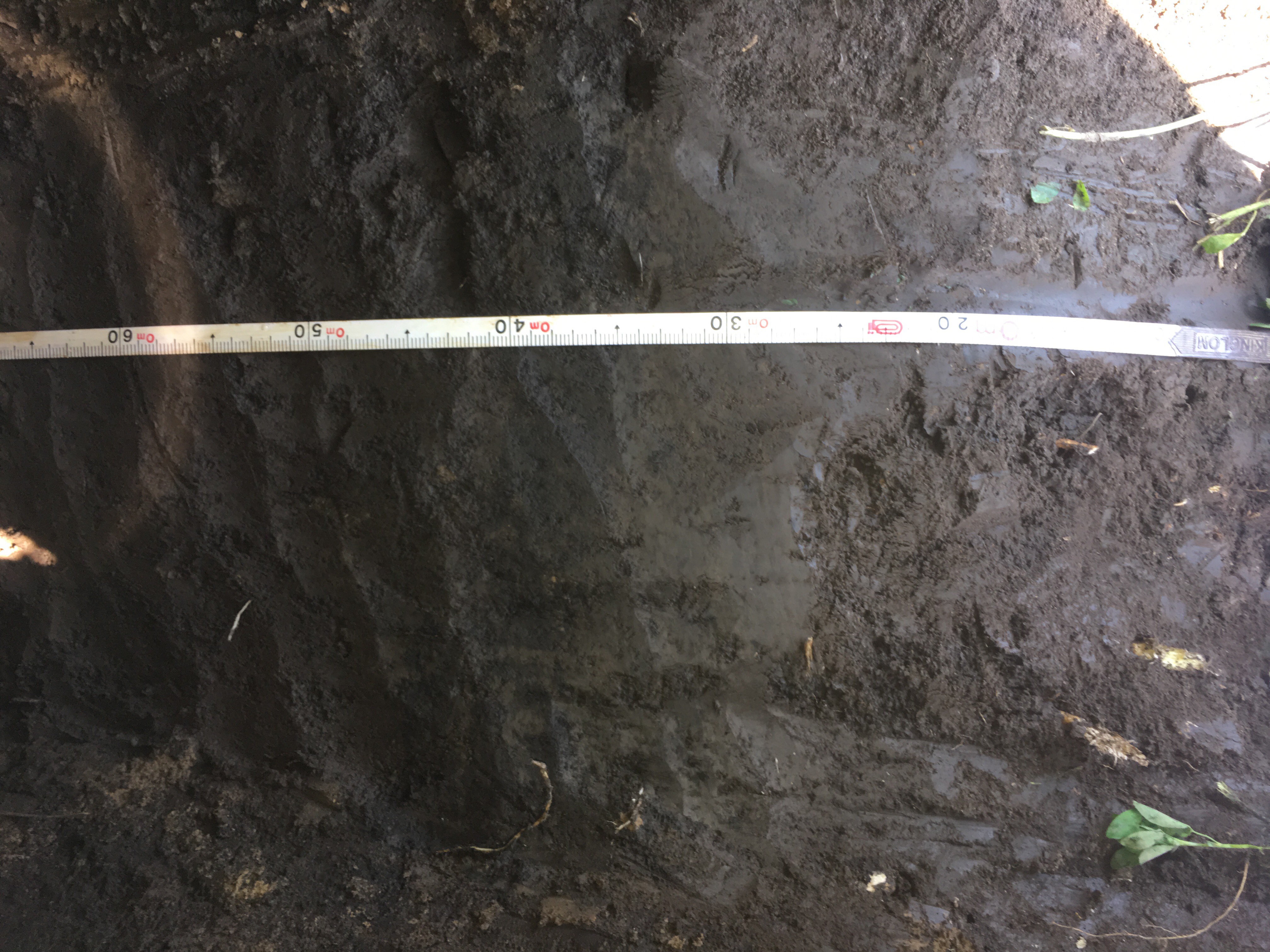

### a341-60cm-80cm.jpg

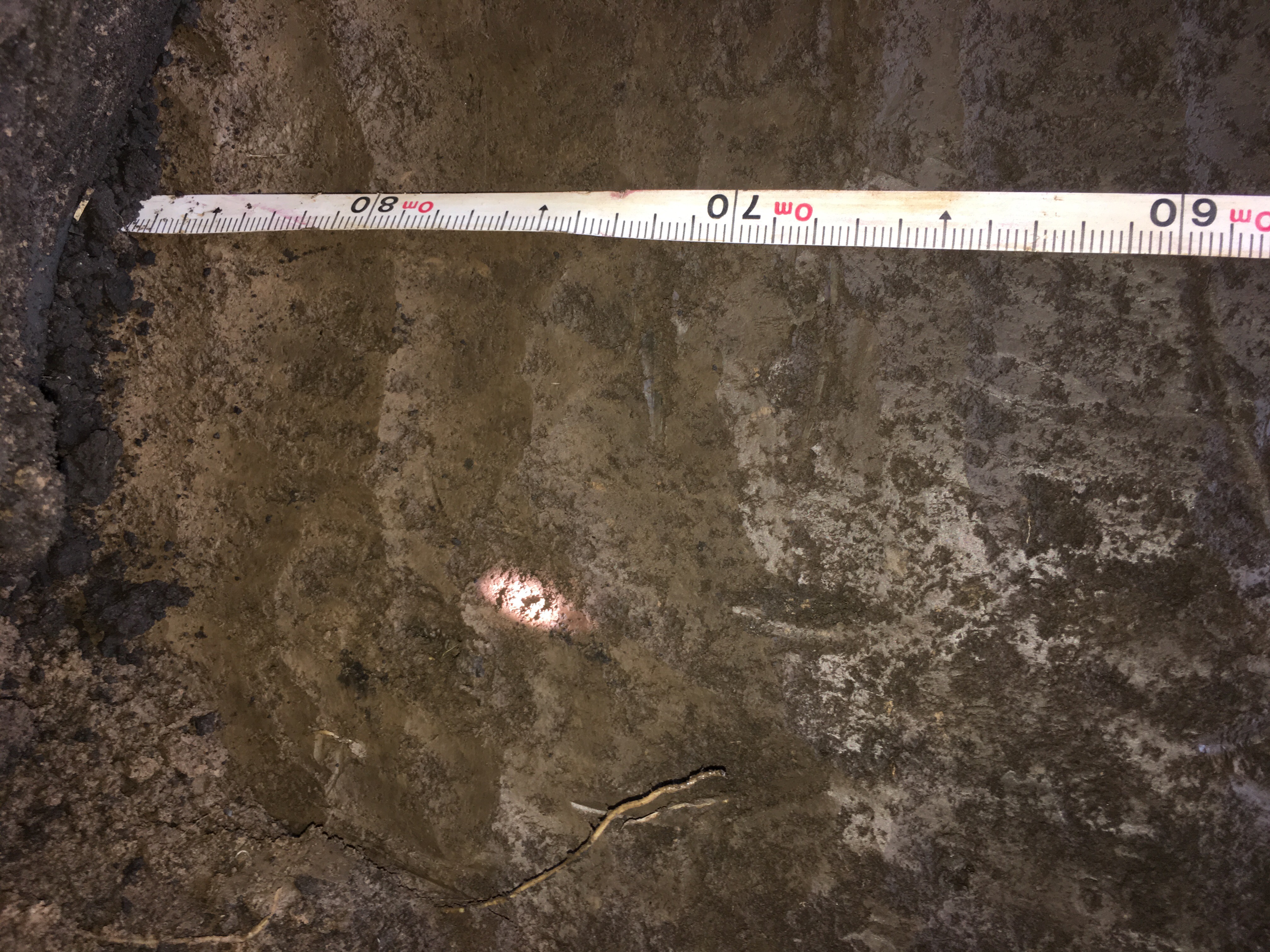

### h1-0cm-70cm.jpg

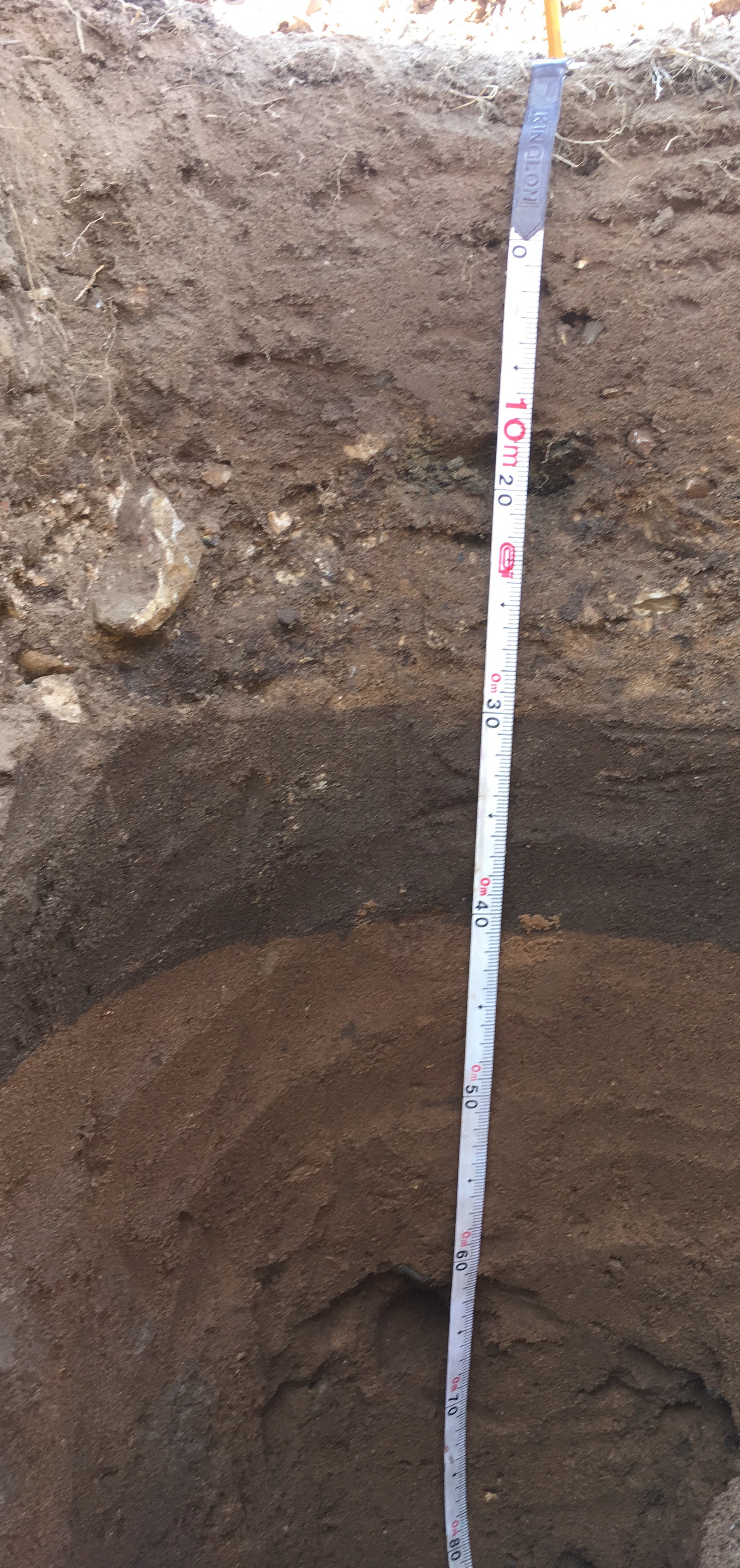

### h2-0cm-70cm.jpg

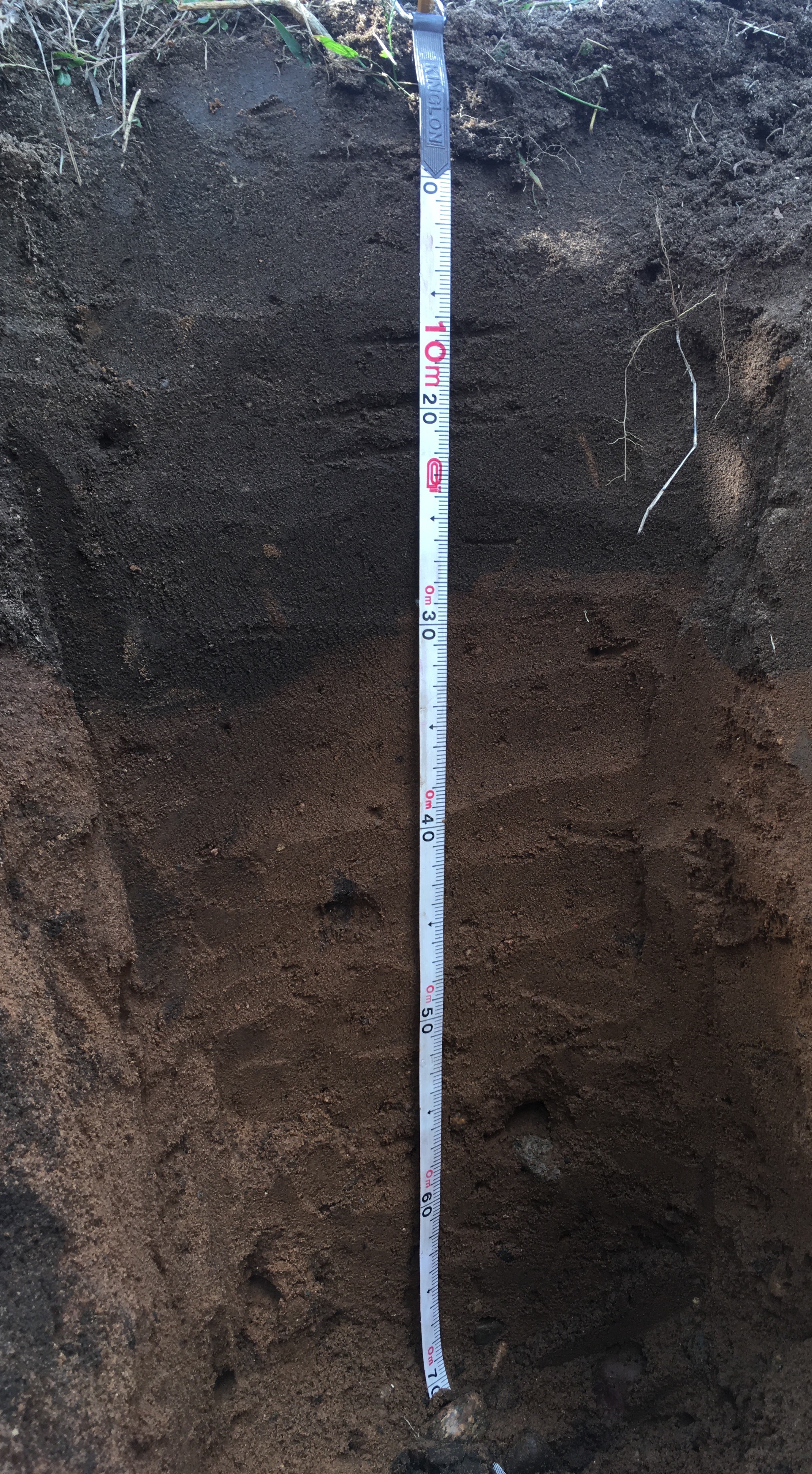

### k1-0cm-60cm.jpg

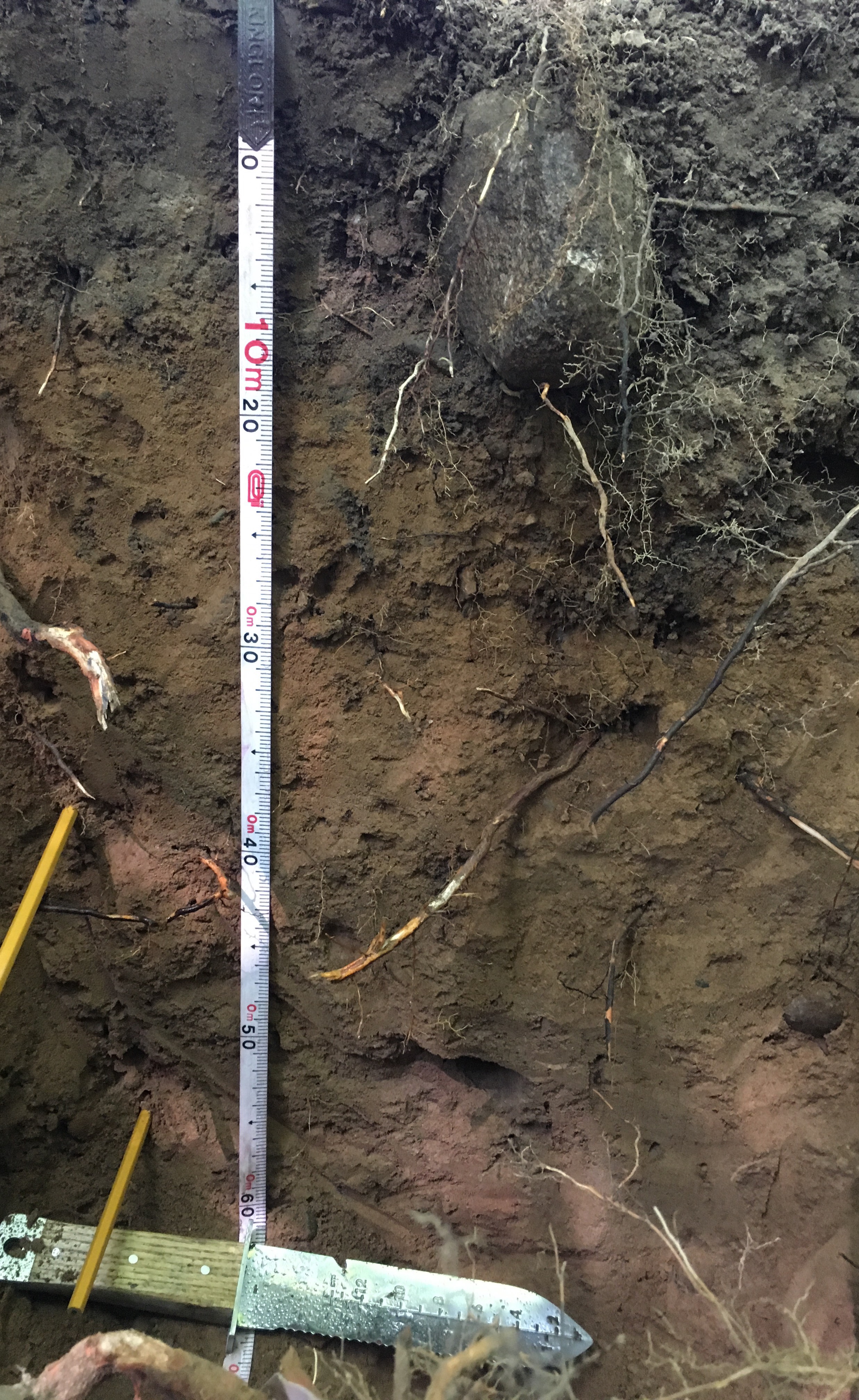

### k3-0cm-40cm.jpg

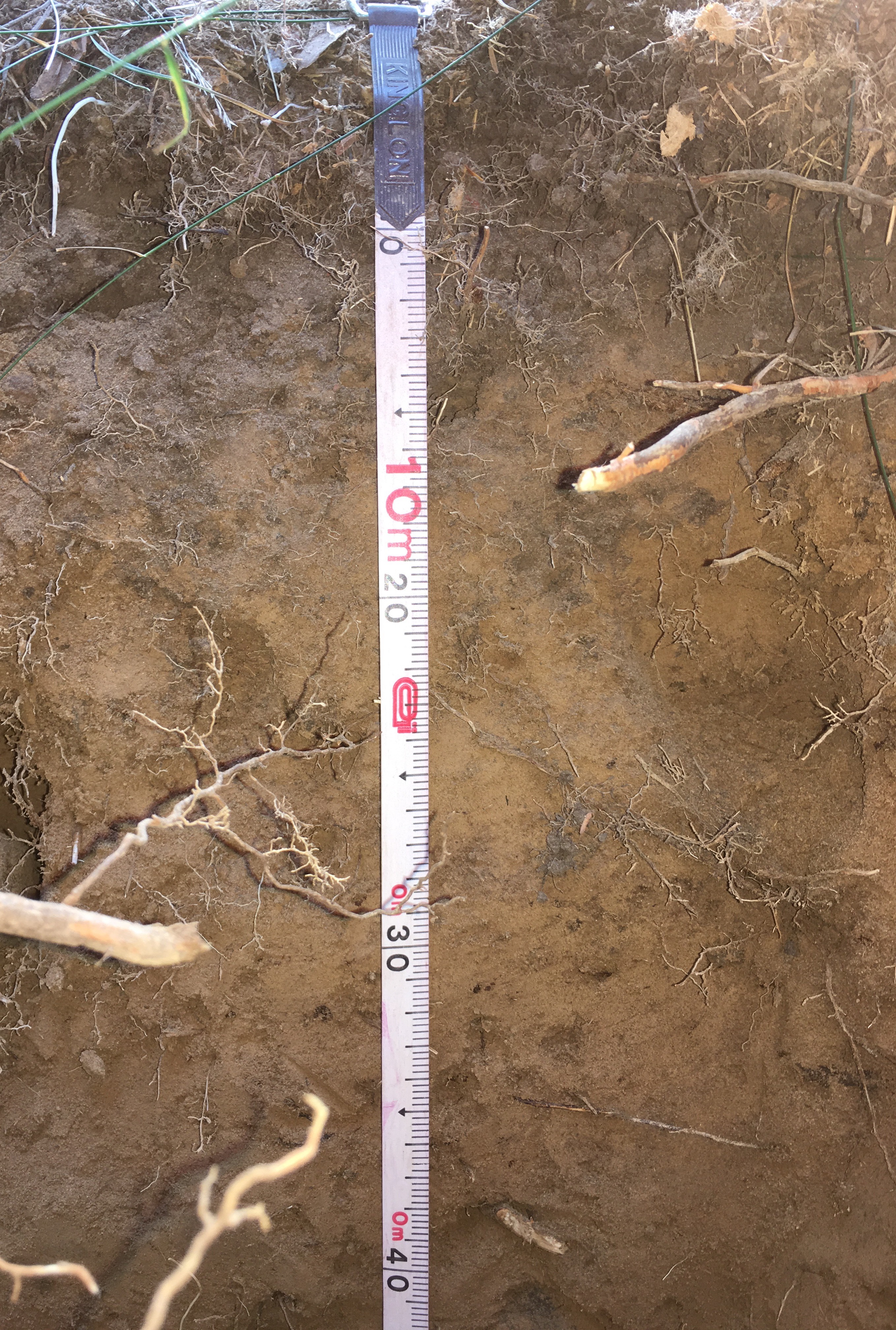

### k4-0cm-60cm.jpg

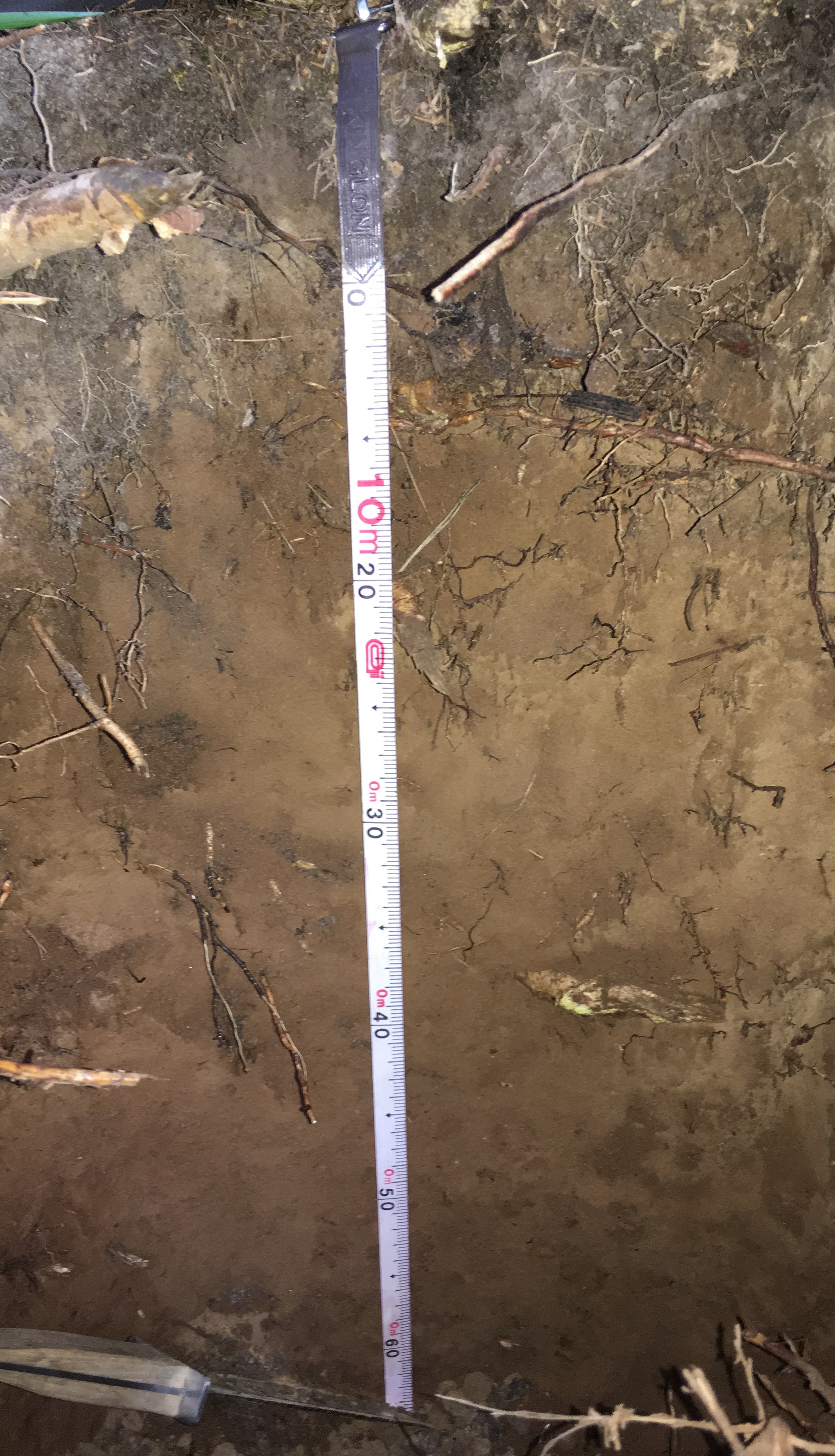

### l2-0cm-40cm.jpg

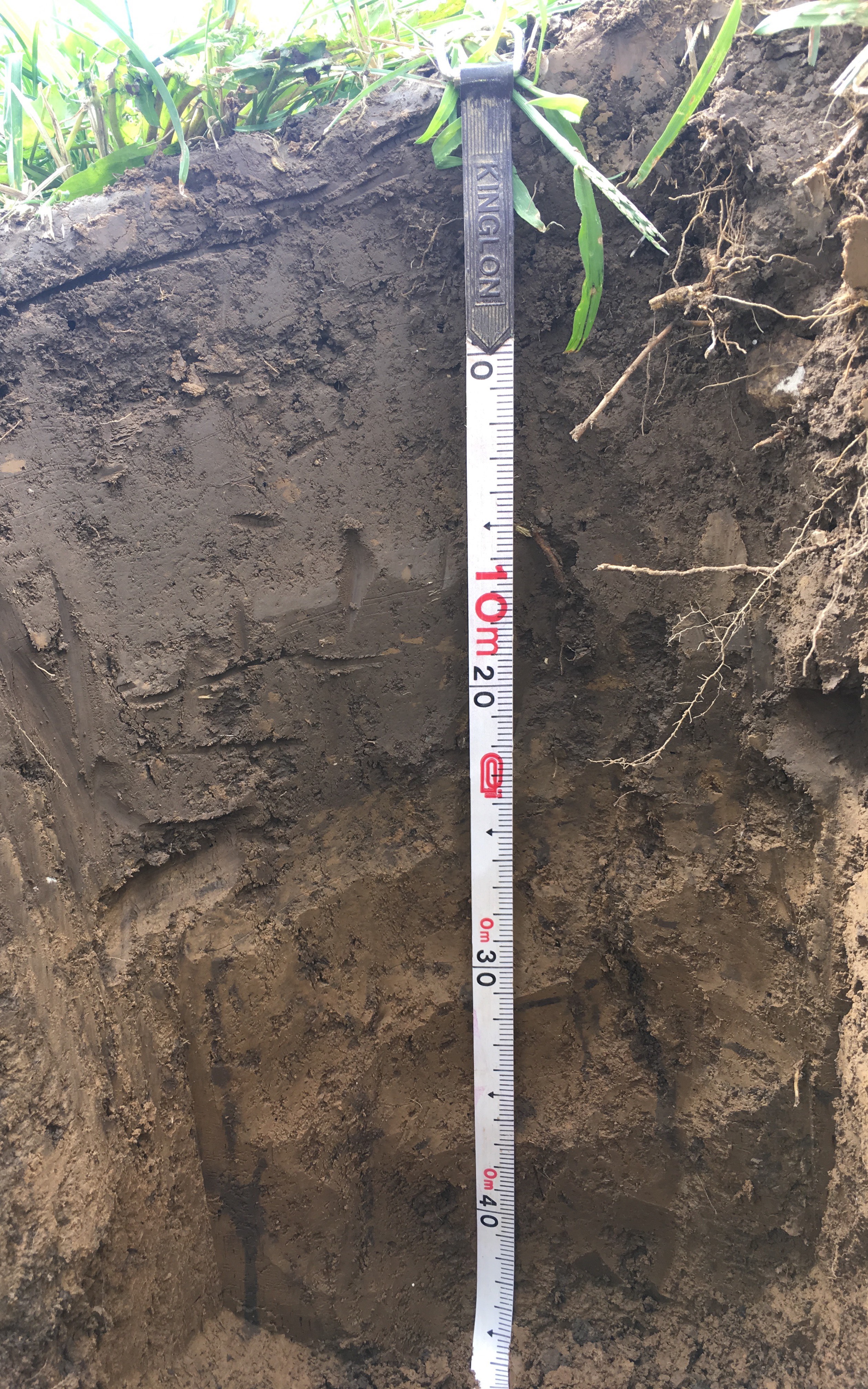

### l3-0cm-50cm.jpg

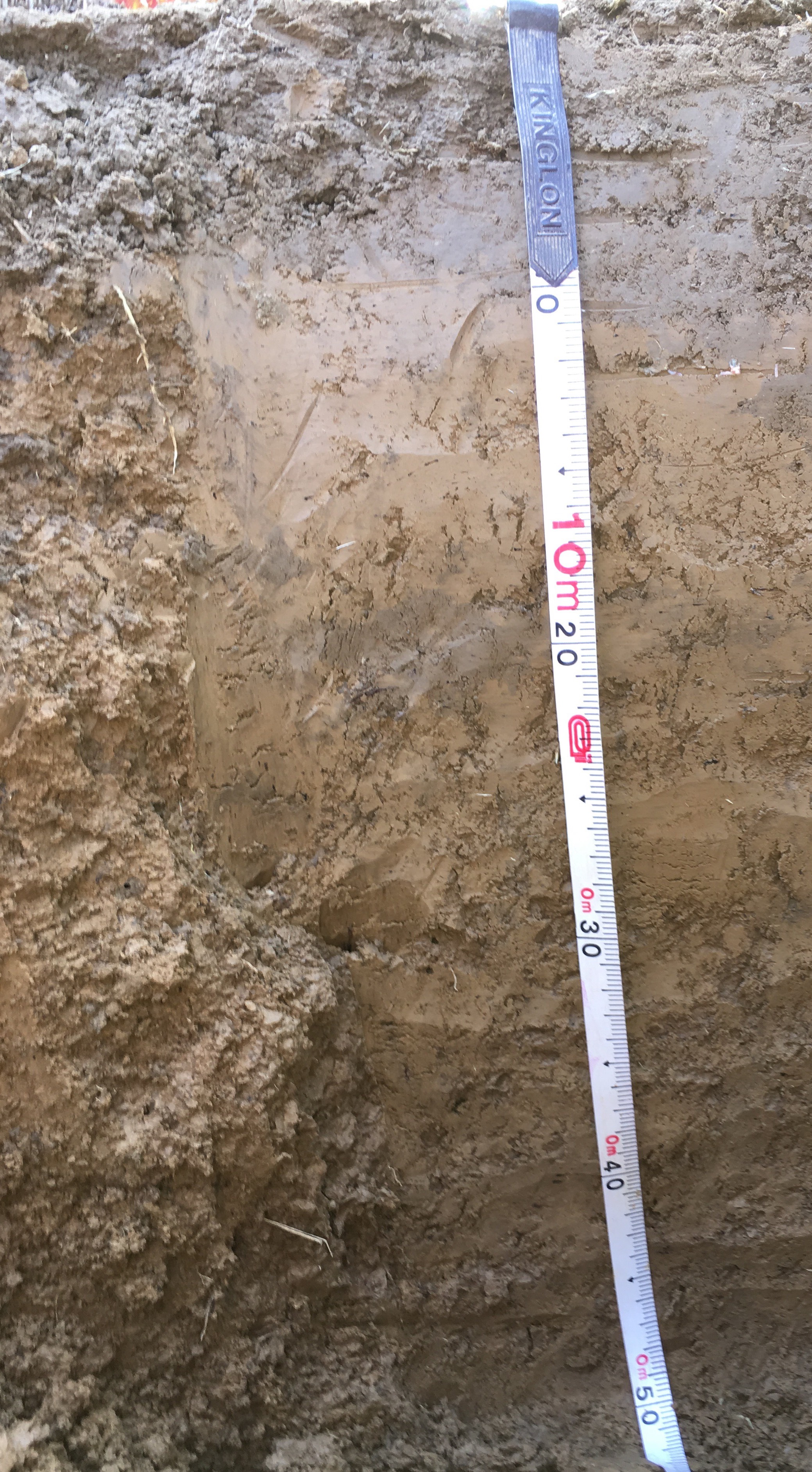

### l4-0cm-50cm.jpg

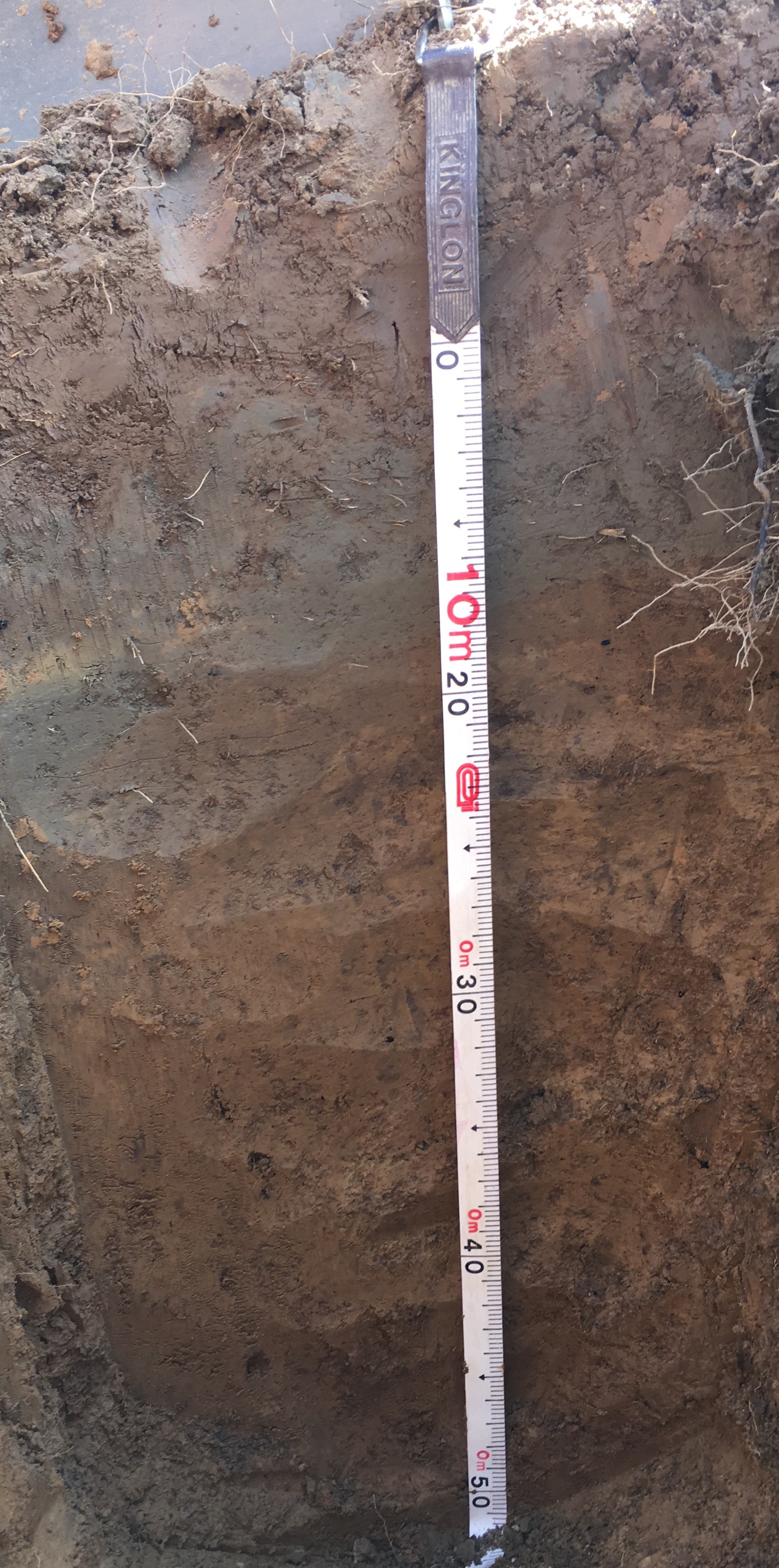

### m1-0cm-75cm.jpg

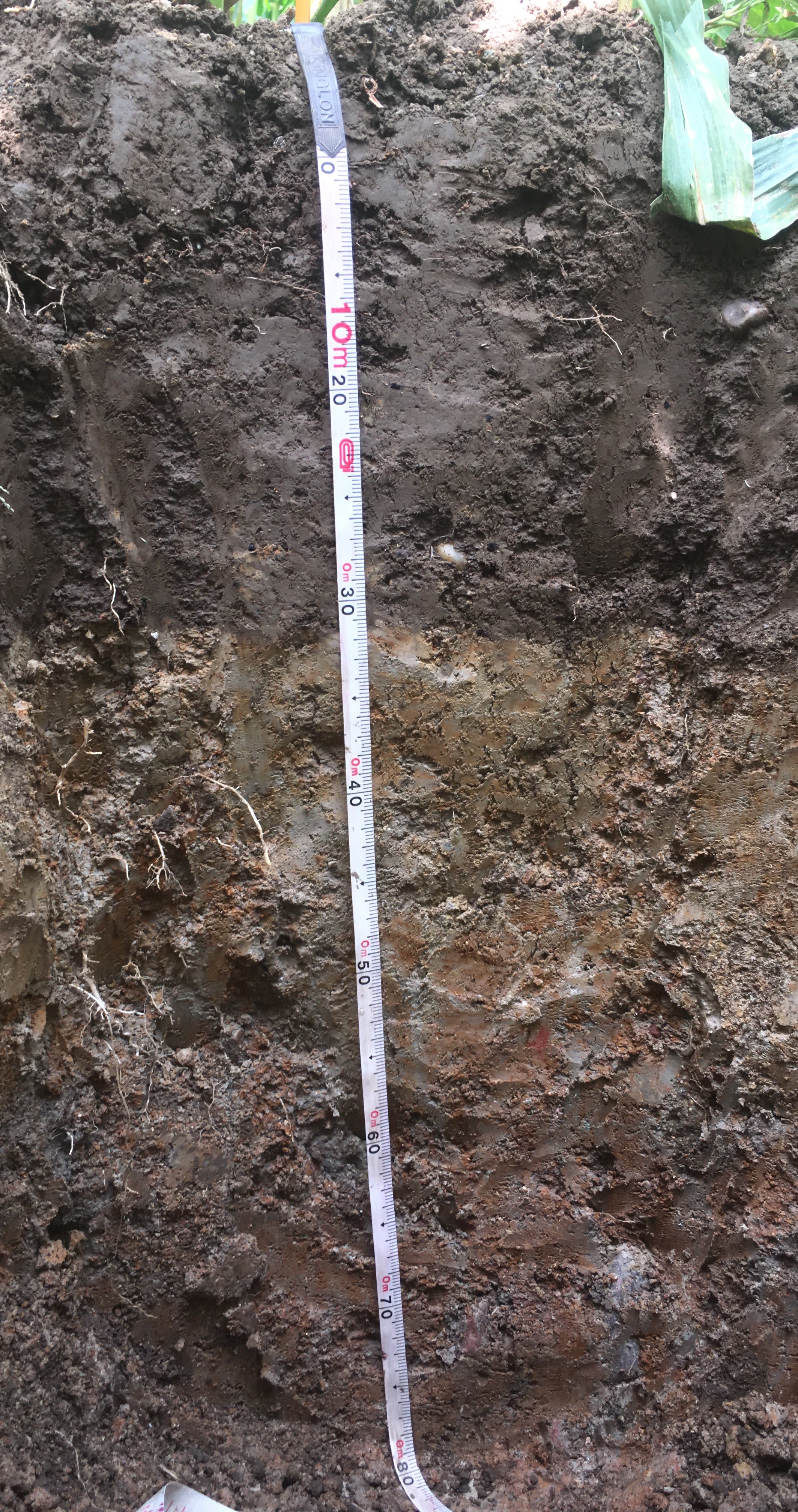

### m2-0cm-60cm.jpg

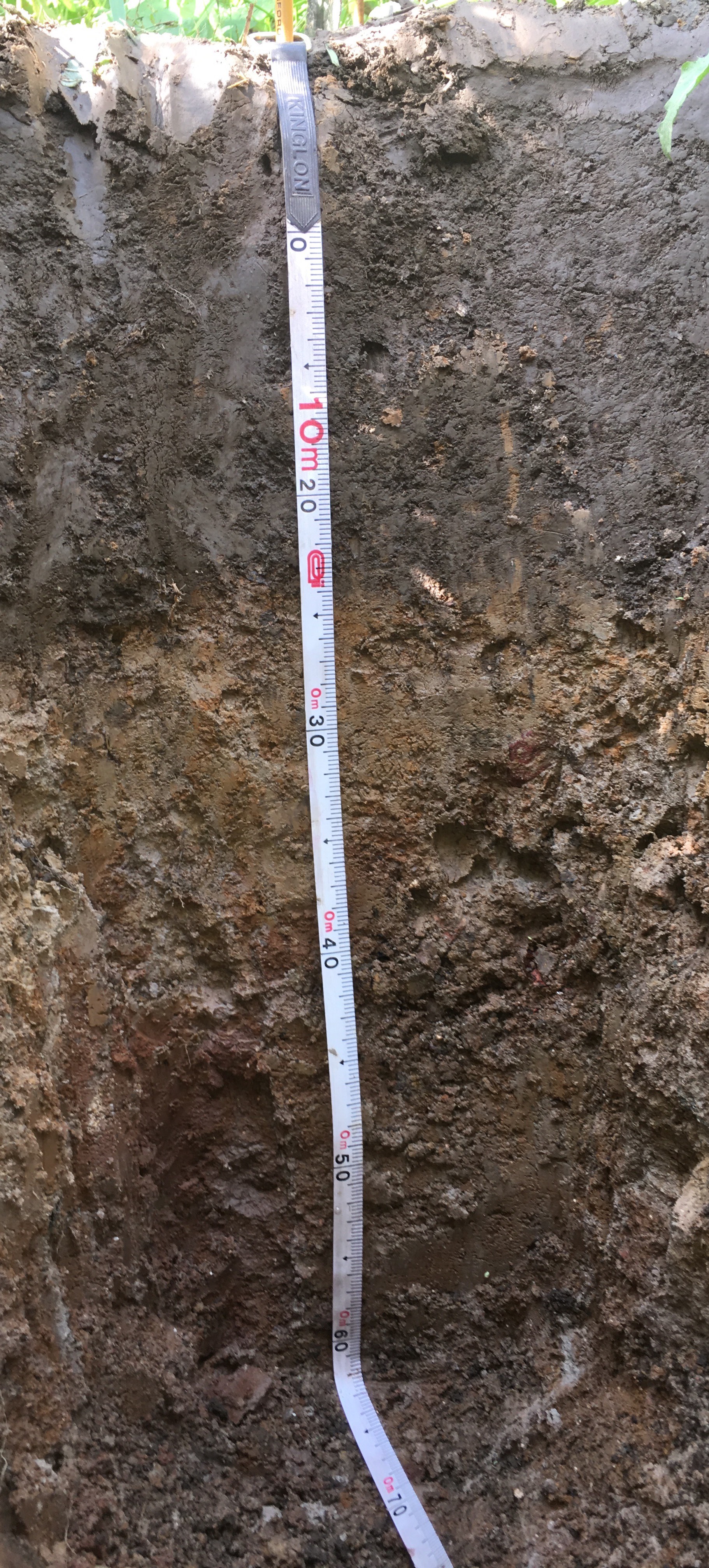

### m3-0cm-40cm.jpg

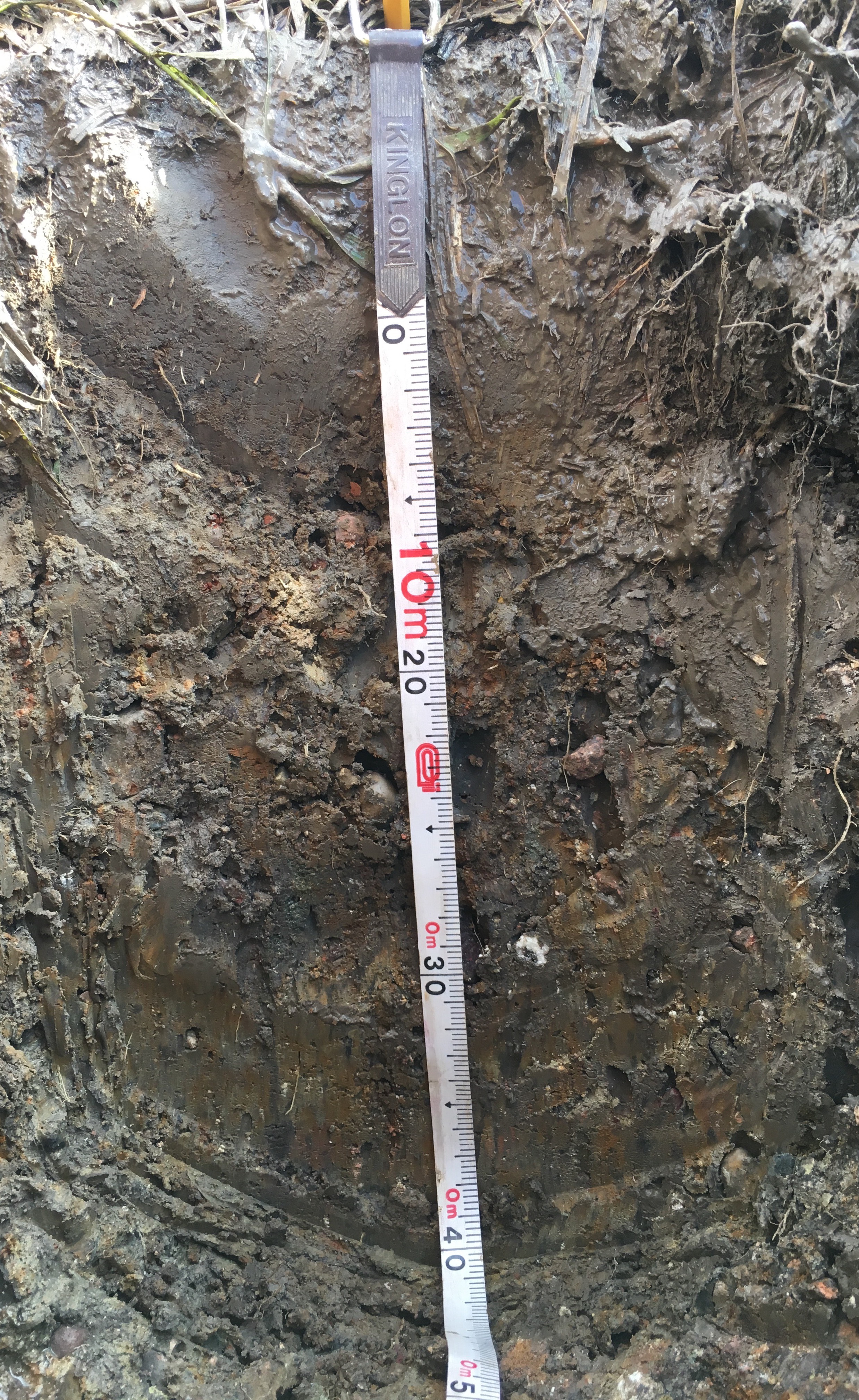

### p1-0cm-60cm.jpg

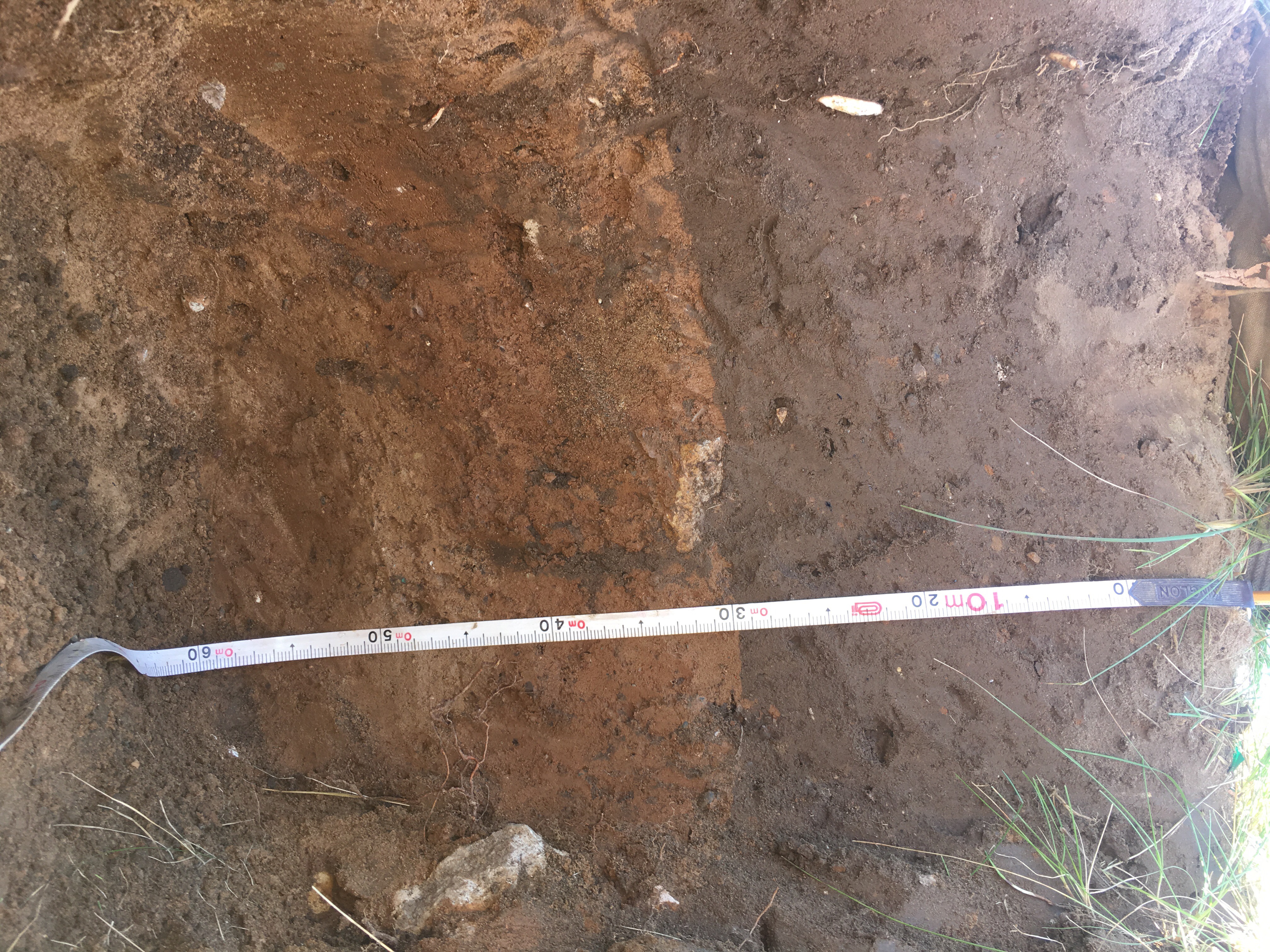

### p2-0cm-50cm.jpg

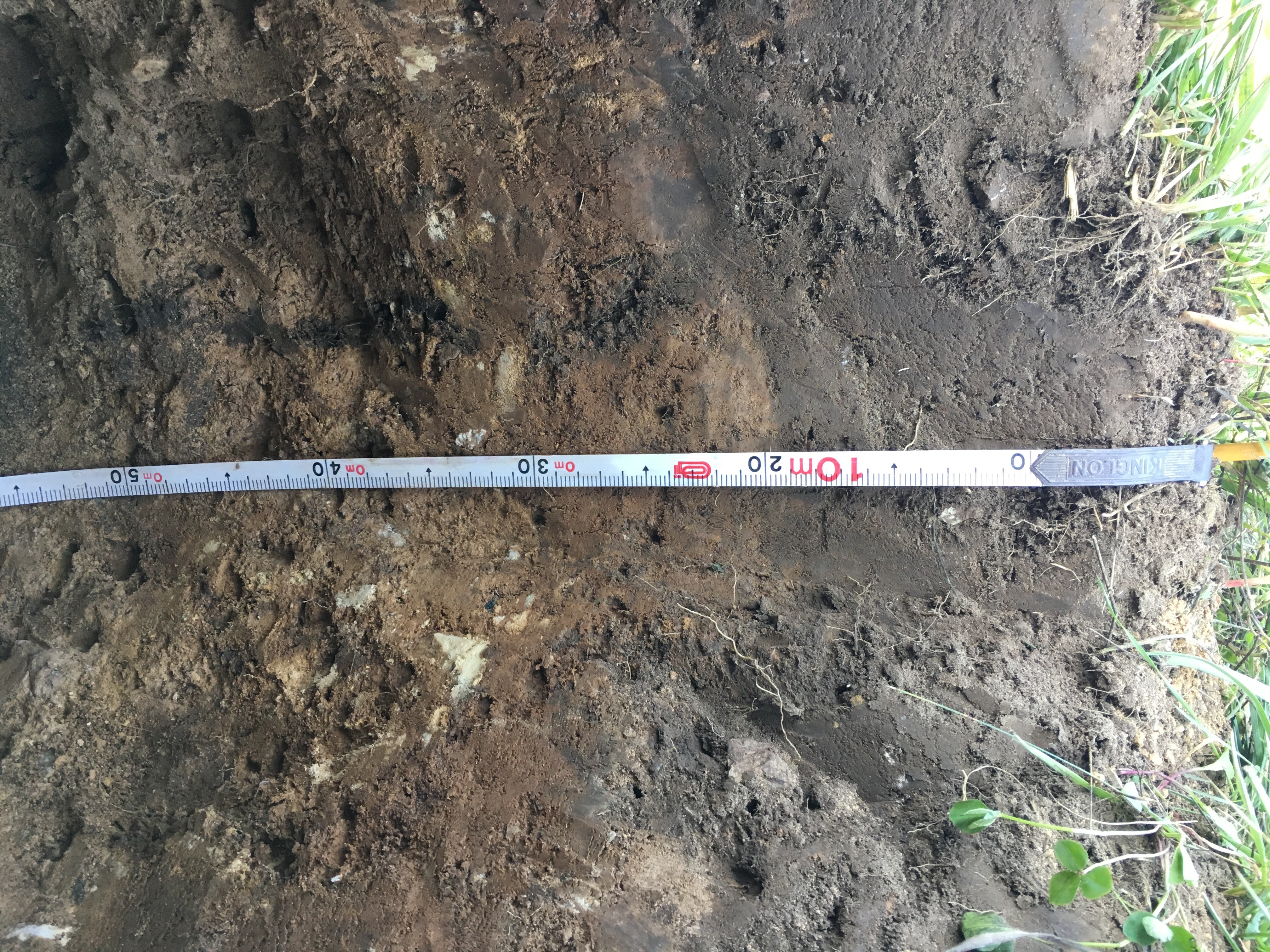

### p4-0cm-50cm.jpg

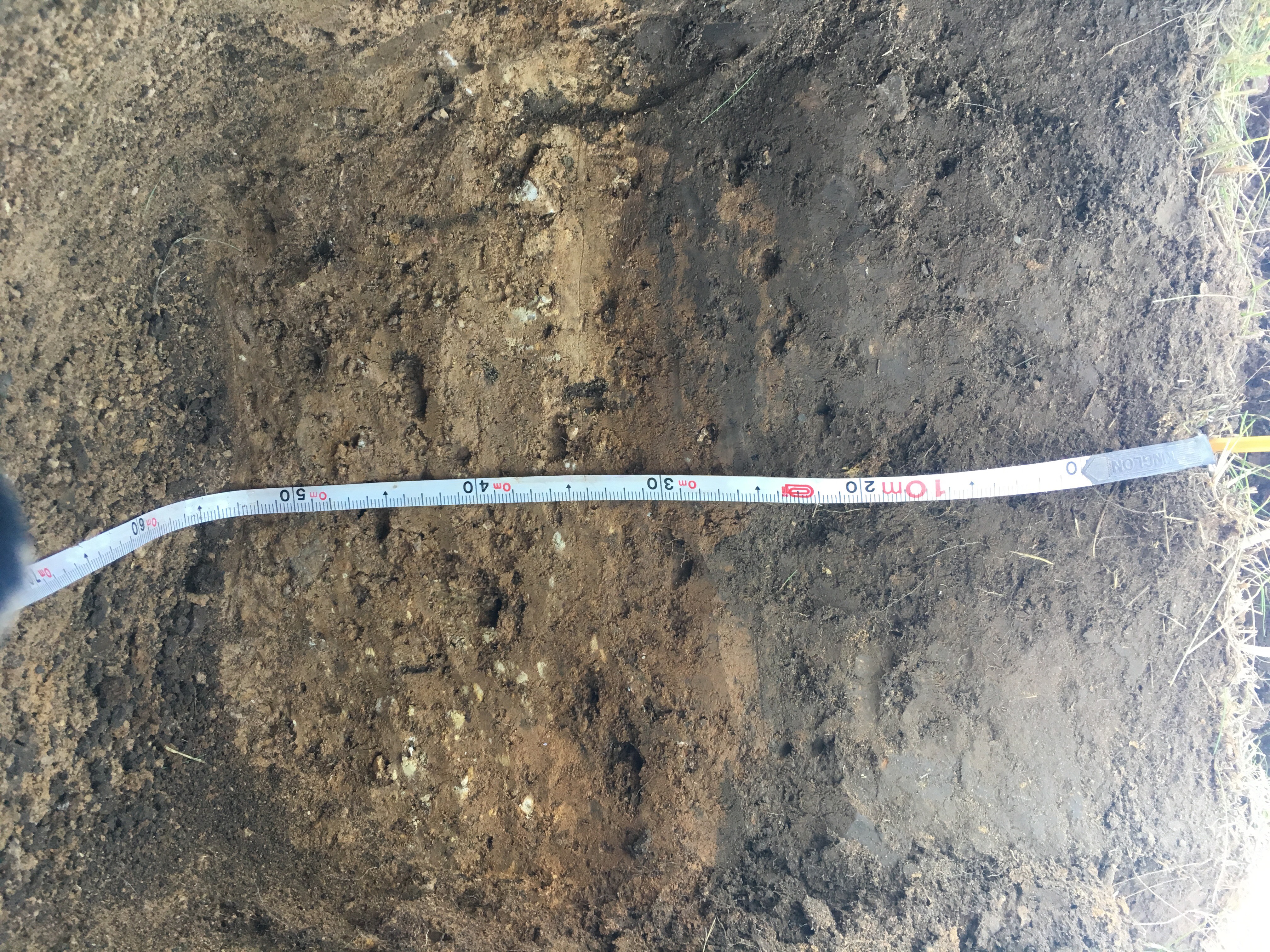

### r1-0cm-50cm.jpg

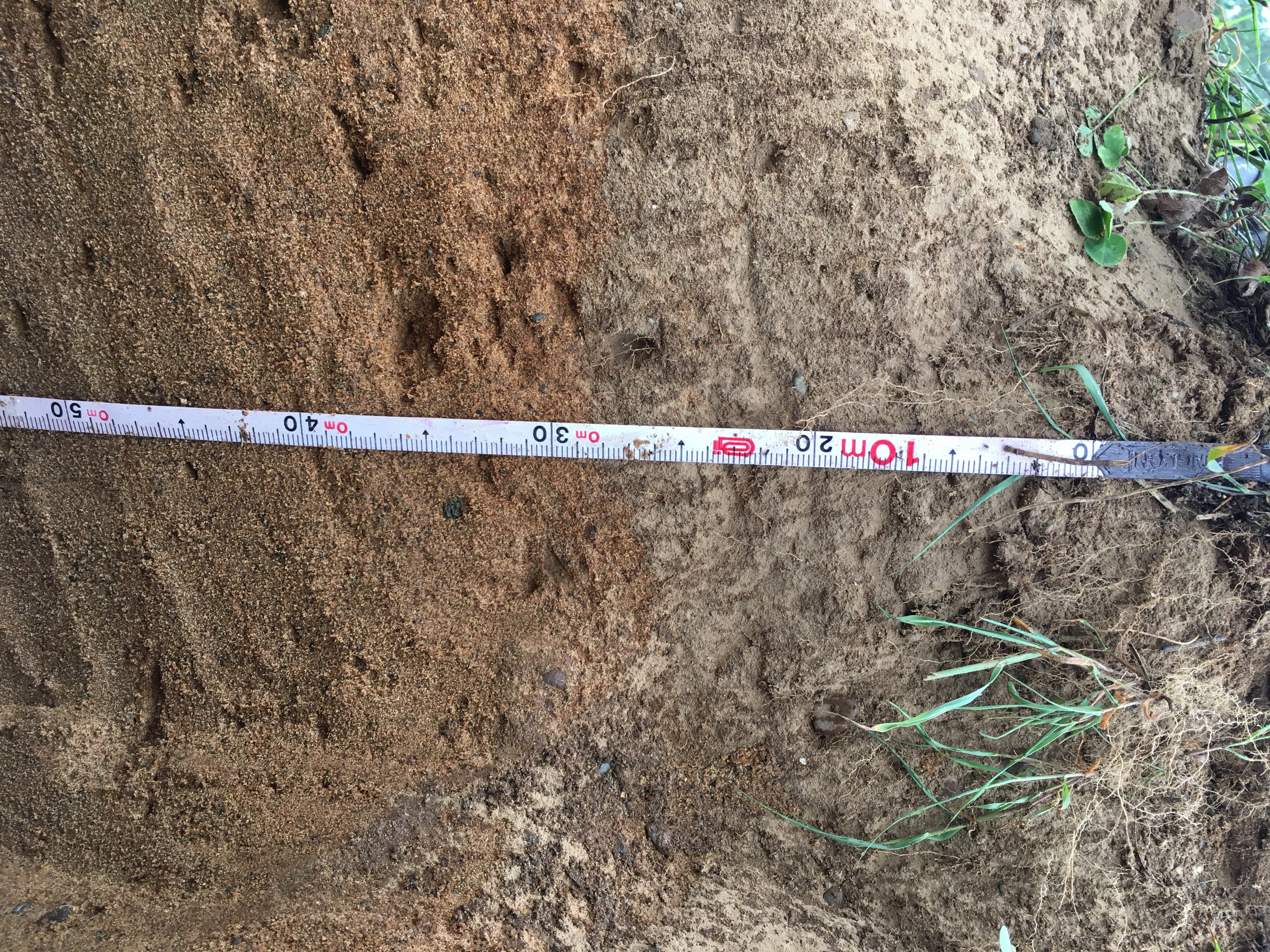

### r2-0cm-170cm.jpg

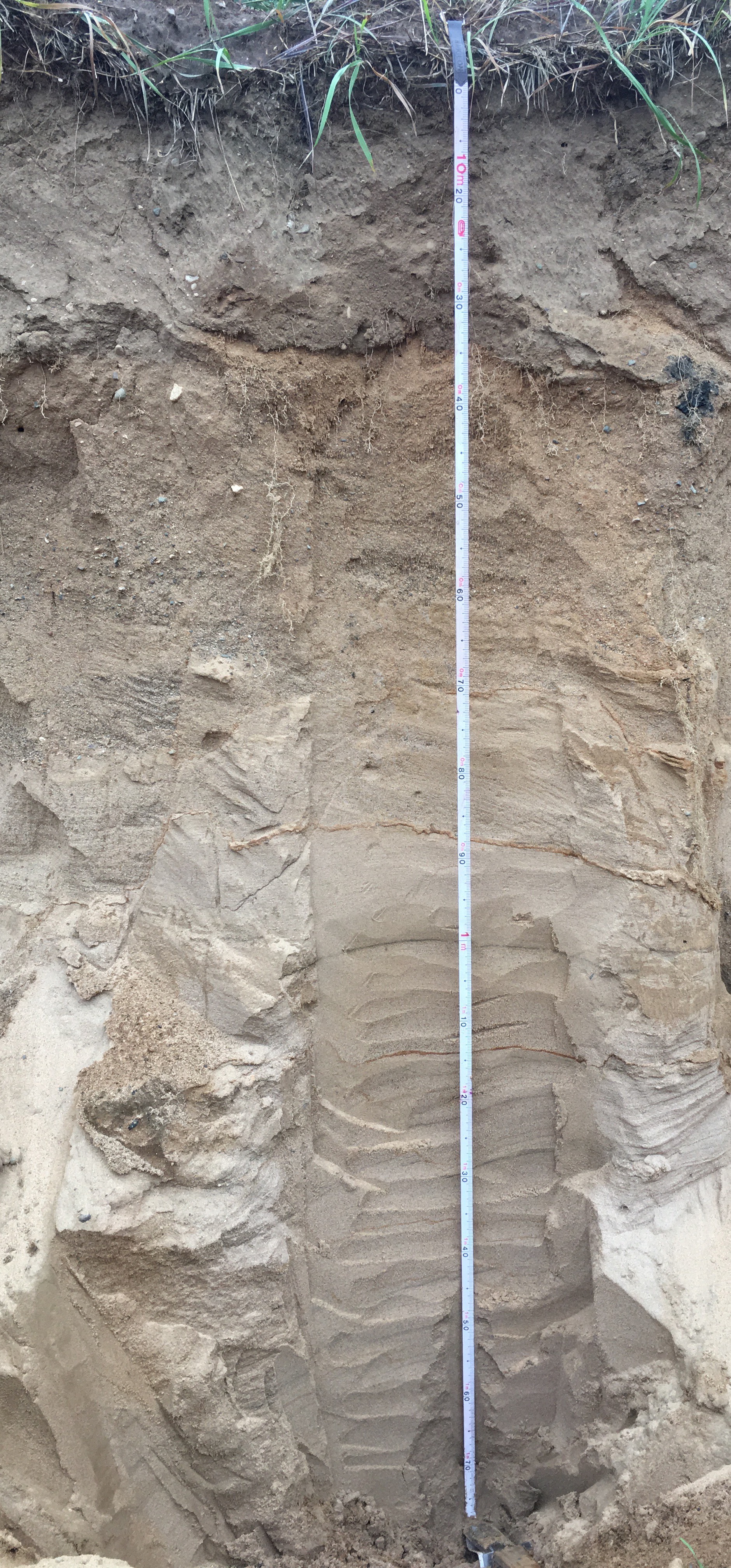

### r3-0cm-55cm.jpg

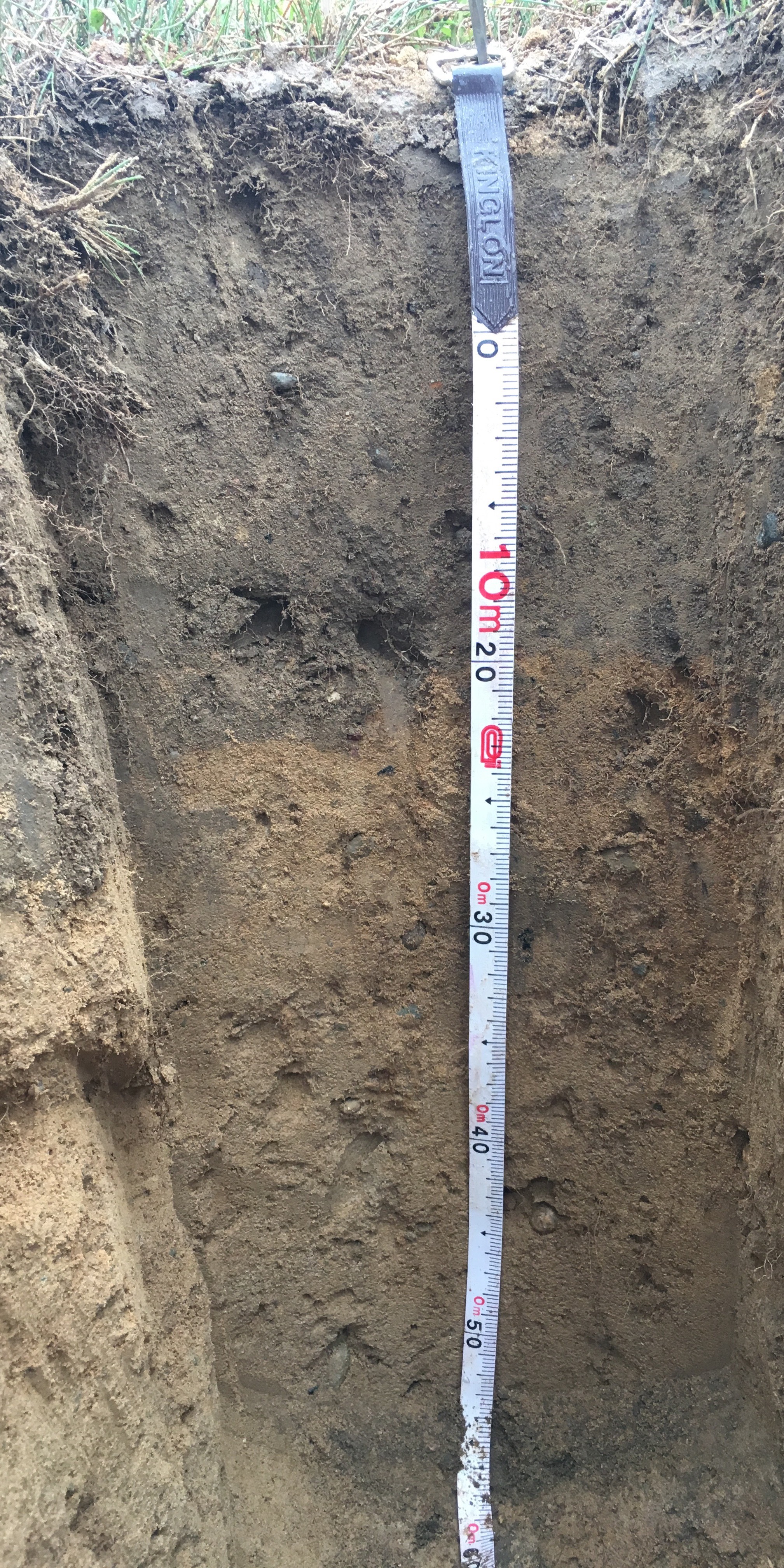

### sp-0cm-75cm.jpg

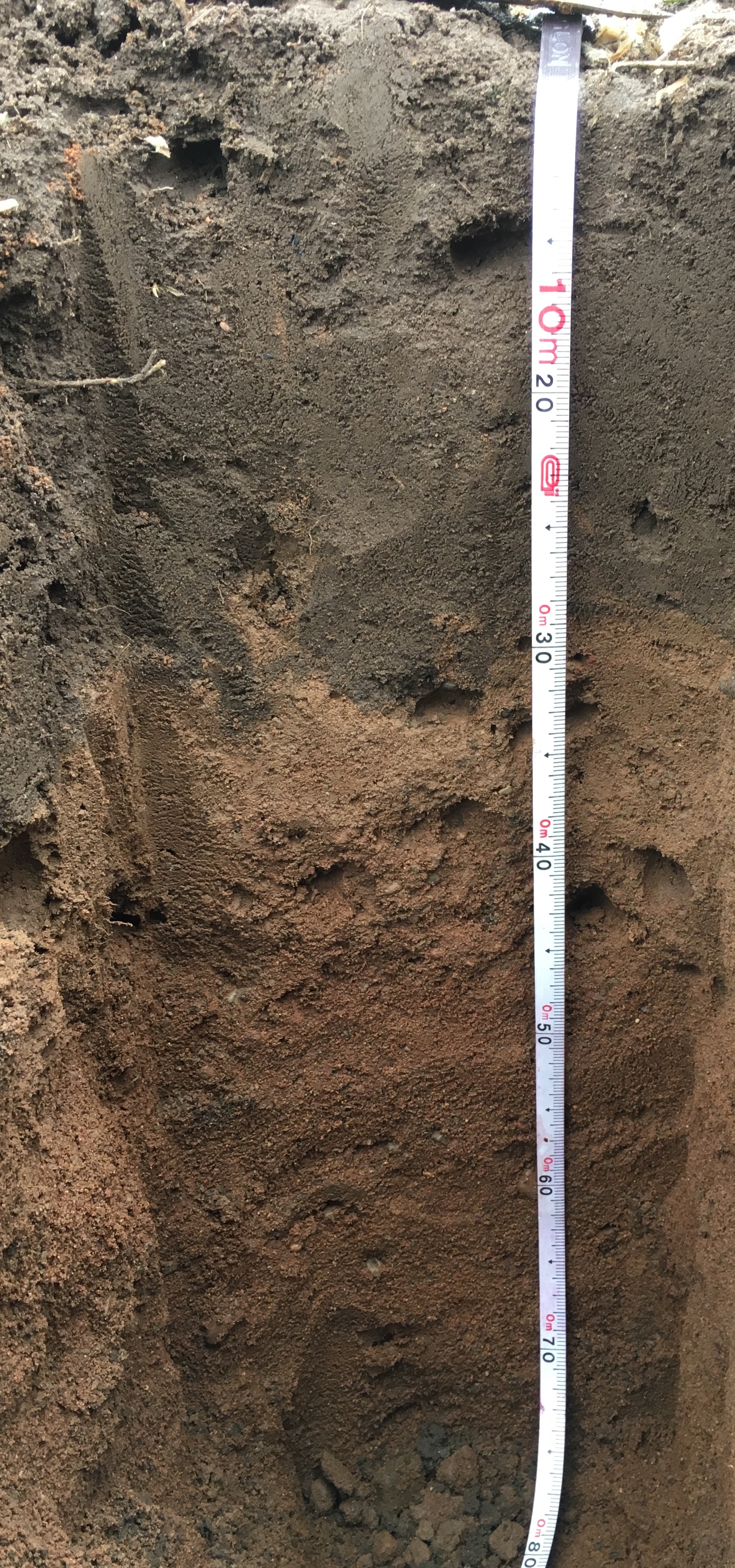

### supplementary-figure-1.png

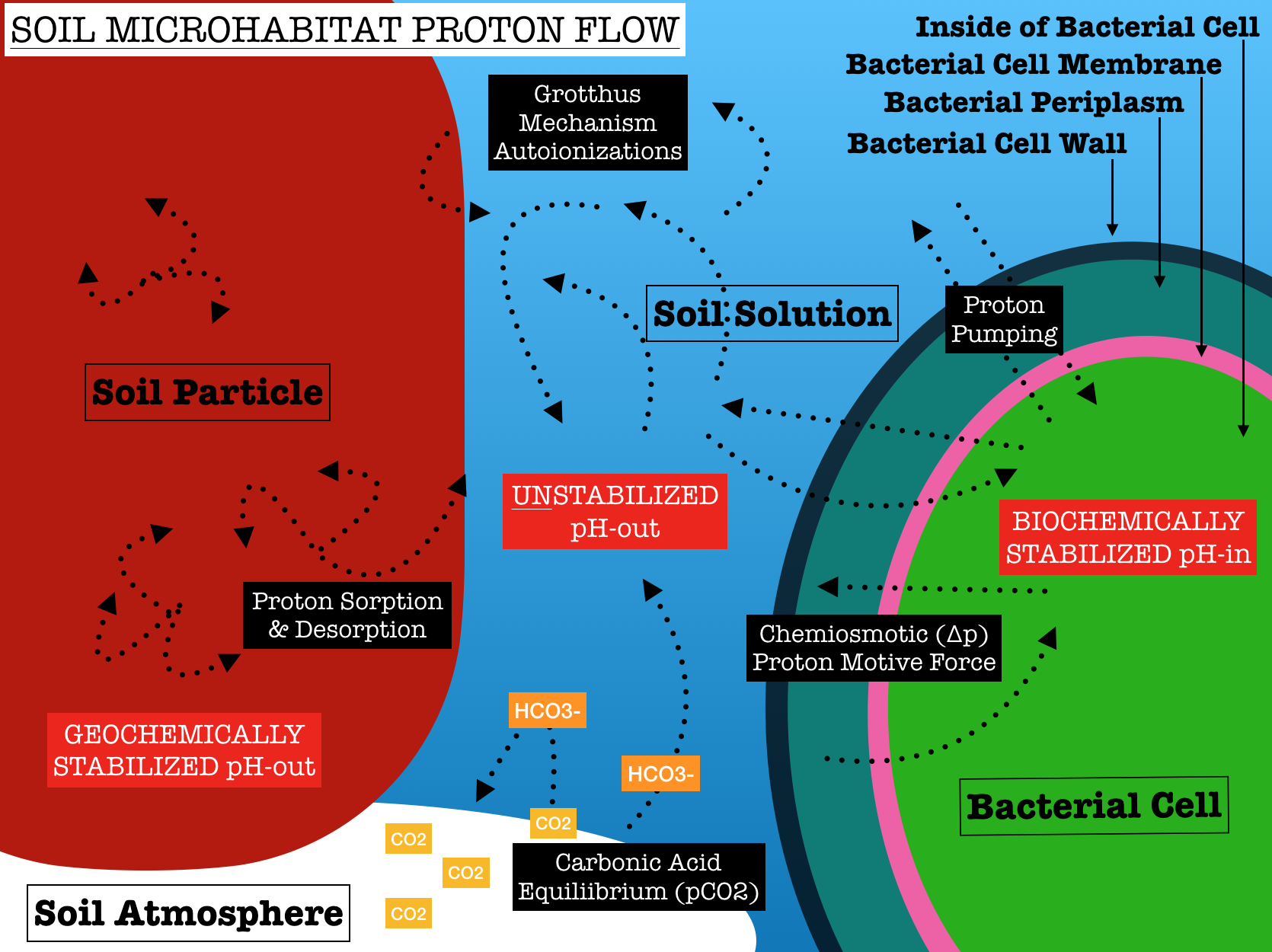

### supplementary-figure-2.png

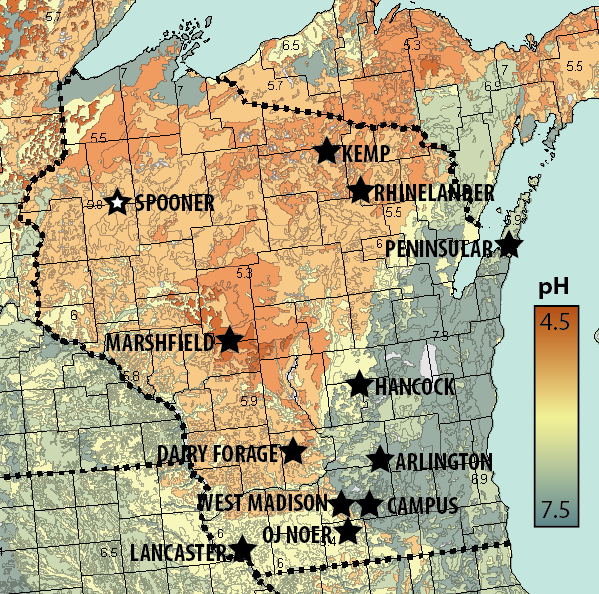

### supplementary-figure-3.jpg

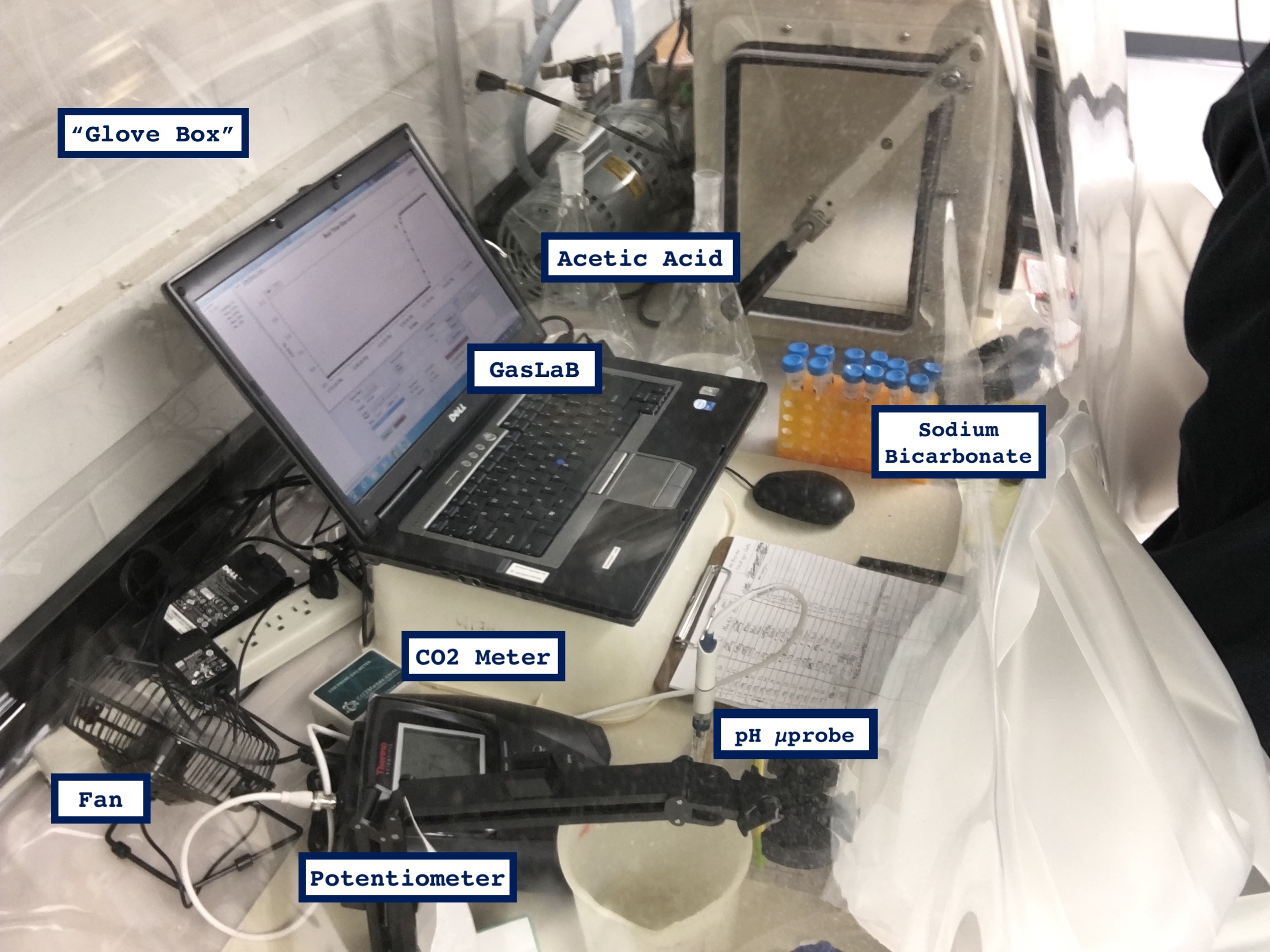

### supplementary-figure-4.png

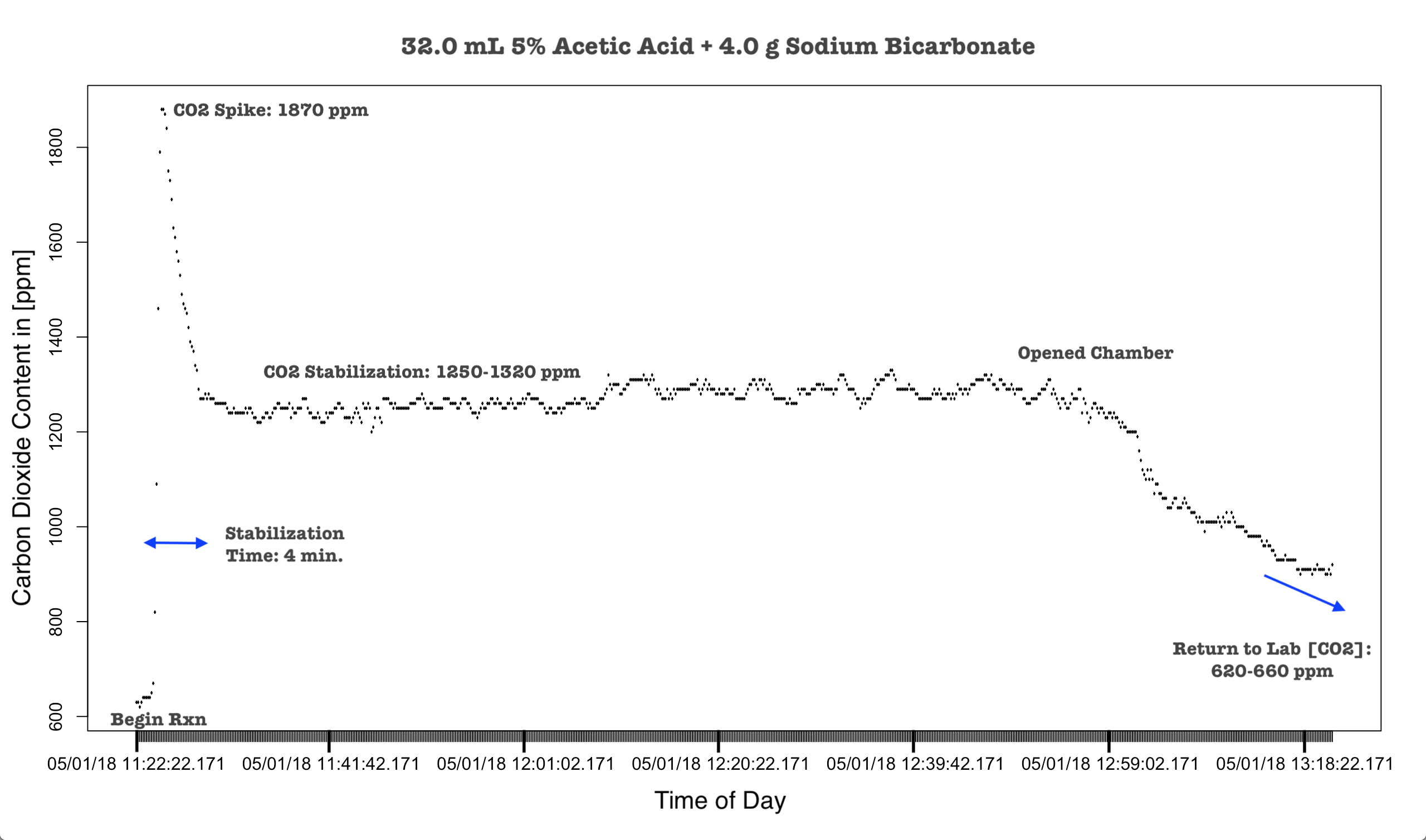

### supplementary-figure-5.png

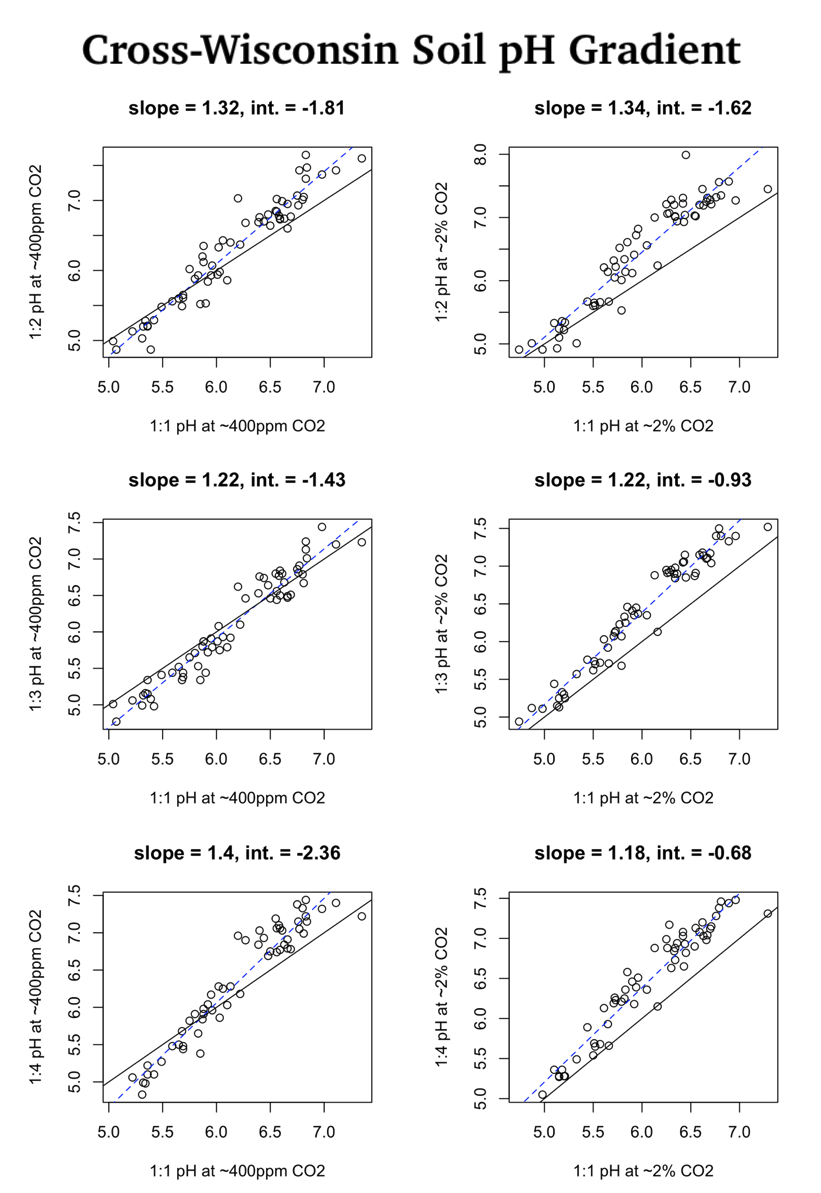

### supplementary-figure-6.png

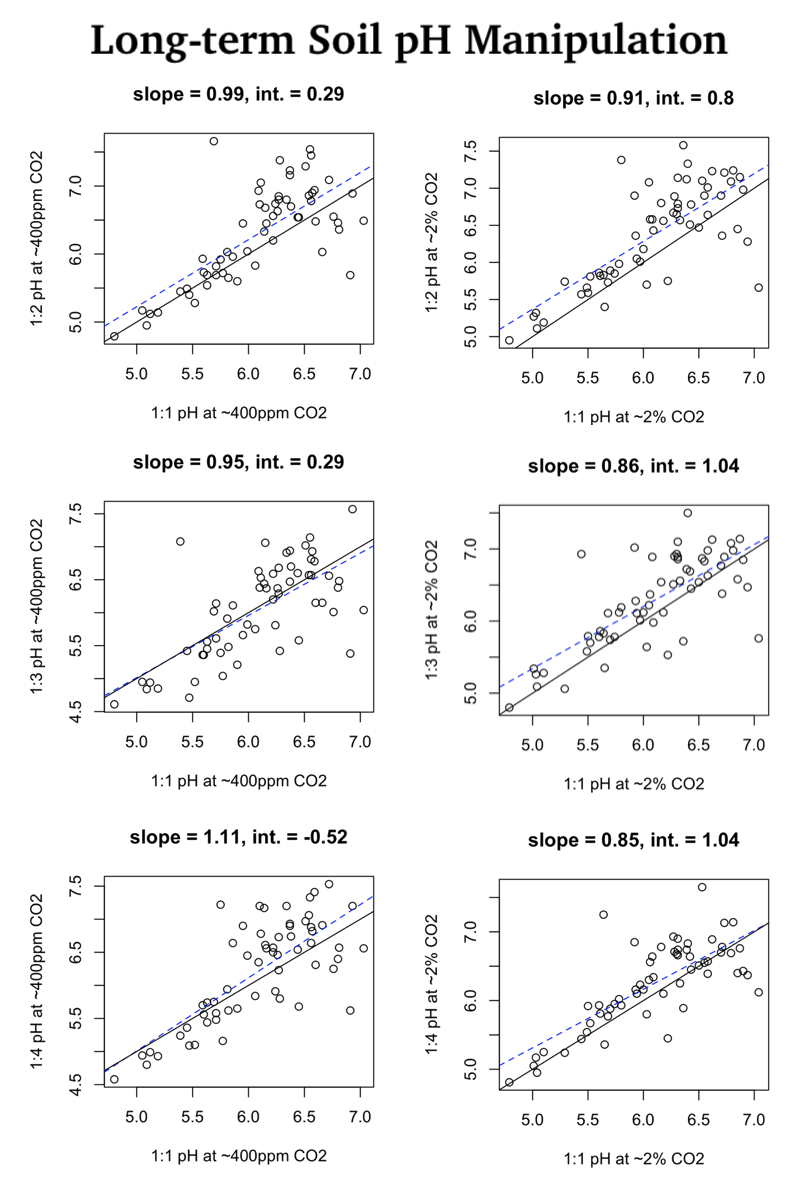

### supplementary-figure-7.png

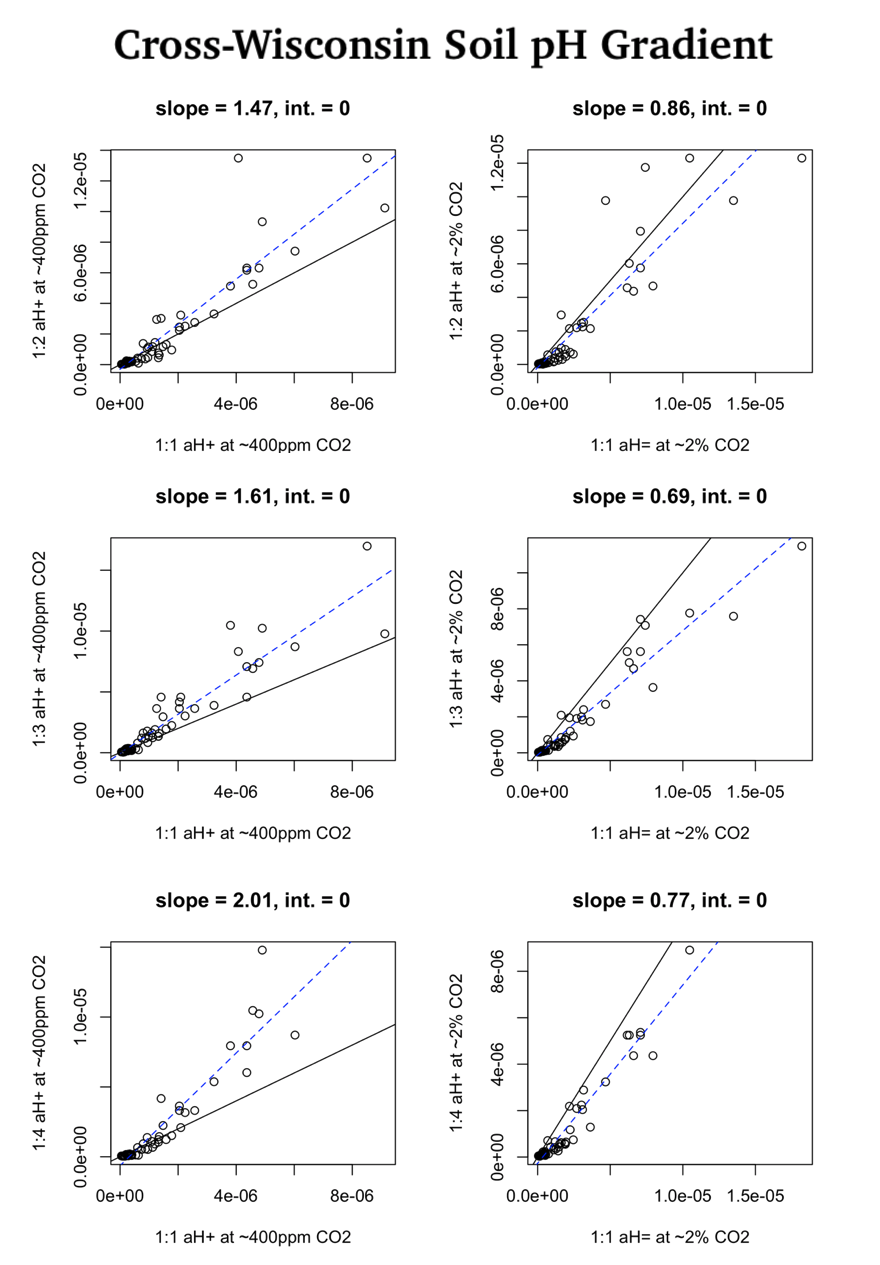

### supplementary-figure-8.png

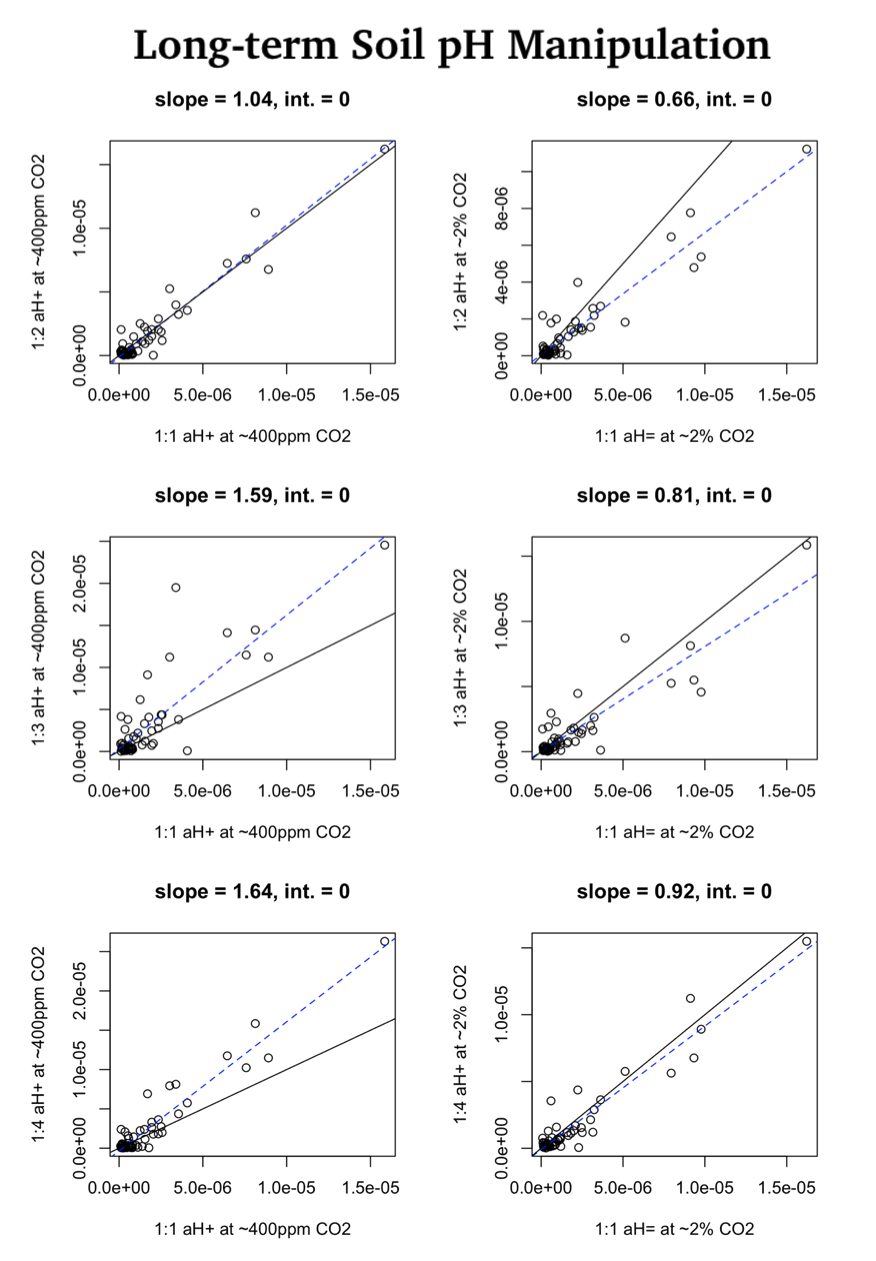

### supplementary-figure-9.png

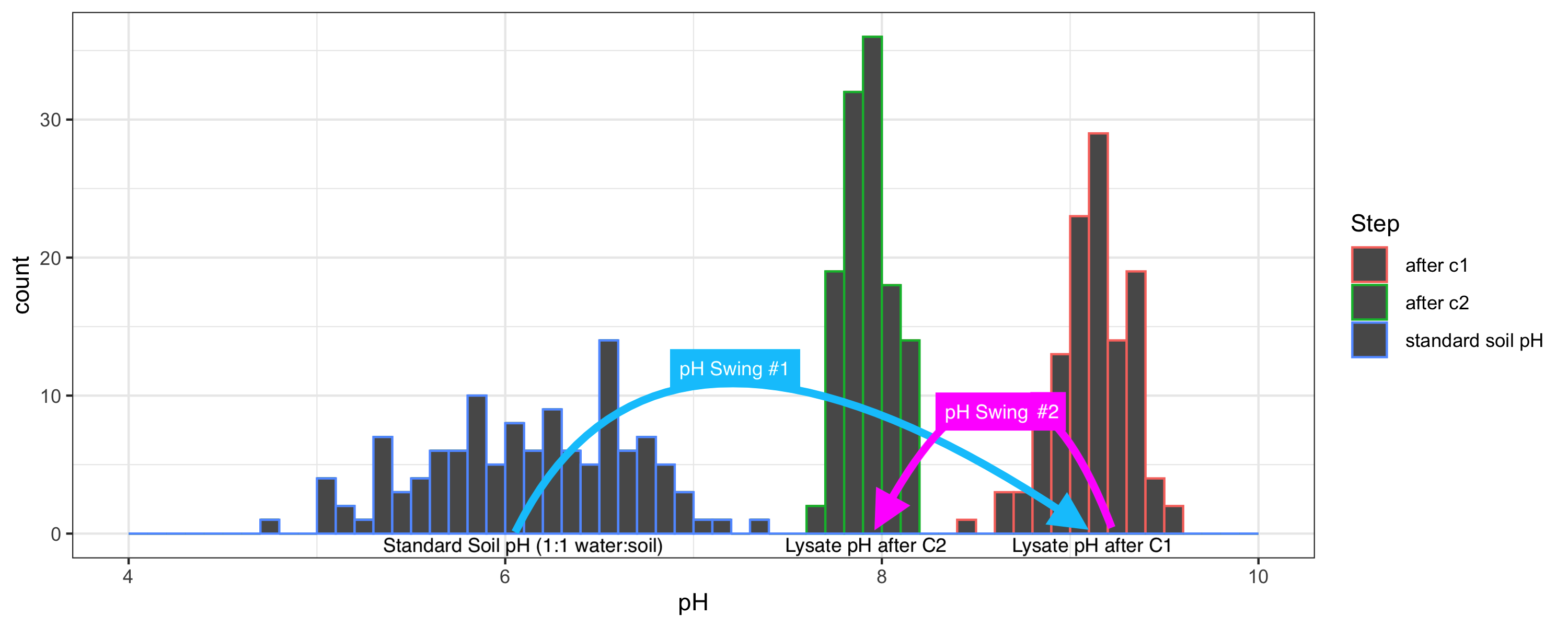

### supplementary-figure-10.png

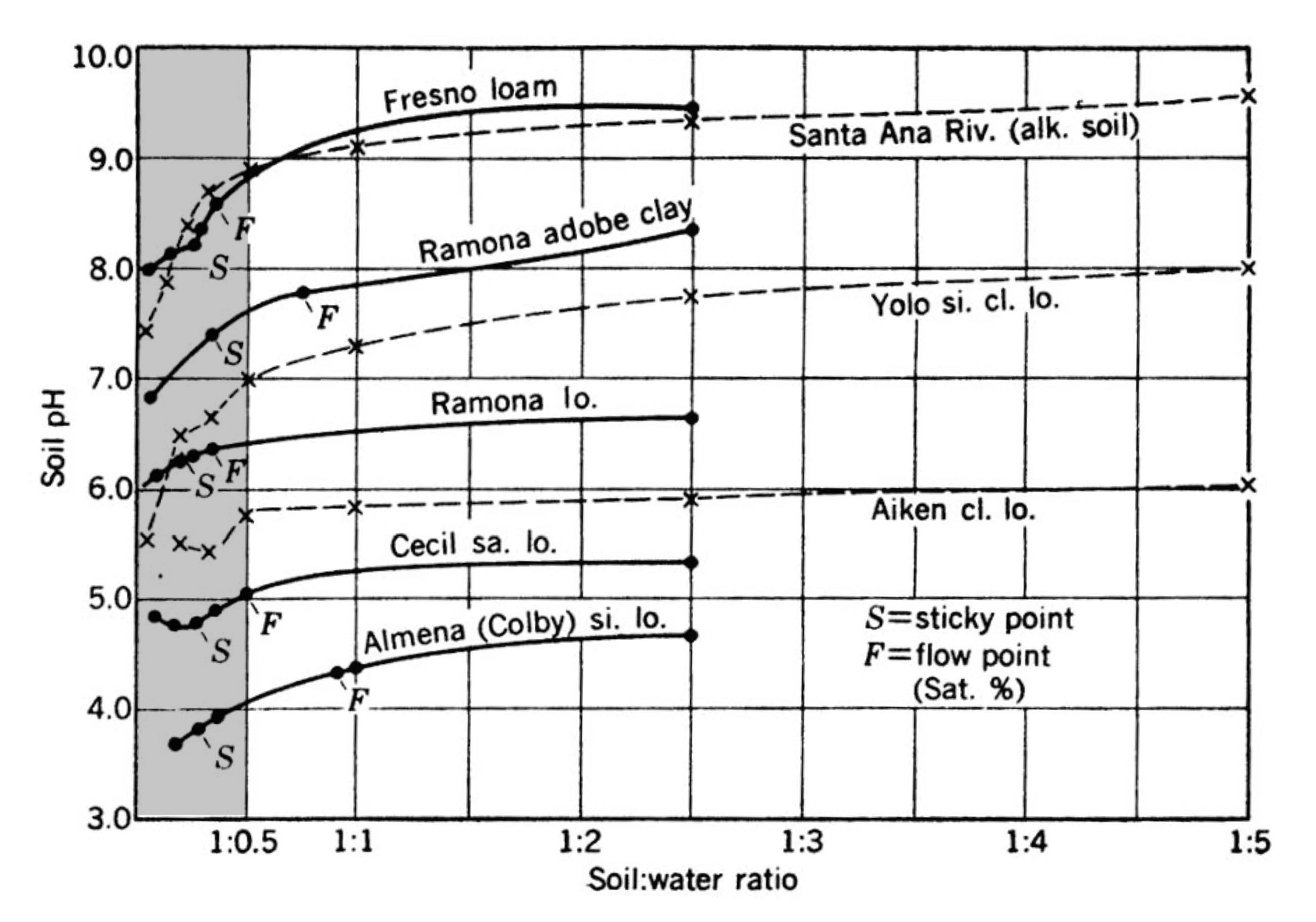

### supplementary-table-2.png

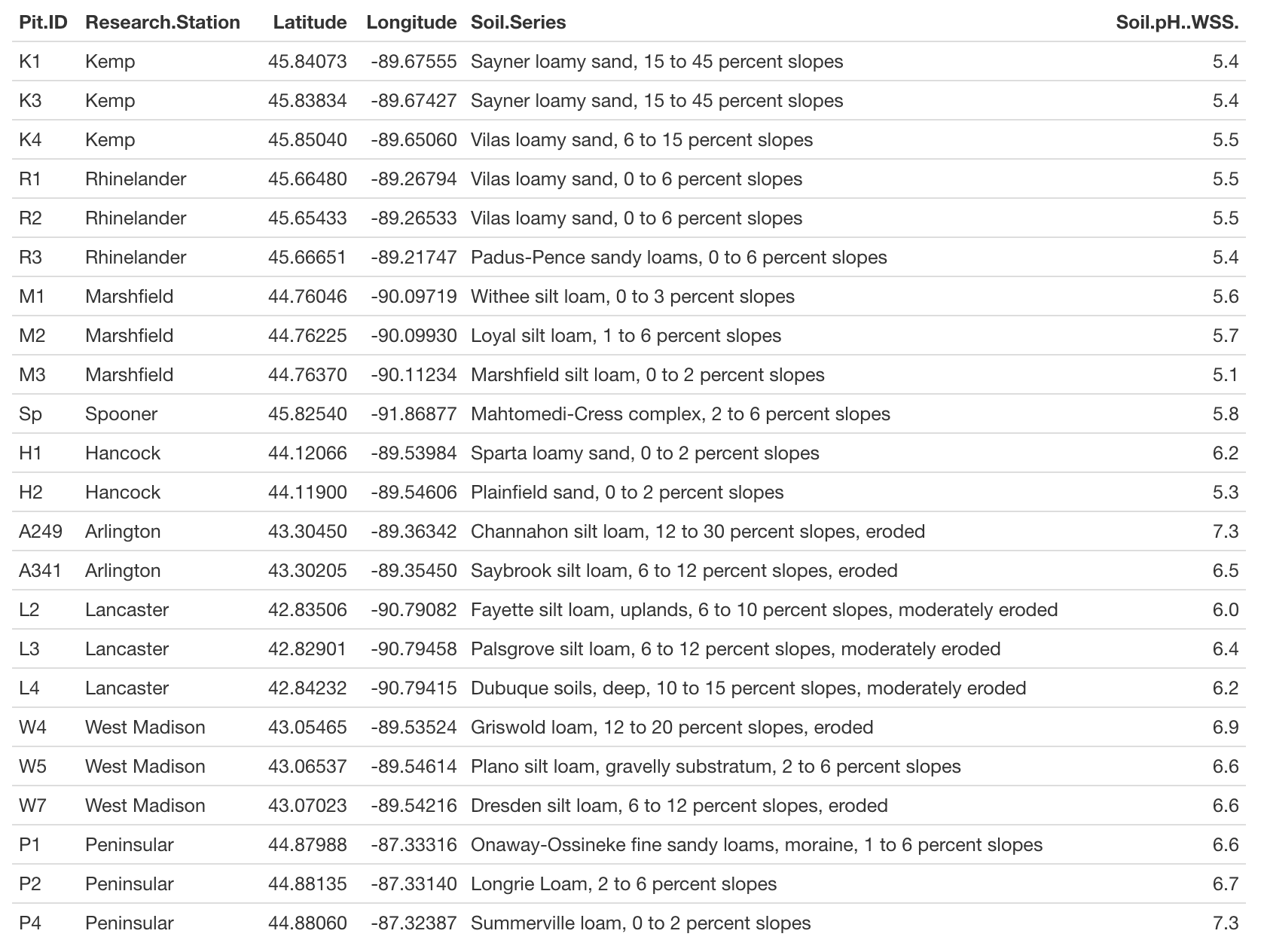

### uw-experimental-farms-major-soils.jpg

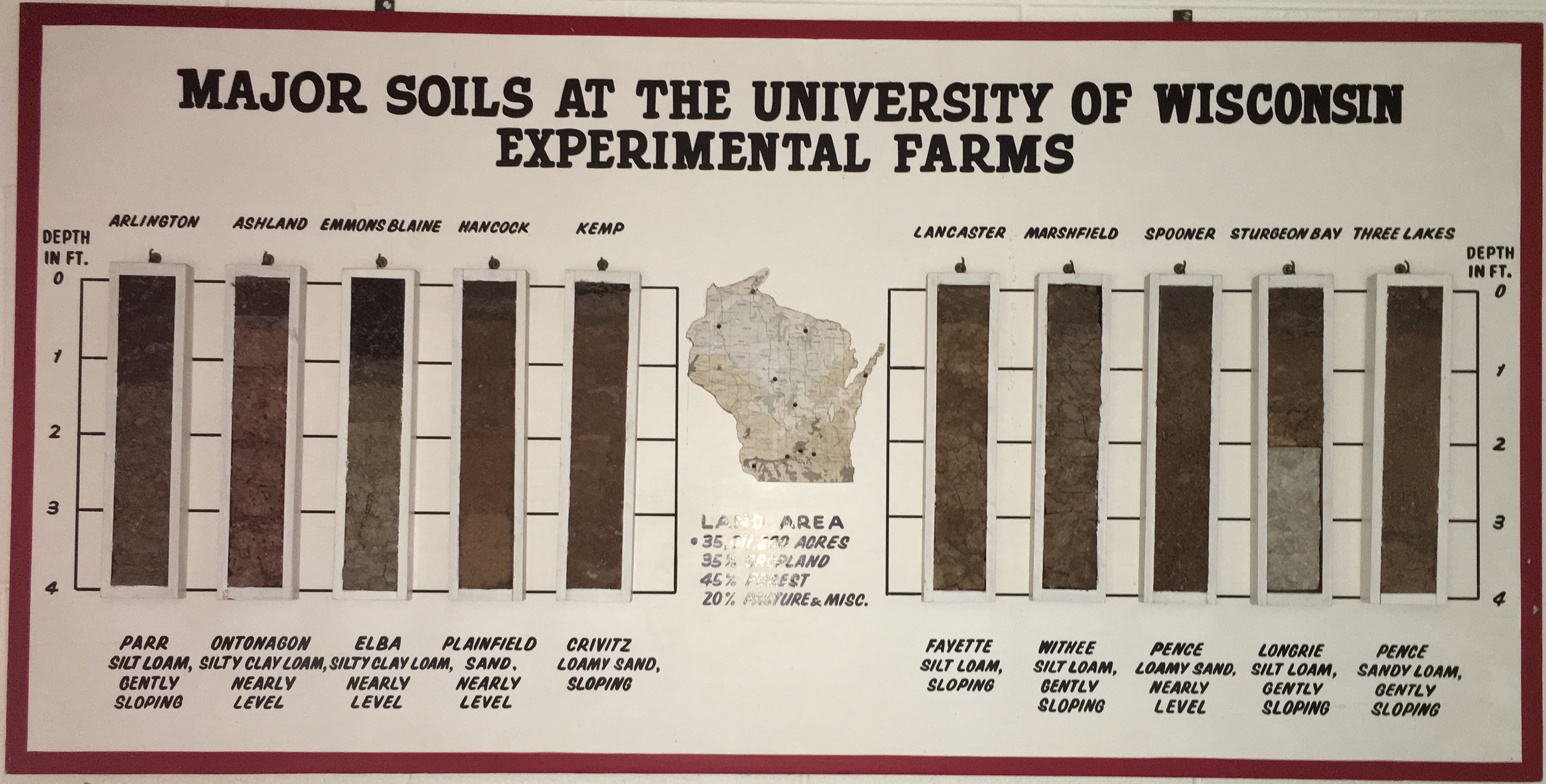

### w4-0cm-60cm.jpg

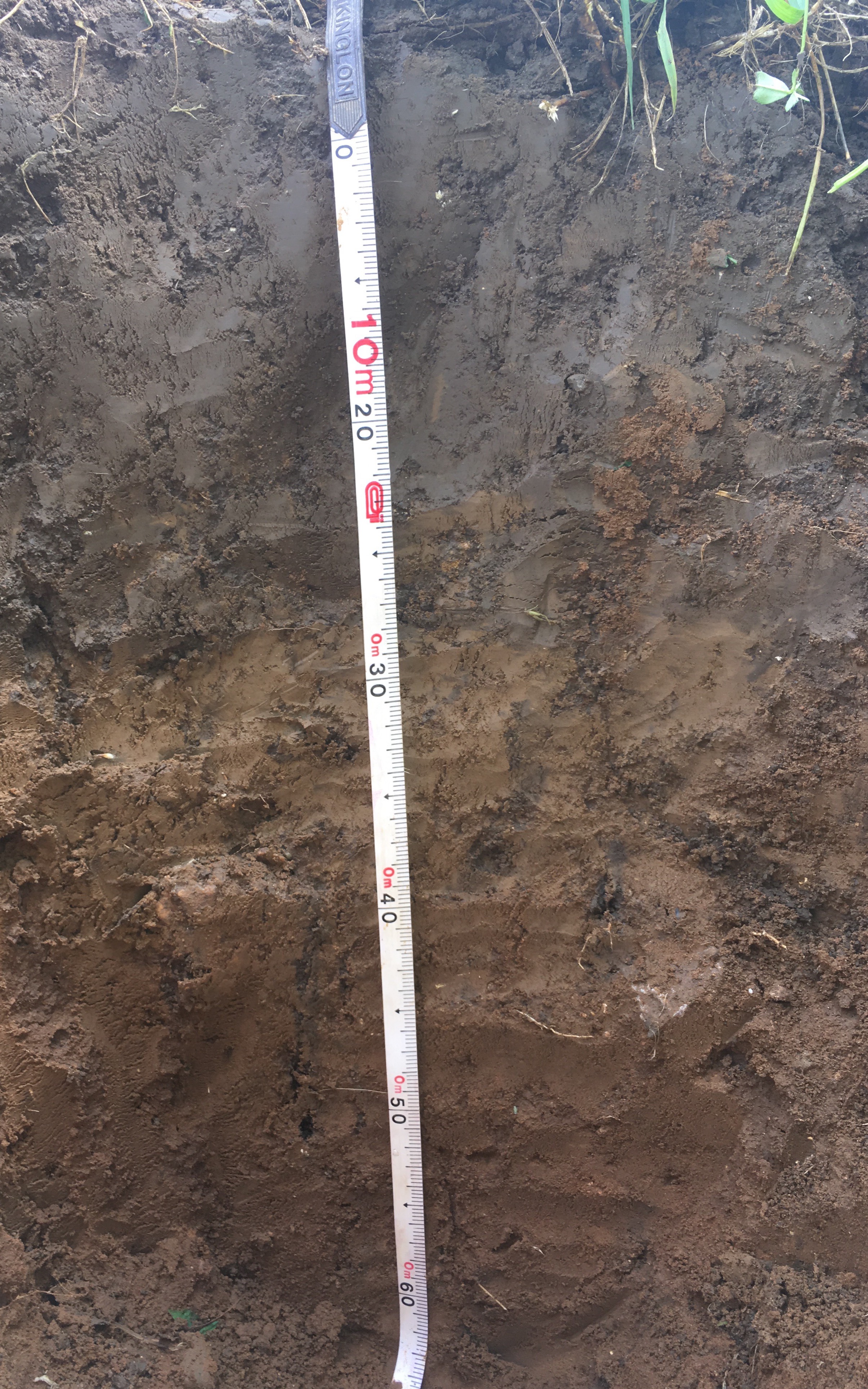

### w5-0cm-60cm.jpg

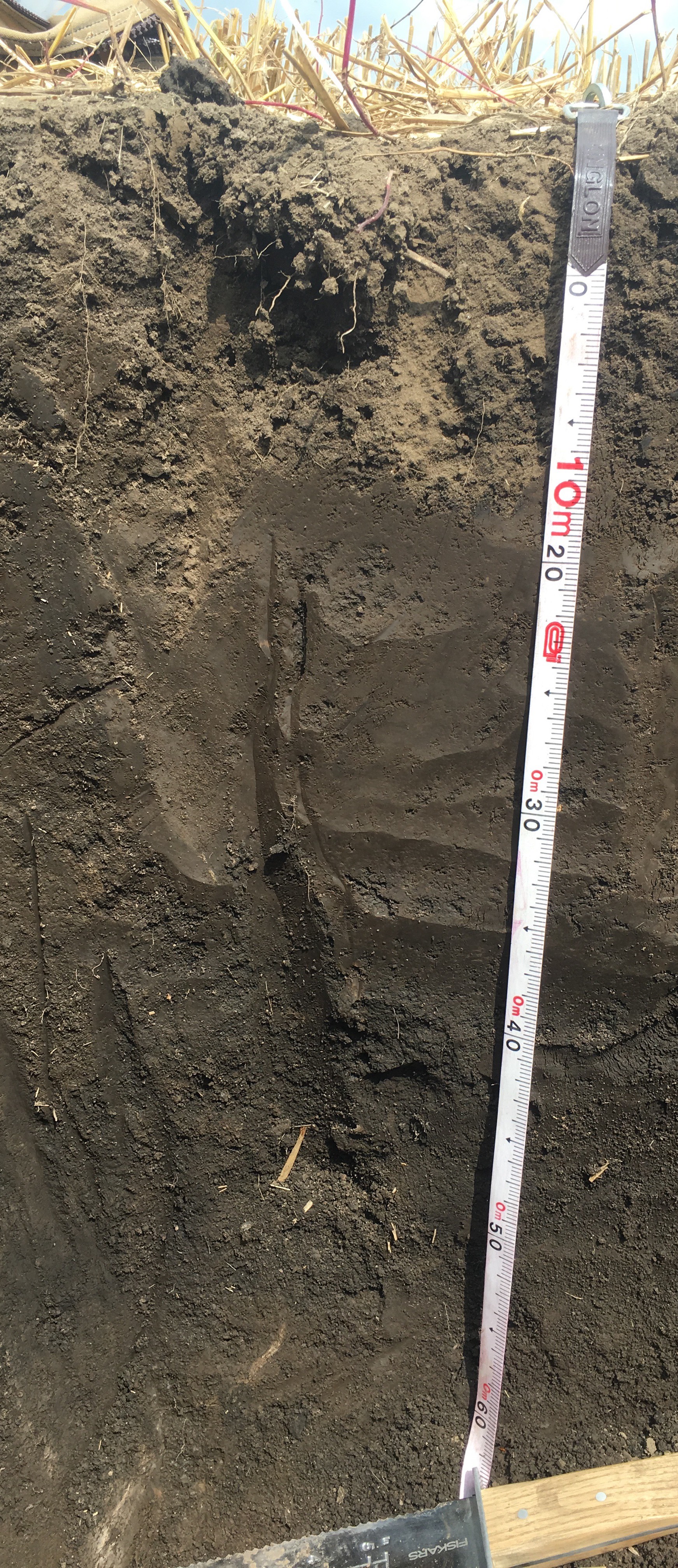
